# Class A PBPs reinforce the septal cell wall following initial synthesis by SEDS–bPBP pairs during bacterial cytokinesis

**DOI:** 10.1101/2025.09.12.675754

**Authors:** Jiajia Wang, Jie Hu, Yue Xin, Lihao Ding, Xinci Wang, Xianghui Ou, Junyi Pang, Yongfeng Liu, Sijing Zhang, Liang Zhang, Hui Wang, Chun-ying Yin, Yongxiang Gao, Ting Xue, Yandong Yin, Jiangliu Yu, Lin Xue, Xinxing Yang

## Abstract

Bacterial cytokinesis requires the coordinated action of septal peptidoglycan (PG) synthases to ensure efficient and accurate septum formation. Two major types of synthases contribute: SEDS–bPBP and bifunctional class A PBPs (aPBPs). How their activities are organized in space and time has remained unclear. Using *Deinococcus radiodurans*, which often contains both constricting and closed septa in the same cell, we show that septal PG synthesis proceeds through distinct steps at different regions. Primary synthesis at the leading edge by SEDS–bPBP pairs drives constriction, whereas aPBPs act later at the lagging edge and in closed septa to thicken the wall, enhance cross-linking, and reinforce septal integrity. At the single-molecule level, both synthase types move directionally to perform processive PG synthesis at distinct sites. We also observed spatiotemporal separation of SEDS–bPBP and aPBP synthases in *Staphylococcus aureus*, suggesting that secondary septal PG synthesis is a conserved mechanism securing successful cytokinesis.

## Introduction

Bacterial cytokinesis demands a complete rebuild of the cell wall at the division site, where septal peptidoglycan (PG) synthesis must be executed with exquisite spatial and temporal precision^1,2^. The septal PG not only drives membrane invagination but also dictates the shape and integrity of daughter cell poles^3,4^. To orchestrate this process, bacteria assemble the divisome^5,6^, a dynamic protein complex that integrates spatial and temporal cues to control PG synthase activity. At its core is the tubulin-like cytoskeletal protein FtsZ, which polymerizes into a ringlike structure (Z-ring). The Z-ring recruits a network of scaffold and regulatory proteins, ultimately guiding the accumulation of PG synthases that construct the septal wall^7–9^.

PG is assembled from the lipid II precursor, which is flipped into the periplasm and incorporated into the existing PG meshwork by glycosyltransferases (GTases) and transpeptidases (TPases). Two major synthase systems mediate this process. The bifunctional class A PBPs (aPBPs) carry both GTase and TPase activities^2,10^, while members of the SEDS (Shape, Elongation, Division, Sporulation) family, such as RodA and FtsW, function as GTases with cognate monofunctional class B PBPs (bPBPs) that provide TPase activity **(Figure 1A)**^11,12^. These SEDS–bPBP pairs are genetically essential in most bacteria and act in parallel with aPBPs^13^, but their precise coordination during septal assembly remains unresolved.

**Fig. 1.**
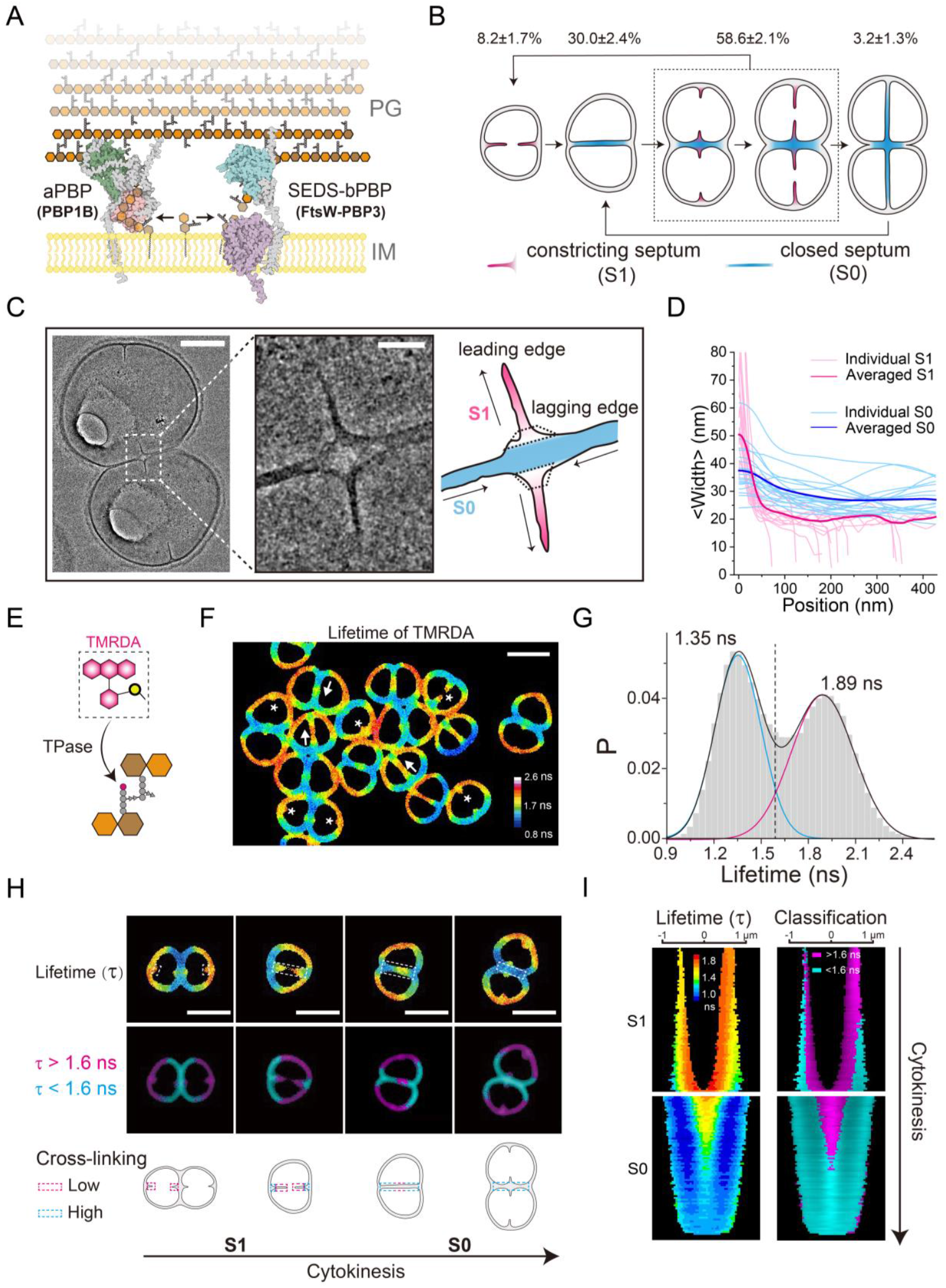
Constricting septa (S1) transition into thickened, highly cross-linked closed septa (S0) during cytokinesis in *D. radiodurans*. **(A)** Schematic of septal PG synthesis by SEDS–bPBP pairs (FtsW–PBP3 in *D. radiodurans*) and the bifunctional aPBPs (PBP1B in *D. radiodurans*). FtsW acts as a glycosyltransferase (GTase), PBP3 as a transpeptidase (TPase), and PBP1B performs both activities. PG, peptidoglycan; IM, inner membrane. **(B)** Cell cycle schematic highlighting five morphological stages. Constricting septa (S1, magenta) and closed septa (S0, cyan) are indicated with population frequencies from exponential-phase cultures. **(C)** TEM images showing a representative diad with both S1 and S0 (left, scale bar 500 nm), a magnified region where S1 emerges from S0 (middle, 100 nm), and a schematic illustrating the leading edge (magenta), wedge-like lagging edge (white, dashed trapezoid), and thickened S0 (cyan). Arrows mark the direction of septum constriction. **(D)** Width profiles of individual and averaged S1 (magenta) and S0 (cyan) along the direction of septum constriction in (C). **(E)** Schematic of TAMRA–Damino acid (TMRDA) incorporation by transpeptidases. **(F)** Fluorescence lifetime image of TMRDA–labeled cells. Asterisks mark nascent S1 with long lifetimes; arrows mark lifetime increases toward the center of S0 septa. Scale bar, 2 μm. **(G)** Bimodal fluorescence lifetime distribution at ~1.35 ns (cyan) and ~1.89 ns (magenta). The dashed line (1.6 ns) was used to classify the two populations. **(H)** Representative cells at successive septal formation stages. Top: lifetime images. Middle: binarized classification (cyan < 1.6 ns; magenta > 1.6 ns). Bottom: schematics of cross-linking at S1 and S0. Scale bars, 2 μm. **(I)** Demographs of septal PG lifetimes (left) and binarized maps (right) throughout the cytokinesis (Methods; Figure S2D).

In many bacteria, SEDS–bPBP pairs move directionally at the septum, driving processive PG synthesis with variable dependence on FtsZ treadmilling^14–19^. In contrast, aPBPs were long thought to play a role in maintaining global PG integrity, repairing damage from autolysins or stress^20–23^. Consistent with this view, aPBPs generally display broad membrane distribution and diffusive dynamics^13,24–26^. Yet disruption of aPBPs activity in several species causes division defects, and genetic studies have revealed their interactions with core divisome proteins^27–30^. In *Escherichia coli* (*E. coli*) and *Staphylococcus aureus* (*S. aureus*), aPBPs have also been implicated in pre-septal (PBP3-independent) PG synthesis^31–34^. These observations suggest that aPBPs do contribute to septum formation, but their spatiotemporal and functional relationship to SEDS–bPBP is still poorly defined. Resolving enzyme activity and spatiotemporal organization is particularly difficult in the small, rapidly constricting septa of model bacteria.

To better resolve enzyme activity and septal architecture, we turned to *Deinococcus radiodurans* (*D. radiodurans*). This large, coccoid bacterium (~1.5 µm in diameter) provides better spatial resolution under light microscopy than smaller rods and cocci such as *E. coli* and *S. aureus*. Moreover, more than half of *D. radiodurans* exist as diads that simultaneously harbor a closed septum from the previous division and nascent constricting septa **(Figure 1B)**^35^. This special cell cycle extends the temporal window for observing cytokinesis, allowing us to compare distinct stages of septal formation within the same cell. *D. radiodurans* also features a Gram-positive–like thick PG cell wall together with an outer membrane, combining traits of both major bacterial lineages^36^. Although *D. radiodurans* is best known for its resistance to DNA damage^37–39^, investigations into its cell envelope biogenesis and cell division have only recently begun to emerge^40,41^. These attributes make *D. radiodurans* an attractive system for resolving septal architecture and enzyme organization potentially deepen our understanding of bacteria cytokinesis.

In this study, we employed fluorescent PG probes combined with fluorescence lifetime imaging microscopy (FLIM) to measure local cross-linking of septal PG, together with electron tomography to revealed a two-step architecture development of the septum during cytokinesis. We further resolved the distinct spatial distributions and dynamics of core PG synthases in *D. radiodurans*. Our results show that septum formation proceeds through two spatially separated steps: SEDS–bPBP pairs mediate primary septal synthesis at the leading edge to drive septum constriction, while aPBPs act later at the lagging edge and in closed septa to thicken the wall, increase cross-linking, and reinforce septal integrity. Extending this study to *S. aureus* demonstrated a similar spatiotemporal separation of SEDS–bPBP and aPBP synthases, suggesting that bacteria likely rely on a conserved mechanism in which distinct synthases act sequentially to complete septum formation during cytokinesis.

## Results

### Constricting (S1) and closed (S0) septa coexist in D. radiodurans

To investigate the morphological changes of the septum during cytokinesis, we grouped *D. radiodurans* cells into five stages **(Figure 1B; Methods)**, following criteria adapted from Floc’h et al^35^. Over half of the cells (58.6 ± 2.1%, mean ± s.e.m.) formed diads that displayed both two nascent constricting septa **(hereafter S1, marked in pink in Figure 1B)** and a closed septum from the previous division **(S0, cyan in Figure 1B)**. Transmission electron microscopy (TEM) revealed that S1 emerged from the center of S0 and displayed a thinner tip **(Figure 1C)**, consistent with earlier cryo-electron tomography analyses^41,42^. Behind the thin leading edge of S1, we observed a wedge-like density (~50 nm along the constricting direction) at the base of the septum, hereafter referred to as the lagging edge. The progressive increase in septal thickness from the leading edge of S1, through its lagging edge and ultimately to the mature S0 suggests sequential incorporation of new materials in septal PG during cytokinesis in *D. radiodurans* **(Figure 1D)**.

### High cross-linking of septal PG at the lagging edge of S1 and in S0

The thickened architecture of the lagging edge of S1 and the closed septum (S0) raised the question of whether the chemistry of septal PG also changes during cytokinesis. We labeled *D. radiodurans* with a fluorescent D-amino acid probe TMRDA (tetramethylrhodamine– conjugated D-amino acid^43^). Since the fluorescent D-amino acids are incorporated into PG by transpeptidases that also catalyze PG crosslinking **(Figure 1E)**^44,45^, probe density reflects TPase activity and the extent of cross-linking. To assese the probe density, we monitored probe incorporation using FLIM. Unlike fluorescence intensity, fluorescence lifetime is largely insensitive to imaging conditions but reports on the chemical environment, including probe density **(Figure S1A)**. High probe concentration shorten fluorescence lifetimes and causes intensity first increase and then decrease, indicating self-quenching effect from nearby probes **(Figure S1B, C)**. Additionally, dual-labeling with two spectrally overlapping probes—RhoDA (donor) and TMRDA (acceptor) resulted in Förster Resonance Energy Transfer (FRET) between the probes, confirming close probe spacing (<6 nm) at high acceptor concentrations **(Figure S1D–F)**.

After a two-hour labeling (approximately one generation time), TMRDA clearly delineated both peripheral and septal PG and revealed a bimodal lifetime distribution, reflecting distinct PG cross-linking states **(Figure 1F, G)**. The leading edge of the S1 exhibited relatively longer lifetime than the lagging edge **(asterisks in Figure 1F)**. In S0, lifetimes were generally shorter than in S1 yet increased progressively from the side near sidewall toward the center **(arrows in Figure 1F)**. Representative cells at successive stages of cytokinesis illustrate this trend **(Figure 1H)**. The leading edge of S1 first formed with long-lifetime signals (τ > 1.6 ns), indicating low cross-linking. A short-lifetime population (τ < 1.6 ns) then emerged at its lagging edge, gradually expanding during septum closure and ultimately covering the entire closed septum. This pattern reveals a secondary PG synthesis step that increases cross-linking. Demograph analysis of the septum formation confirmed that the highly cross-linked population originated at the lagging edge near the sidewall and followed behind the low-cross-linked leading edge, eventually occupying the full septum **(Figure 1I; Figure S2; Methods)**. Consistently, high FRET efficiency was detected at the lagging edge of S1 in the dual-label experiment, corroborating that this region undergoes additional cross-linking through a secondary PG synthesis step **(Figure S1G, H)**.

### FtsW-PBP3, the SEDS-bPBP in D. radiodurans, localizes to the leading edge of S1

The distinct architecture and cross-linking patterns at the leading and lagging edges of S1 prompted us to ask whether septal PG synthases occupy specific regions within the septum. Two types of SEDS–bPBP pairs—RodA–bPBP and FtsW–bPBP—function as canonical PG synthases in diverse bacteria, driving sidewall elongation and septum synthesis, respectively^11,13,14,46,47^. In *D. radiodurans*, we identified a single class B PBP–encoding gene (***dr_1868***) and a single SEDS-family gene (***dr_2497***), encoding proteins with predicted TPase and GTase domains, respectively **(Figure S3A, B)**. Given that *D. radiodurans* lacks a RodA homolog and displays a coccoid morphology^48^, we reason that the proteins encoded by ***dr_1868*** and ***dr_2497***, renamed as PBP3 and FtsW, together constitute the dedicated septal PG synthase pair in this organism.

To investigate their subcellular localization, we constructed strains expressing C-terminal FtsW-mNeonGreen^49^ (FtsW–mNG) and N-terminal mNG–PBP3 **(Table S1, 2; Figure S4A, B)**. Co-staining with the membrane dye Potomac Red^50^ revealed that both FtsW and PBP3 were enriched at the leading edge of S1, closely resembling the distribution of the Z-ring formed the core division protein FtsZ, which assemble into a Z-ring^51,52^ **(arrows in Figure 2A; Table S3)**. Three-color imaging confirmed that FtsW and PBP3 colocalized with FtsZ to a greater extent than with the membrane **(Figure S4C; Figure S5A, B; Table S1-3)**, supporting that FtsW and PBP3 constitute the divisionspecific SEDS-bPBP pair for septal PG synthesis.

**Fig. 2.**
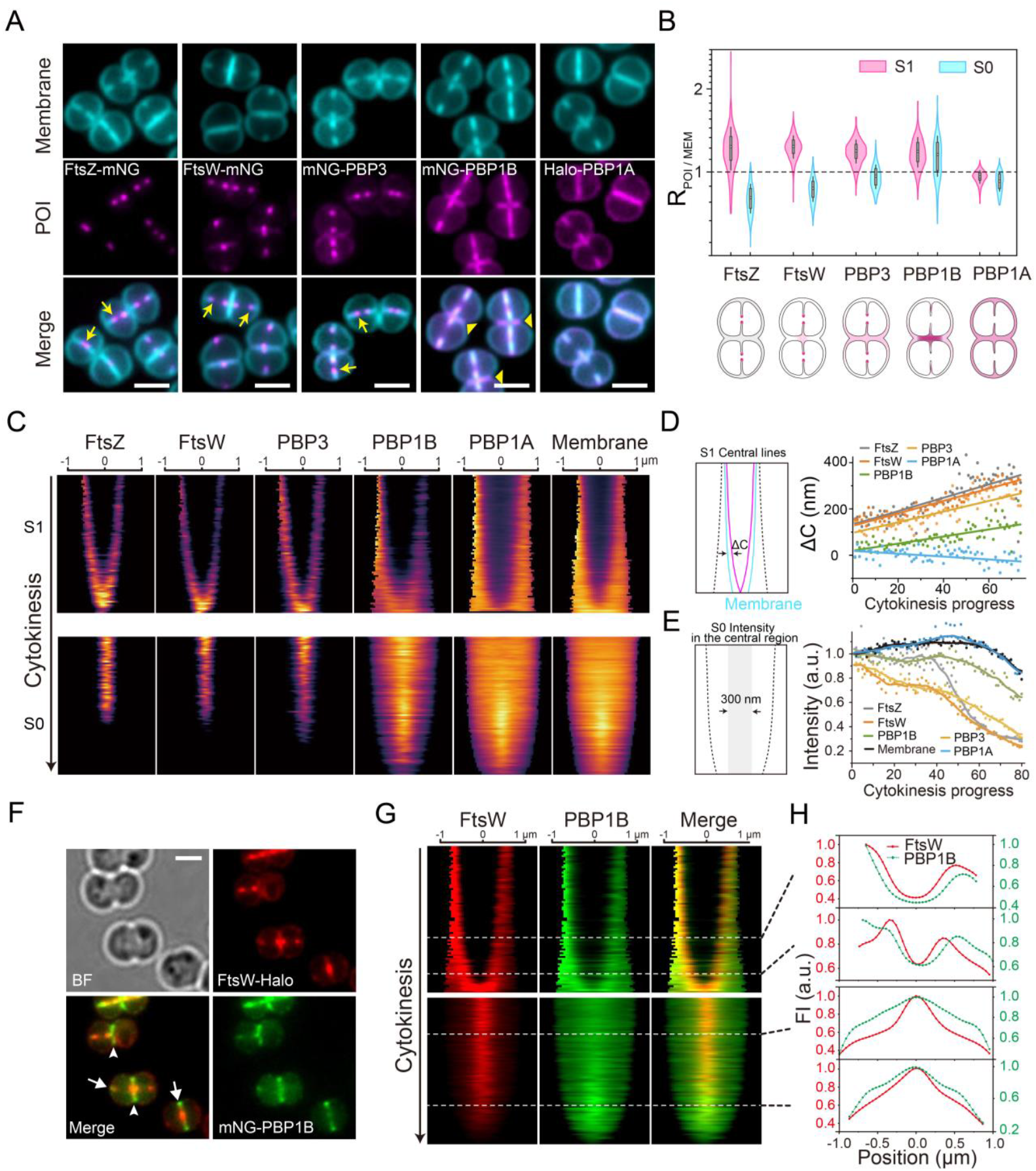
FtsW–PBP3 localizes to the leading edge of constricting septa, whereas PBP1B accumulates at the lagging edge and in closed septa. **(A)** Representative fluorescence images showing localization of FtsZ, FtsW, PBP3, PBP1B, and PBP1A (magenta) relative to membrane staining (cyan). Arrows, FtsZ/FtsW/PBP3 enrichment at the leading edge of S1; arrowheads, PBP1B accumulation at S0. Scale bars, 2 μm. **(B)** Quantification of septal enrichment. Top: the fraction of each protein at S1 or S0 normalized to the membrane signal (RPOI/MEM); RPOI/MEM > 1 indicates enrichment. Bottom: schematics summarizing localization patterns. Violin plots show data distributions; boxes mark the 25–75% range with SD whiskers, and medians are indicated by squares. MEM: Membrnae. **(C)** Demographs of protein and membrane signals at S1 (top) and S0 (bottom) throughout the cytokinesis. **(D)** Average displacement (ΔC) between protein and membrane signals at S1, derived from demographs. **(E)** Average fluorescence intensity within the central 300 nm of S0 septa. **(F)** Co-localization of FtsW–Halo and mNG–PBP1B. Arrows, PBP1B positioned flanking FtsW at S1; arrowheads, PBP1B accumulation at S0. Scale bar, 1 μm. **(G)** Demographs of FtsW (red) and PBP1B (green) across cytokinesis. Merged patterns resemble cross-linking transitions shown in Figure 1I. **(H)** Line profiles of FtsW (red) and PBP1B (green) across S1 and S0 at different constriction stages (from (F)). PBP1B consistently lags behind FtsW at S1 and persists at S0 after FtsW disassembly.

We next quantified the relative enrichment of FtsZ, FtsW, and PBP3 at S1 and S0 by normalizing their septal fluorescence signals to that of the membrane **(Figure 2B, Methods; Figure S4E)**. Both PBP3 and FtsW showed enrichment ratios (R) greater than 1 at S1, demonstrating preferential localization at S1, likely through recruitment by FtsZ^53^. By contrast, their R values at S0 were < 1, reflecting rapid disassembly of FtsW and PBP3 after septum closure. Demograph analyses showed that FtsW and PBP3 accumulated at the leading edge of S1, tracking closely with the Z-ring **(Figure 2C, D)**. Together, these results indicate that FtsW and PBP3 colocalize with FtsZ at the leading edge of S1, where they likely mediate the primary septal PG synthesis, but disassembled after septum closure.

### PBP1B, the class A PBP type synthase, localizes at the lagging edge of S1 and in S0

The high cross-linking detected at the lagging edge of S1 and in S0 raised the possibility that a different class of PG synthases operates at these sites. In addition to FtsW–PBP3 pairs, class A PBPs have been implicated as septal PG synthases in other bacteria^1,2,5,10,25,27,28,30,54^, prompting us to examine their role in *D. radiodurans. D. radiodurans* encodes two class A PBP homologs (*dr_0479* and *dr_1417*), which we designate PBP1A and PBP1B, respectively **(Figure S3C, D)**. To assess their subcellular localizations, we generated functional HaloTag^55^–PBP1A (Halo-PBP1A) and mNG–PBP1B fusions and analyzed their localization alongside FtsW and PBP3 **(Figure 2A; Figure S4A; Tabel S1–3)**. PBP1A displayed an even distribution around the cell envelope, indistinguishable from membrane staining, and showed no enrichment at either S1 or S0 **(Figure 2A–D; Figure S5A, B)**, indicating it is not division-specific. By contrast, PBP1B accumulated at constricting septa but, unlike FtsW or PBP3, did not form sharp foci at the leading edge. Demograph analyses showed a broad distribution of PBP1B centered toward the lagging edge, approximately 160 nm behind the FtsZ-marked leading edge **(Figure 2C, top; Figure 2D)**. Notably, PBP1B also displayed strong enrichment at the closed septum (S0) (arrowheads in Figure 2A), as reflected by enrichment ratios (R) >1 **(Figure 2B)** and a slowed decay of signal intensity across the septal midline in S0 **(Figure 2C, D)**.

Direct comparison with FtsW revealed that PBP1B either flanked FtsW foci at S1 or accumulated at S0 **(Figure 2F; Figure S4D; Table S1, 3)**. Colocalization analysis further revealed that PBP1B exhibited weaker overlap with FtsZ or PBP3 compared to the strong associations observed among FtsW–PBP3, FtsW–FtsZ, and PBP3–FtsZ. Quantification by Pearson correlation coefficient (PCC) analysis confirmed the reduced colocalization of PBP1B with either FtsZ or PBP3 **(Figure S5)**. Two-color demographs showed that PBP1B consistently localized behind FtsW along the constricting septum S1 and remained at S0 after FtsW had departed **(Figure 2G, H)**. This parallels the emergence of highly cross-linked PG at the lagging edge of S1 and in S0 **(Figure 1I)**.

These observations suggest that PBP1B, unlike PBP1A, functions as the secondary septal PG synthase at the lagging edge of S1 and in S0.

### Inhibition of PBP3 blocks the septum constriction, whereas inhibition of PBP1B prevents thickening of the lagging edge of S1 and S0

To directly test whether the spatiotemporal separated FtsW–PBP3 and PBP1B mediate septal PG synthesis at the leading versus lagging edge of S1 (and S0), we examined septal architecture under selective inhibition of their enzymatic activities. We first verified that two β-lactams— aztreonam and cefsulodin—specifically inhibit the TPase activity of PBP3 and PBP1B, respectively **(Figure S6A–C; Methods)**^56^. In aztreonam-treated *D. radiodurans* cells, three-dimensional singlemolecule localization microscopy (3D-SMLM) revealed a failure to complete septal closure, resulting in very few fully closed septa (S0) **(Figure 3A; Figure S7B; Methods)**. Live-cell imaging also confirmed blocked constriction and failed cell division **(Figure 3F; Movie S1, 2)**. Moreover, aztreonam-treated cells displayed blurry, broadened septa at the lagging edge of S1 **(arrowheads in Figure 3A)**, suggesting continued PG incorporation by other synthases but loss of primary septal synthesis at the leading edge. By contrast, cefsulodin treatment did not impair septum constriction or closure **(Figure 3A, F; Figure S7C; Movie S3)** yet produced a distinct phenotype of dim and distorted S0 septa **(arrows in Figure 3A)**. Additionally, inhibition of the GTase activity of PBP1A and PBP1B with moenomycin^57^ (No MtgA-like GTase has been identified in the genome of *D. radiodurans*.) caused similar defects **(Figure 3A, F; Movie S4)**. These distorted and likely weakened septa indicate that PBP1B is required to reinforce the septal structure primarily laid down by FtsW–PBP3 pairs.

**Fig. 3.**
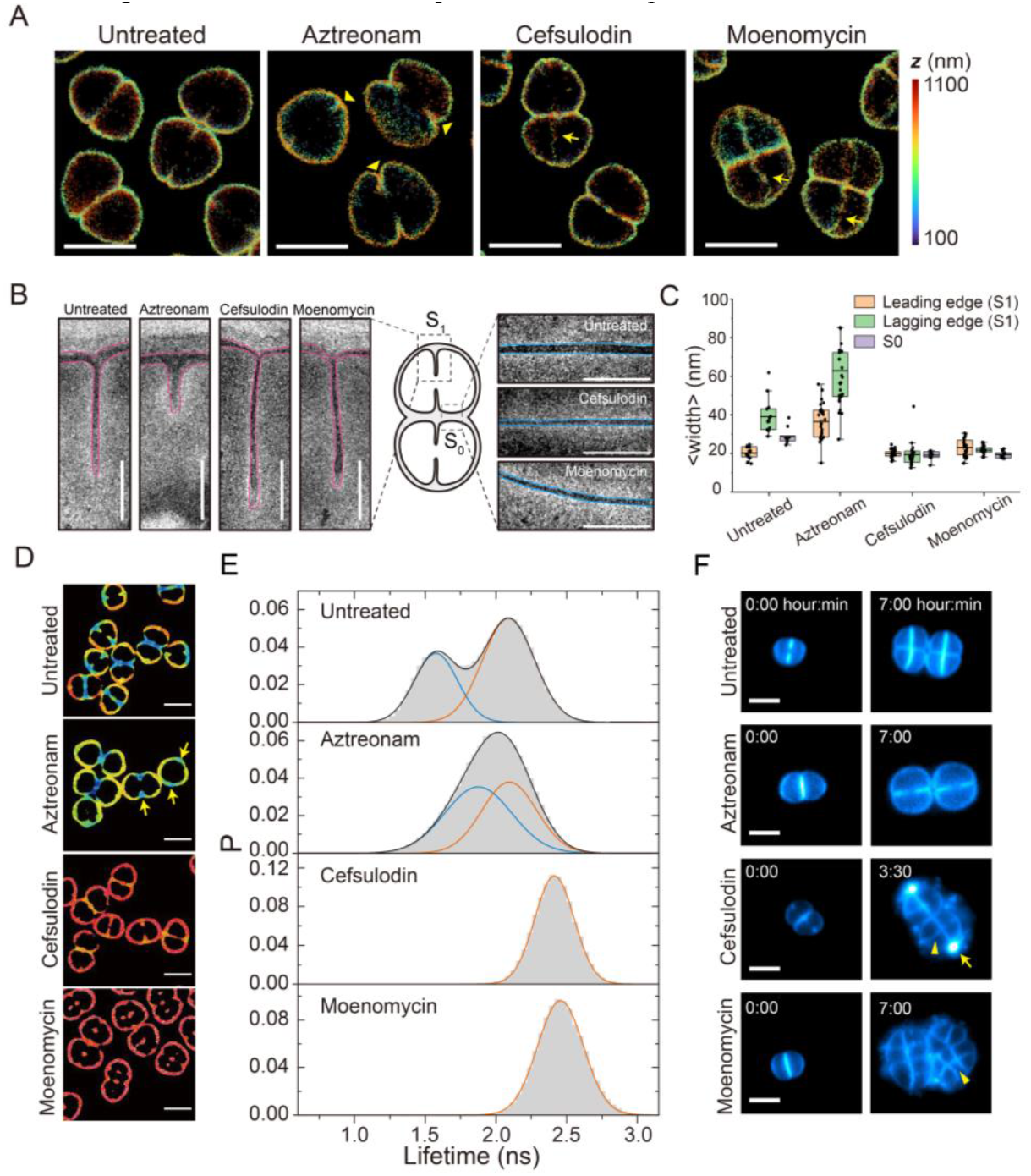
Secondary septal PG synthesis by PBP1B increases cross-linking and reinforces septal integrity. **(A)** 3D-SMLM images of Cy5DA–labeled *D. radiodurans* cells untreated or treated with aztreonam (PBP3 TPase inhibitor), cefsulodin (PBP1B TPase inhibitor), or moenomycin (PBP1B/A GTase inhibitor). Arrows, broadened constricting septa (S1); arrowheads, dim and distorted closed septa (S0). Scale bars, 2 μm. **(B)** Room-temperature electron tomography of S1 (left) and S0 (right) septa under the same conditions as in (A). PG outlines are traced for width measurements in (C). Scale bars, 100 nm. **(C)** Quantification of septal width of the leading edge, lagging edge of S1 and S0. The lagging edge of S1 is defined as the first 50 nm from the sidewall. Box plots show mean (lines), interquartile ranges (boxes), and 1.5× IQR (whiskers). **(D)** Representative FLIM images of TMRDA–labeled cells, untreated or after aztreonam, cefsulodin, or moenomycin treatment. Arrows, short-lifetime regions at the lagging edge of S1 septa. Scale bars, 2 μm. **(E)** Lifetime distributions: bimodal in aztreonam-treated cells, monomodal at long lifetimes in cefsulodinand moenomycin-treated cells. **(F)** Time-lapse images of cells labeled with Potomac Gold dye, untreated or after drug treatment. Aztreonam blocked septal closure; cefsulodin and moenomycin caused membrane rupture at the lagging edge (arrows) and produced dim, distorted septa unable to complete division (arrowheads). Scale bars, 2 μm.

Because the ~20–40 nm spatial resolution of 3D-SMLM was insufficient to resolve septal ultrastructure **(Figure S7E)**, we turned to electron tomography **(Figure 3B)**. In wild-type cells, the constricting septa (S1) exhibited straight, thin tips (20.1 ± 3.0 nm, mean ± s.d.; Table S4) that extended inward, followed by a wedge-like region, while closed septa (S0) were thicker on average (29.0 ± 4.5 nm) **(Figure 3B, C; Figure S8A; Methods; Table S4)**. Upon aztreonam treatment, S1 thickness increased to 47.5 ± 15.3 nm, accompanied by exaggerated wedge-like bulges **(Figure S8B; Table S4)**, confirming continued synthesis was blocked. In contrast, cefsulodin or moenomycin treatment left the leading edge of S1 unaffected but abolished the wedge-like structure: septa lacking PBP1B activity displayed comparable widths at the lagging and leading edges **(Figure S8C, D; Table S4)**. These cells also lacked the normal thickening of S0 and instead often showed distorted septa **(Figure S8E–G; Table S4)**. These perturbations show that FtsW–PBP3 pairs drive primary synthesis at the leading edge to promote constriction, while PBP1B mediates a secondary synthesis step that thickens the septum from the lagging edge and fortifies the closed septum.

### Secondary septal PG synthesis by PBP1B increases the cross-linking and reinforces the septum integrity

We next asked whether the high cross-linking at the lagging edge of S1 and in S0 **(Figure 1E–H)** are generated by PBP1B. Cells treated with aztreonam, cefsulodin, or moenomycin were subsequently labeled with TMRDA for FLIM analysis. Aztreonam-treated cells formed many incomplete septa but still exhibited short lifetimes at the lagging edge **(arrows in Figure 3D)**, indicating continued cross-linking by other synthases. The overall lifetime distribution shifted toward longer values, due to the loss of most closed septa (S0), which normally contribute the highest cross-linking **(Figure 1H, I)**. In contrast, inhibition of PBP1B’s TPase activity with cefsulodin eliminated the short-lifetime population, even though septa could still constrict and close **(Figure 3D, E; Movie S3)**. These less cross-linked and thinner septa frequently ruptured during division **(arrowhead in Figure 3F)**, leading to membrane disruption and leakage of cytoplasmic contents **(arrow in Figure 3F)**. Similarly, inhibition of the GTase activity with moenomycin lowered cross-linking and destabilized the septal wall **(Figure 3D–F; Movie S4)**, indicating that the TPase and GTase activities of PBP1B act in concert.

To exclude contributions from other transpeptidases, we deleted *pbp1A* (encoding PBP1A) **(Figure S9A)**. The Δ*pbp1A* strain showed no growth or morphological defects relative to wild-type *D. radiodurans* (DRWT) **(Figure S9C)** and exhibited comparable cross-linking levels at the lagging edge of S1 and in S0 **(Figure S9D, E)**, consistent with its uniform spatial distribution **(Figure 2A, B)**. Thus, PBP1A is a nonessential PG synthase that plays little, if any, role in septal synthesis. We also deleted *pbp4* (*dr_0176*), encoding a class C PBP predicted as a D-Ala–D-Ala carboxypeptidase **(Figure S3E)**. The Δ*pbp4* strain exhibited no major changes in growth or cell size **(Figure S9B, C)**, but showed even shorter lifetimes at the lagging edge and in S0 **(Figure S9D, E)**. This is consistent with loss of probe removal from pentapeptide stems, leading to increased TMRDA density and lifetime shortening. Finally, we found that the genome of *D. radiodurans* contains no predicted homologs of L,D-transpeptidases. Together, these results demonstrate that secondary septal PG synthesis reinforces the septum is performed mainly by the class A PBP, PBP1B.

### Directional movements of FtsW and PBP1B at different septal sites drive primary and secondary PG synthesis

To dissect the molecular mechanisms of septal PG synthesis, we performed single-molecule tracking (SMT) of FtsW and PBP1B in live *D. radiodurans* cells. Consistent with its subcellular localization, most FtsW trajectories were detected at the leading edge of S1 **(86.0 ± 3.7%, mean ± s.d.; Figure 4A)**. Most FtsW molecules exhibited two characteristic behaviors: directional movement and confined diffusion **(Figure 4B; Figure S10A)**, as reported for homologs in *E. coli*^14^, *Caulobacter crescentus* (*C. crescentus*)^46^, *Bacillus subtilis* (*B. subtilis*)^17^, *S. aureu*s^18^, and *Streptococcus pneumoniae* (*S. pneumoniae*)^58^. Directionally moving FtsW molecules displayed a bimodal speed distribution, with fast (15.5 ± 1.4% nm/s, 81.9%) and slow (2.7 ± 3.6% nm/s, 18.1%) populations **(Figure 4F; Movie S5-7)**.

**Fig. 4.**
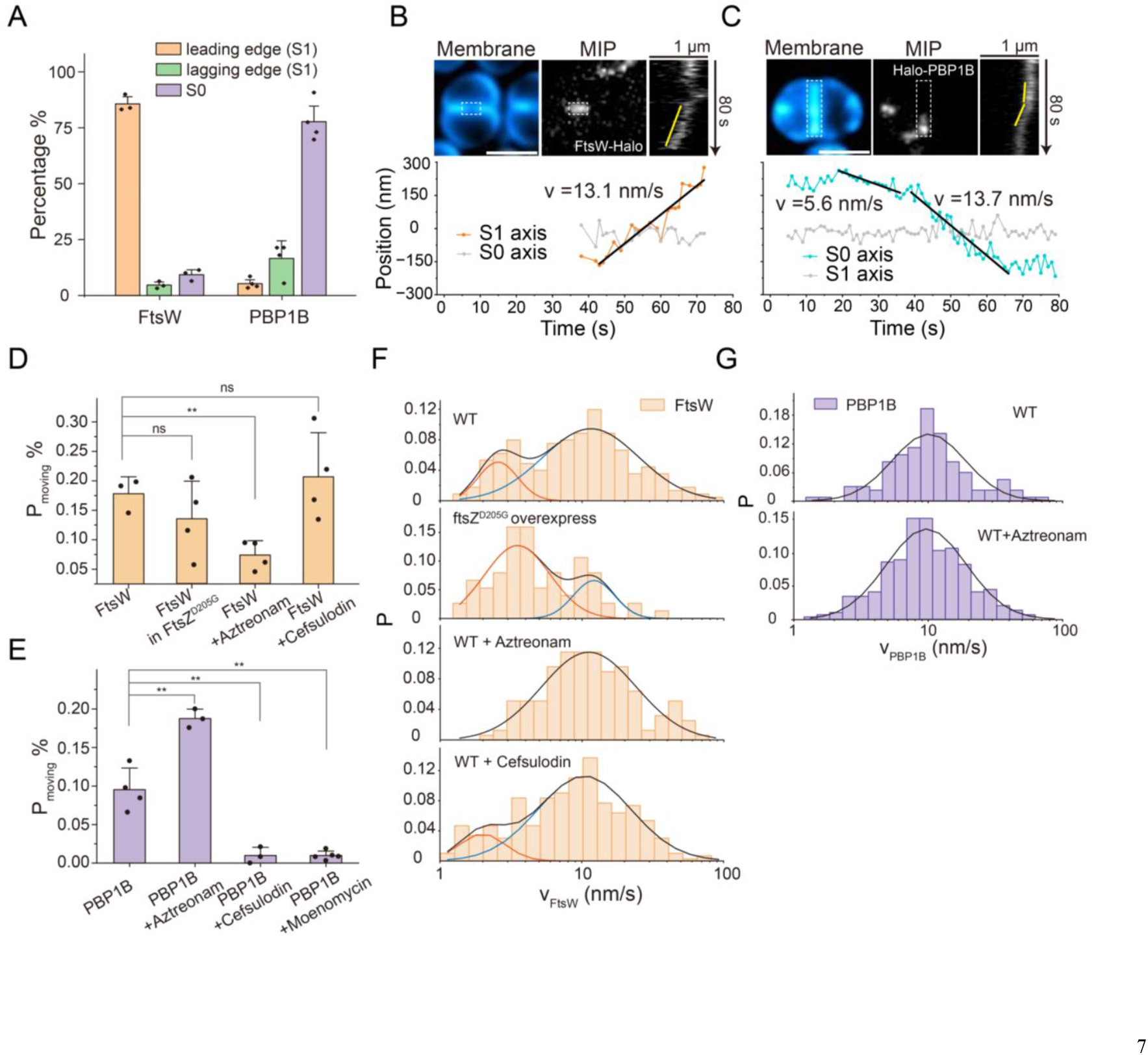
FtsW processively synthesizes PG at the leading edge of S1, whereas PBP1B acts at the lagging edge of S1 and in closed septa, revealed by single-molecule tracking. **(A)** Spatial distribution of single-molecule trajectories of FtsW and PBP1B at division sites. FtsW molecules predominantly localizes to the leading edge of S1, whereas PBP1B molecules concentrates at S0. **(B)** Representative trajectory of a directionally moving FtsW–Halo molecule labeled with JFX650. Top: membrane staining (left), maximum intensity projection (MIP, middle), and kymograph of septal line scans (right). Bottom: extracted trajectories at the leading edge of S1 (dashed rectangle). Scale bars, 2 μm. **(C)** Representative trajectory of a directionally moving Halo–PBP1B molecule labeled with JFX650. Top: membrane staining (left), MIP (middle), and septal kymograph (right). Bottom: extracted trajectorie along S0 (dashed rectangle). Scale bars, 2 μm. **(D)** Fraction of directionally moving FtsW molecules at S1 under different conditions. Aztreonam significantly reduces directional motion, whereas FtsZD205G overexpression and cefsulodin show no significant effect. Values are mean ± s.d. from ≥ 3 experiments; **, P < 0.01, unpaired two-sample t-test (Welch’s test). **(E)** Fraction of directionally moving PBP1B molecules at S0 under different conditions. Aztreonam increases directional motion, whereas cefsulodin and moenomycin reduce it to ~1%. Statistical testing as in (D). **(F)** Speed distributions of directionally moving FtsW molecules. WT shows bimodal peaks at 2.7 nm/s and 15.5 nm/s. FtsZD205G reduces the fast-moving fraction and peak speed (12.7 nm/s). Aztreonam eliminates the slow-moving population, yielding a unimodal peak at 14.9 nm/s. Cefsulodin mildly reduces the slow-moving population. Black curves, lognormal fits. **(G)** Speed distributions of directionally moving PBP1B molecules. WT shows a unimodal peak at 12.3 nm/s. Aztreonam does not alter the distribution. Black curves, lognormal fits.

To test whether these populations couple to FtsZ treadmilling and PG synthesis, we perturbed both processes. Overexpression of an FtsZ mutant (D205G), which abolishes GTPase activity^59,60^, reduced FtsZ treadmilling speed from 22.0 ±0.4 nm/s to 6.5 ±0. 6 nm/s **(mean ±s.e.m.; Figure S10G, H; Movie S8, 9; Table S5)** and decreased the fraction of fast-moving FtsW molecules to 24.6% **(Figure 4F; Table S5)**. By contrast, inhibition of PBP3 with aztreonam eliminated the slow-moving population, collapsing the distribution into a single peak **(14.9 ± 1.3 nm/s; Figure 4F; Table S5)**. Together, these results indicate that fast-moving FtsW molecules track FtsZ treadmilling, whereas the slow-moving FtsW molecules are driven by active PG synthesis. The overall fraction of moving FtsW molecules also decreased when PBP3 was inhibited **(Figure 4D)**, likely due to stalled FtsW–PBP3 pairs entering futile synthesis cycles^14^. We note that the apparent slower speed of fast-moving FtsW compared to FtsZ likely reflects 2D projection of single-molecule trajectories, particularly in cells with highly constricted septa **(Figure S10I–L; Table S6; Methods)**.In contrast, PBP1B trajectories were enriched at closed septa (S0; 78.0 ± 9.2%) and, to a lesser extent, at the lagging edge of S1 (16.7 ± 7.6%) **(Figure 4A)**. Most PBP1B molecules were stationary, but a subset (9.5 ±1.4%) exhibited directional movement **(Figure 4C, E; Movie S10, 11)**. Unlike FtsW, moving PBP1B molecules displayed a unimodal speed distribution **(centered at 12.3 nm/s; Figure 4E, G; Table S5)**. Because FtsZ is absent from closed septa, this directional movement likely reflects processive PG synthesis, as recently reported for aPBPs in *S. pneumoniae*^13,24^. Supporting this view, inhibition of PBP1B’s TPase or GTase activity abolished the moving population **(Figure 4E; Table S5)**. Interestingly, aztreonam treatment did not affect the speed of PBP1B but substantially increased the fraction of moving trajectories **(Figure 4E, G)**, suggesting compensatory activation of PBP1B when PBP3 is inhibited. Collectively, the SMT results demonstrate that FtsW processively synthesizes the leading edge of S1 guided by FtsZ treadmilling, thereby driving constriction. By contrast, PBP1B acts at the lagging edge of S1 and in S0, also processively synthesize PG in the absence of the Z-ring.

### Secondary septal PG synthases by SaPBP2 in S. aureus

Having shown that PBP1B mediates a secondary septal PG synthesis on the primary septum formed by FtsW-PBP3 in *D. radiodurans*, we next asked whether this mechanism is conserved in other bacteria. Recent studies characterized biphasic changes in *S. aureus* septal architecture during constriction^61,62^, proposed to be associated with *Sa*FtsW-*Sa*PBP1 (the SEDS-bPBP pair in *S. aureus* divisome)^47^ and *Sa*PBP2 (the only class A PBP in *S. aureus*) ^63^, respectively. ***S. aureus*** undergoes three stages in its cell cycle with no septum (stage I), with a constricting septum (stage II) and a closed septum (stage III) **(Figure 5A)**. To examine the spatial distribution of the PG synthases, we co-expression of *Sa*FtsW– Halo and mNG–*Sa*PBP2 in methicillin-sensitive *S. aureus* (MSSA RN4220). Both proteins localized to the septum, but *Sa*PBP2 consistently positioned more the close to sidewall (analogous to the lagging edge in *D. radiodurans*), spatially separated from *Sa*FtsW (arrows in Figure 5B). Using *Sa*FtsZ as a marker, *Sa*FtsW strongly overlapped with the leading edge of septum throughout the constriction, whereas *Sa*PBP2 occupied a broader region on the septum behind **(Figure 5C; Figure S11A-D)**. This arrangement mirrors the spatially separation of FtsW and PBP1B in *D. radiodurans*.

**Fig. 5.**
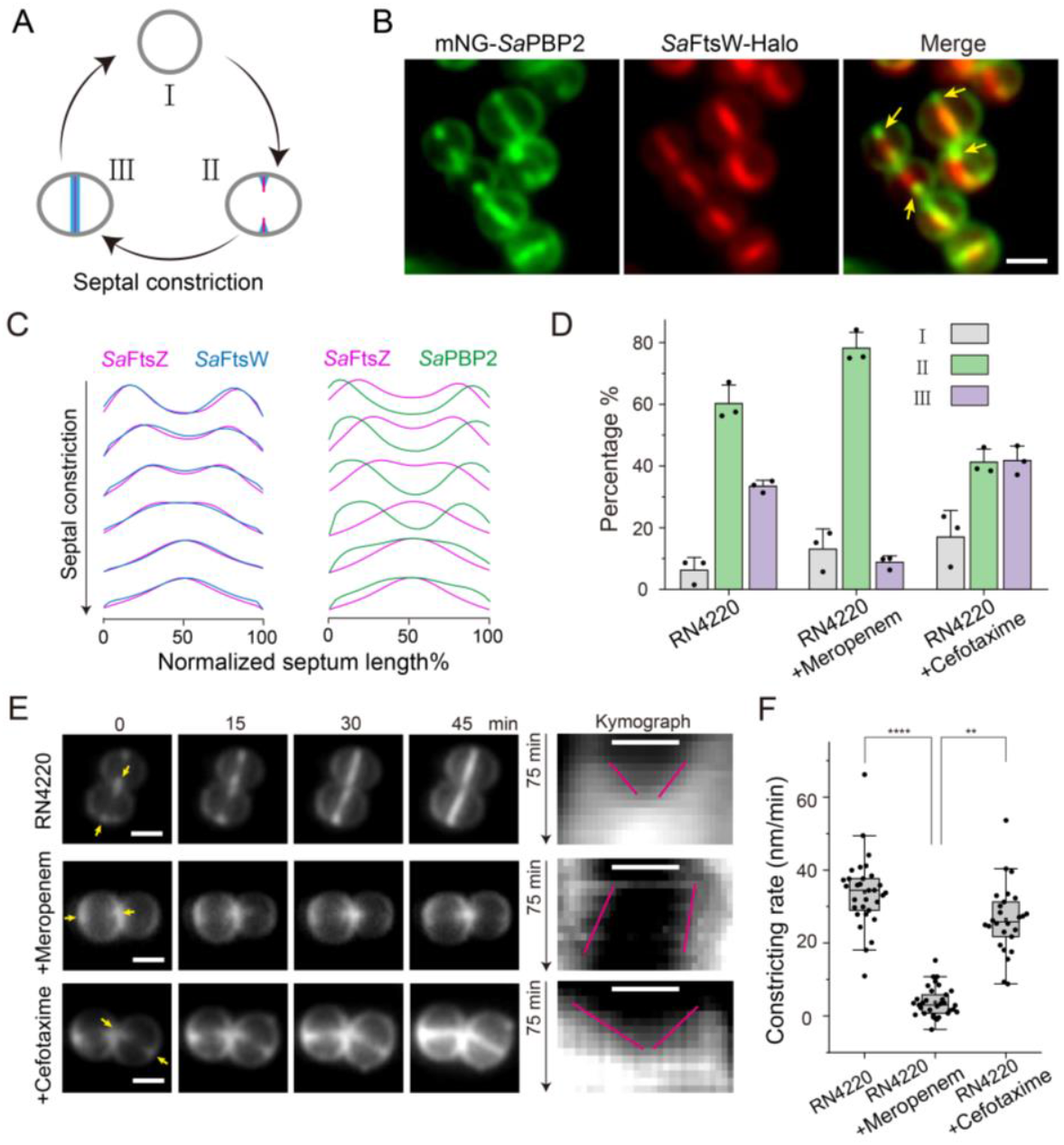
Spatially separated septal PG synthesis by *Sa*FtsW–*Sa*PBP1 and *Sa*PBP2 in *S. aureus*. **(A)** Schematic showing septal architecture change in *S. aureus* division cycle. Stage I: no septum; stage II: constricting septum; stage III: closed septum. **(B)** Spatial distribution of *Sa*FtsW (red) and *Sa*PBP2 (green). Both localize to the septum, but *Sa*PBP2 also shows peripheral membrane distribution. During constriction, *Sa*FtsW enriches at the leading edge, whereas *Sa*PBP2 localizes flanking *Sa*FtsW toward the sidewall (arrows). Scale bar, 1 μm. **(C)** Normalized mean intensity profiles of *Sa*FtsW (blue) and *Sa*PBP2 (green) compared with *Sa*FtsZ (magenta) that marks the leading edge of the constricting septum. *Sa*PBP2 consistently lies outside the *Sa*FtsZ ring. **(D)** Proportions of cells in stages I–III without treatment or after inhibition of *Sa*PBP1 (meropenem) or *Sa*PBP2 (cefotaxime). Meropenem increased the fraction of cells with unclosed septa. **(E)** Time-lapse imaging of septal constriction with or without drug treatment. Left: representative images; right: kymographs. Meropenem treatment stalls constriction. Scale bar, 500 nm. **(F)** Quantification of septal constriction rates from (E). Box plots show mean (squares), interquartile ranges (boxes), and 1.5× IQR (whiskers). **, P < 0.01; ****, P < 0.0001, unpaired two-sample t-test (Welch’s test).

To probe function, we inhibited their activity with different β-lactams. Meropenem, which targets *Sa*PBP1^64,65^, blocked constriction and caused accumulation of stage II cells with incomplete septa **(Figure 5D; Figure S11E)**. In contrast, inhibition of *Sa*PBP2 with cefotaxime^64,66^ allowed progression into stage III, though with fewer closed septa, while enrichment of stage I cells suggests that *Sa*PBP2 also contributes to septum initiation **(Figure 5D; Figure S11E)**^67^. Time-lapse imaging confirmed that meropenem nearly arrested constriction, whereas cefotaxime caused only a moderately reduced constriction rate **(Figure 5E; Movie S12-14)**. These results indicate that *Sa*FtsW–*Sa*PBP1 drives primary septal PG synthesis for constriction, whereas *Sa*PBP2 contributes an additional, secondary septal PG synthesis.

Lastly, we examined how inhibition these synthases affected septal PG cross-linking using FLIM. Without β-lactams treatment, septal PG exhibited higher cross-linking than the periphery cell wall, particularly concentrated at the septal center **(Figure S11E, F)**. This differs from *D. radiodurans*, possibly reflecting stronger hydrolytic activity at the lagging edge of the septum in *S. aureus*. Meropenem treatment, which blocked constriction, also produced incomplete septa with long lifetimes **(arrows in Figure S11E)**. Importantly, closed septa in cefotaxime-treated cells exhibited uniformly long lifetimes, including at the septal center **(asterisks in Figure S11E)**, demonstrating that *Sa*PBP2 carries out a secondary PG synthesis step that reinforces the septum formed by *Sa*FtsW–*Sa*PBP pairs.

## Discussion

Bacterial cytokinesis requires precise spatial and temporal control of septal PG synthesis. Although both SEDS–bPBP pairs and aPBPs are known to participate in septum formation, how their activities are coordinated has remained unresolved. *Deinococcus radiodurans*, a bacterium that combines Gram-positive–like peptidoglycan thickness with a Gram-negative–like outer membrane, provides a unique system to study septal PG synthesis because of its relatively large cell size and dual-septa morphology, containing both constricting septa (S1) and closed septa (S0)^35,36,41,42^. We show that its SEDS-bPBP pairs mediate primary synthesis at the leading edge of S1, generating inward septum constriction, while the division-specific aPBP acts at the lagging edge of S1 and persists in S0 to thicken the PG, increase cross-linking, and reinforce structural integrity. This spatiotemporal separation defines two steps of architecture development: a primary synthesis that drives constriction and a secondary synthesis that fortifies the septum **(Figure 6)**.

**Fig. 6.**
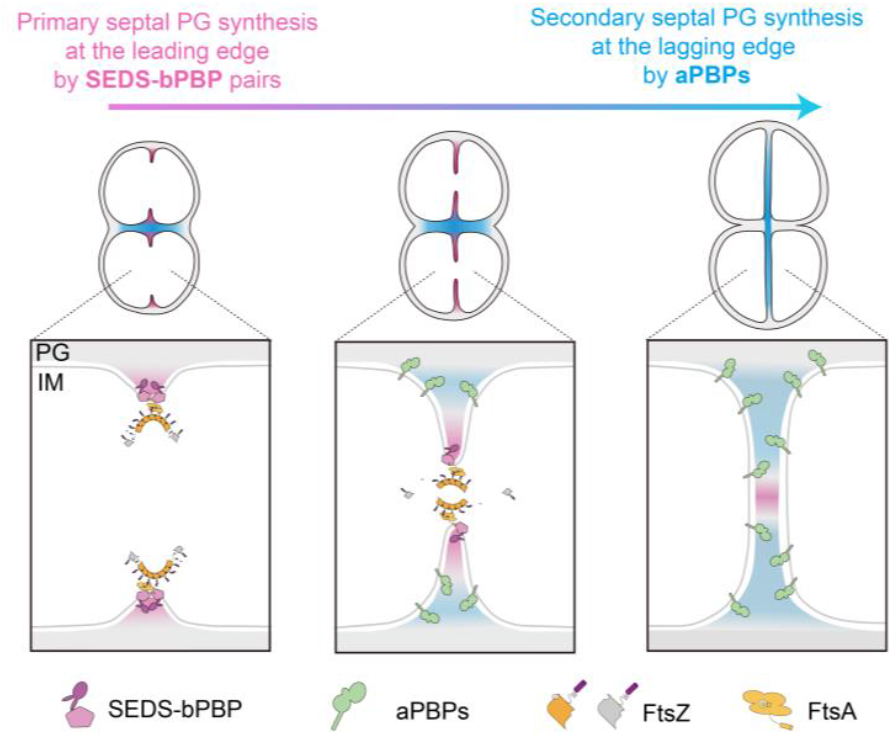
A working model of biphasic septal PG synthesis during bacterial cytokinesis. After septal initiation, division-specific SEDS–bPBP pairs (FtsW–PBP3 in D. radiodurans; SaFtsW–SaPBP1 in *S. aureus*) is recruited to the leading edge by the FtsZ ring. SEDS–bPBP drive primary septal PG synthesis, producing the thin leading edge that progresses inward to constrict the septum. Subsequently, class A PBPs (PBP1B in *D. radiodurans*; *Sa*PBP2 in *S. aureus*) accumulate at the lagging edge, where they execute secondary septal PG synthesis. This phase thickens and fortifies the septum by inserting additional PG and increasing crosslinking, continuing even after septal closure (S0). Together, these sequential activities generate a mature, mechanically stable septal wall.

### Spatial organization of synthases in D. radiodurans

Multi-color imaging and demograph analyses revealed that FtsW (SEDS) and PBP3 (bPBP) localize tightly with FtsZ at the leading edge of S1, consistent with their role as the canonical septal PG synthase complex demonstrated in other bacteria^14,19,47,68^. This colocalization holds throughout the septum constriction and dismissed from the closed septum with the disassembly of Z-ring. In contrast, PBP1B (aPBP) occupied a broader zone displaced toward the lagging edge and remained in S0 after S1 closure. This separation was difficult to detect in *E. coli*, probably due to the rapid constriction and separation of the septal cell wall.

### Distinct dynamics and biochemical signatures of the two septal PG synthases systems

Single-molecule tracking provided mechanistic insight into the *in vivo* activities of the two synthases systems. FtsW molecules at S1 exhibited both fast, FtsZ-dependent and slow, synthesis-dependent movements, consistent with previous observations in *E. coli* and *C. crescentus*^14,16^. By contrast, PBP1B molecules localized predominantly to the lagging edge of S1 and to closed septa (S0), where a subset exhibited clear directional movements. This motion was abolished upon inhibition of either its TPase or GTase activities, but remained unaffected by PBP3 inactivation, demonstrating that PBP1B functions independently of FtsW–PBP3 during secondary septal PG synthesis. Notably, blocking PBP3 increased PBP1B mobility, suggesting compensatory upregulation when primary synthesis is impaired. While aPBPs were traditionally thought to lack direction motion, recent work in *S. pneumoniae* identified processive dynamics of an elongasome aPBP^58^, echoing our observations of PBP1B. These results establish that aPBPs can act processively at the septum, reinforcing cell wall architecture during late cytokinesis.

Fluorescence lifetime imaging microscopy connected enzyme activity with PG chemistry. Regions enriched in PBP1B corresponded to higher cross-linking, detected as shorter lifetimes of fluorescent D-amino acid probes. Inhibition of PBP1B eliminated this highly cross-linked population, confirming its role in reinforcing the septal wall. By contrast, inhibition of PBP3’s TPase blocked constriction of S1 but left high crosslinking at the lagging edge, consistent with its role in primary septal synthesis. Thus, septal maturation in *D. radiodurans* proceeds through a secondary, PBP1B-dependent synthesis step. Importantly, loss of this secondary synthesis produced closed septa with reduced thickness and often caused membrane rupture at the lagging edge, highlighting its essential role in reinforcing septa to maintain mechanical stability.

### Conservation across bacteria

The sequential roles of SEDS–bPBP and aPBP type synthases extend beyond *D. radiodurans*. In *S. aureus* we observed a similar separation: *Sa*FtsW (SEDS) colocalized with *Sa*FtsZ at the leading edge of the constricting septum, whereas the sole aPBP, *Sa*PBP2, was distributed at the lagging edge of the septum and broadly across cell periphery, consistent with previous observations^63^. Perturbation experiments confirmed that *Sa*FtsW–*Sa*PBP1 pairs drive primary constriction, while *Sa*PBP2 contributes to additional septal PG crosslinking. Two distinctive septal architectures in *S. aureus* haven been revealed by atomic force microscopy (AFM): an initial septum composed of concentric ring structure and a later porous meshwork structure after further modification^61,62^, corroborating the two-step septal PG synthesis model in which *Sa*PBP2 executes the secondary synthesis. Likewise, in *E. coli*, the aPBP-type synthase *Ec*PBP1B was recently shown to fortifies the wedge-like structure at the base of the septum and protects against osmotic rupture^25^, mirroring the secondary synthesis step carried out by PBP1B in *D. radiodurans*. Together, these findings point to a conserved strategy of septal formation: SEDS–bPBP pairs advance the septum, while aPBPs reinforce and remodel the structure to meet the mechanical demands of closure and to maintain the integrity before separation **(Figure 6)**.

While our data establish distinct roles for SEDS–bPBP pairs and aPBPs, two key questions remain unresolved. First, the mechanism recruiting aPBPs to the lagging edge is unclear: although transient interactions with divisome proteins such as FtsA have been reported in *E. coli*^25^, it is not understood why aPBPs localize away from the Z-ring. One possibility is that the particular PG architecture synthesized by SEDS-bPBP pairs may provide the spatial cue to recruit aPBPs, but the underlying mechanism remains unknown. Second, the contribution of PG hydrolysis to this two-step architecture development is not fully resolved. In other bacterial species, hydrolysis has been shown to feedback on or regulate synthesis^25,69^. In *D. radiodurans*, our PBP4 knockout results indicates that the hydrolysis system actively remodels more at the lagging edge during cytokinesis. However, whether and how hydrolytic activity coordinates the septal PG synthase in *D. radiodurans* remains unclear.

## Materialsand Methods

### Bacteria culture and growth conditions

All strains used in this study are listed in Table S1. *E. coli* D5Hα strain was grown on Luria–Bertani agar (1% Tryptone, 0.5% yeast extract, 0.5% NaCl, 1% agar, Oxoid) or in Luria–Bertani broth at 37 °C with shaking at 220 rpm. *S. aureus* RN4220 strain was grown on tryptic soy agar (TSA, Difco) or in tryptic soy broth (TSB, Difco) at 37 °C with shaking at 220 rpm. All the strains of *D. radiodurans* were grown on 2×TGY agar (1% Tryptone, 0.2% Glucose, 0.6% Yeast extract, 1% Agar, Oxoid), in 2×TGY broth for routine cell culture or in MM minimal medium ^70^ for fluorescence imaging at 30 °C with shaking at 220 rpm. When necessary, media were supplemented with antibiotics at the following final concentrations: kanamycin at 20 µg/mL for *D. radiodurans* and 50 µg/mL for *E. coli*; chloramphenicol at 3 µg/mL for *D. radiodurans*, 15 µg/m L for *S. aureus*, and 70 µg/mL for *E. coli*; streptomycin at 20 µg/mL for *D. radiodurans*; and carbenicillin at 50 µg/mL for *E. coli*. For induction, 0.5 mM isopropyl β-D-1-thiogalactopyranoside (IPTG, BioFroxx) was used to induce both pCN5-derived vectors (Table S1) in *S. aureus* and the pET-28a(+)-derived vector in *E. coli* (Table S1), while 50 nM anhydrotetracycline (aTc, OriLeaf) was used for induction in *D. radiodurans*. Detailed growth and induction conditions for each experiment are listed in Table S3.

### Transformation of D. radiodurans cells

Competent *D. radiodurans* cells were prepared following the CaCl_2_ method for *E. coli* with modifications^71^. Briefly, mid-log phase cells (OD_600_ ≈ 1.0) were harvested by centrifugation at 5000 × *g* for 3 minutes. The cell pellet was gently resuspended and washed twice with 500 μL of buffer A (2×TGY medium supplemented with 60 mM CaCl2). Cells were then resuspended in 500 μL of buffer A and incubated with shaking at 30 °C for 90 minutes. Fresh competent cells were mixed with 100 ng of double stranded DNA (dsDNA) fragments or plasmids and incubated on ice for 30 minutes. The mixture was subsequently transferred to a glass tube containing 3 mL of 2×TGY medium and incubated at 30 °C with shaking for at least 8 hours. Finally, 100 μL of the culture was spread onto 2×TGY agar plates containing the appropriate antibiotics and incubated at 30 °C for approximately 48 hours until formation of single transformants.

### Construction of plasmids and strains

#### D. radiodurans strains

All strains used in this study were derivatives of the wild-type strain R1 (ATCC 13939)^72^. And all primers used for Polymerase Chain Reaction (PCR) are listed in Table S2. Chromosomal gene knockout, knockdown, and insertion were performed using a homologous recombination strategy similar to that described previously^73^. For gene knockout, two ~300 bp dsDNA fragments flanking the target gene and a streptomycin resistance cassette (*str*) were separately amplified by PCR, with approximately 20 bp overlapping sequences involved in the primers (Table S2). The fragments were then fused via overlap extension PCR to generate a dsDNA fragment containing the *str* cassette flanked by the upstream and downstream homologous arms^74^. Similarly, for the insertion of fluorescent protein (FP) or HaloTag encoded sequences, the upstream homologous sequence, *fp/halo* sequence, *str* sequence, and the downstream homologous sequence were fused by overlap extension PCR. The resulting PCR products were introduced into the DRWT via chemical transformation, followed by antibiotic selection.

Since *D. radiodurans* is polyploid, containing 4 to 10 genome equivalents per cell^75^, homologous recombination resulted in the replacement of one or multiple wild-type alleles in different transformants. To obtain strains with the target gene complete replaced, two to three rounds of re-streaking and antibiotic selection were performed. Genomic DNA from the resulting transformants was extracted using the TIANamp Bacteria DNA Kit (DP302), and mutant genotypes were confirmed by PCR (Figure S9A, B) and Sanger sequencing. Stable gene knockouts were successfully constructed for *pbp4* (JW116) and *pbp1A* (JW117). Other strains carrying chromosomally integrated alleles, including *ftsW-mng* (JW335), *ftsW-halo* (JW337), *mng-pbp1B*(JW378), *mcherry*(*mch*)^76^*-pbp1B*(JW152), and *halo-pbp1A* (JW375), were generated by introducing fusion constructs at the corresponding endogenous loci (Table S1). Doublestranded DNA fragments encoding *gfpmut2*^77^, *mng, mch*, and *halo* gene were amplified from pXY027, pXW001, pJH032, and pJH024. A flexible linker sequence (GGGGSPAPAPGGGGS, linker0^14^) was used between the protein of interest and the fusion tag.

#### D. radiodurans plasmids

All plasmids used in *D. radiodurans* in this study were derived from a shuttle plasmid, pJWK. pJWK is a minimal-size, single-resistance plasmid engineered from pRADK^78^. All modifications were performed via seamless cloning using the ClonExpress method (Vazyme, ClonExpress II One Step Cloning Kit, #C116). First, the kanamycin resistance gene (*aph*) in pRADK was replaced with the GFPmut2 coding sequence for quantitative assessment of the expression level. Next, the constitutive promoter P_groES_ was substituted with P_1261_, a native strong promoter from *D. radiodurans*^75^, yielding the intermediate plasmid pJW31. The ampicillin resistance gene (*bla*) in pJW31 was then removed. Additionally, the promoter P_groES_ (initiate transcription in both *D. radiodurans* and *E. coli*) was introduced to drive expression of a single chloramphenicol resistance gene (*cat*). The *orfB* and *orfH* genes were deleted to minimize plasmid backbone size, resulting in pJW32. More refinements of the plasmid included truncation of the AT-rich box region, removal of the residual *lac* promoter (yielding pJW33), and replacement of the chloramphenicol resistance cassette with a kanamycin cassette, culminating in the construction of pJWK, a compact 4,684 bp plasmid.

Subsequently, the constitutive P_1261_ promoter in pJWK was replaced with various inducible promoters, including P_spac1_^79^ and P_tetdr_^80^, generating the expression vectors pJW64, pJW65, and pJW67, respectively (Table S1).

Plasmids encoding fusion protein genes, including pJH208 (P_1261_::*ftsz-mng*), pJH150 (P_tetdr_::*ftsz-mng*), pJH158 (P_tetdr_::*mng-pbp3*), pJH195 (P_tetdr_::*N-mng-pbp3-ftsz-mch*), pJWK84 (P_tetdr_::*ftsz-mch*), pJWK81 (P_tetdr_::*mng-pbp1B*), pJWK7 (P_1261_::*mch-pbp1A*), and pJWK87 (P_spac1_::*halo-pbp1B*) were first constructed and verified in *E. coli* DH5α using the same seamless cloning method. Plasmids with single residue substitution pJH180 (P_tetdr_::*ftsz*^*D205G*^*-mng*) were introduced by sitedirected mutagenesis using primers listed in Table S2.

#### S. aureus plasmids and strains

The plasmid pCN5 used in this study for *S. aureus* was derived from pCNX^81^ through a two-step modification process. First, the kanamycin resistance gene in pCNX was replaced with a chloramphenicol resistance gene, resulting in the intermediate plasmid pCN1. Second, the inducible promoter P_cad-cadC_ in pCN1 was substituted with the inducible promoter P_spac_^3^, yielding pCN5.

The construction of fluorescent *S. aureus* strains was performed similarly to that of *D. radiodurans*. The strains JW461, JW460 and JW462 were constructed by introducing the pCN5 replicative plasmid encoding the transcriptionally linked genes fusion proteins: *SaftsW*-*halo*-(TAA)-*SaftsZ-mng, mng-Sapbp2-*(TAA)*-SaftsZ-halo* and *mng-Sapbp2-* (TAA)*-SaftsW-halo*. Using primers containing overlapping sequences, the dsDNA fragments encoding *S. aureus ftsW, ftsZ*, and *pbp2* were amplified from the genomic DNA extracted from RN4220 *S. aureus* strain, while *mng* and *halo* gene were amplified from plasmids pJH208 and pJWK87. These fragments were then linked with the amplified backbone of plasmid pCN5 using the ClonExpress method (Vazyme, ClonExpress II One Step Cloning Kit, #C116). Recombinant plasmids were first propagated in *E. coli* strain DH5α. Validated by colony-PCR and Sanger-sequencing, the purified plasmids were then introduced into electrocompetent *S. aureus* RN4220 cells according to a previously established protocol^82^.

#### E. coli plasmids and strains

All plasmids used in *E. coli* BL21 (DE3) were derived from the pET-28a(+) vector. The dsDNA fragments encoding *pbp1A*^*Δ1-73*^*-His*_*10*_, *pbp1B*^*Δ1-48*^*-His*_*10*_, and *pbp3*^*Δ1-35*^*-His*_*10*_ were amplified from genomic DNA of DRWT using primers with overlapping sequences. The *Twin-Strep-tag* gene was amplified separately from the plasmid pCAG-Hrd1-CS^83^. These fragments were assembled with the amplified pET-28a(+) backbone using the ClonExpress method (Vazyme, ClonExpress II One Step Cloning Kit, #C116). The recombinant plasmids, including pJH130 (P_T7_::*pbp1A*^*Δ1-73*^*-His*_*10*_*-Twin-Strep-tag*), pJH131 (P_T7_::*pbp1B*^*Δ1-48*^*-His*_*10*_*-Twin-Strep-tag*), and pJH132 (P_T7_::*pbp3*^*Δ1-35*^*-His*_*10*_*-Twin-Strep-tag*), were initially constructed and verified in *E. coli* DH5α using the same seamless cloning method. The purified plasmids were introduced into *E. coli* BL21 (DE3) via standard electro-transformation protocol^84^.

### Growth curve and Minimum inhibitory concentration (MIC) measurement

#### Growth curve measurement

The *D. radiodurans* strains were first allowed to grow to the stationary phase (OD_600_ ≈ 2.0) from 2×TGY agar plates supplemented with appropriate antibiotics in 2×TGY medium. The culture was then diluted back to an OD_600_ of ~ 0.02. 100 μL diluted cultures were dispensed into 96-well microplates. Growth was monitored by measuring absorbance at 600 nm every 5 minutes for ~24 hours at 30 °C using a Tecan SPARK 20M plate reader. Between each measurement, the plate was shaken for 15 seconds to ensure proper mixing. Three biological replicates were performed for each strain using independent single colonies.

#### MIC determination

The MIC of *D. radiodurans* was determined by the broth microdilution method in 2×TGY medium, following the guidelines of the Clinical and Laboratory Standards Institute (CLSI, 2018). Briefly, DRWT cells were grown to the stationary phase (OD_600_ ≈ 2.0) and then diluted in 2×TGY medium with antibiotics to an OD _600_ of ~ 0.04. The 2×TGY medium contained two-fold serial dilutions of aztreonam (from 0-6.4 µg/mL), cefsulodin (from 0-25 µg/mL), or moenomycin (from 0-1.64 µg/mL). Diluted cultures (100 μL per well) were dispensed into 96-well microplates. The assays were incubated at 30 °C for 40 hours. The MIC was defined as the lowest antibiotic concentration that completely inhibited visible growth of the DRWT (Table S7).

The MIC of *S. aureus* RN4220 was determined using the same microdilution method. Cultures were grown in TSB medium at 37 °C for 24 hours with serial dilutions of meropenem or cefotaxime, and growth curves were recorded to assess antibiotic sensitivity (Table S7).

### Antibiotic specificity

#### Protein purification

*D. radiodurans* PBP1A, PBP1B, and PBP3 were heterologously expressed in *E. coli* strain BL21 (DE3). Cultures were grown at 37 °C to OD _600_ ~ 0.7, then induced with 0.5 mM IPTG overnight (22 hours) at 16 °C. Cells were harvested by centrifugation at 4300 × *g* for 10 minutes, lysed by sonication in the presence of 1 mM phenylmethylsulfonyl fluoride (PMSF), and clarified by centrifugation at 10000 × *g* for 60 minutes at 4 °C. PBP1A and PBP1B were purified using Ni-NTA affinity chromatography (Qiagen), followed by a second purification step using Strep-Tactin resin (IBA Lifesciences). PBP3 was purified using Ni-NTA affinity chromatography (Qiagen), and subsequently dialyzed against Strep Wash Buffer (100 mM Tris, 150 mM NaCl, pH 8.0). Protein concentrations were determined by measuring absorbance at 280 nm and purity verified by SDS-PAGE.

#### Ampicillin-TMR competitive binding assay

Purified *D. radiodurans* PBPs (PBP1A, PBP1B, and PBP3) were diluted to 2 μM final concentration in 20 μL reaction volumes. Each reaction was supplemented with 2 μM ampicillin-TMR and increasing concentrations of aztreonam or cefsulodin. The mixtures were incubated at 30 °C for 60 minutes using a gradient thermal cycler (Biometra TAdvanced). Reactions were quenched by adding 5 μL of pre-heated 5×SDS loading buffer (250 mM Tris, 10% SDS, 50% glycerol, 0.5% bromophenol blue) followed by denaturation at 98 °C for 10 minutes. Samples were separated by SDS-PAGE, and fluorescently labeled PBPs were visualized using a multifunctional imager (Shenhua; AlexaFluor546 channel). Fluorescence intensities of target PBP bands were quantified in ImageJ^85^, with background subtraction. Band intensities were normalized to the no-competitor control.

Assuming that the reactions of each PBP with Ampicillin-TMR and the competing antibiotic (aztreonam or cefsulodin) follow unidirectional, first-order kinetics, the concentration of ampicillin–PBP complexes decreases as the concentration of aztreonam or cefsulodin increases. A more rapid reduction in the ampicillin-TMR signal indicates a higher relative reaction rate constant of the competing antibiotic with the PBP. To measure this rate, the fluorescence reduction of Amp-PBP was fitted using the following competitive binding model:

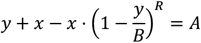

where ***y*** is the concentration of PBP bound to ampicillin-TMR, ***x*** is the concentration of the competing antibiotic (aztreonam or cefsulodin), ***A*** is the total PBP concentration (2 μM in this study), ***B*** is the initial concentration of ampicillin-TMR (2 μM), and ***R*** is the relative rate constant representing the competitive efficiency of aztreonam or cefsulodin versus ampicillin-TMR. Curve fitting was performed in OriginLab using data from three independent experiments. The resulting rate constant ratios ***R*** reflect the relative binding affinities of aztreonam and cefsulodin toward each PBP tested.

#### Measurements of fluorescence spectra

Fluorescence emission spectra of TMRDA and RhoDA were recorded using a microplate reader (Spark 20M, Tecan), with excitation at 500 nm and 488 nm (bandwidth: 20 nm), respectively. Emission signals were collected from 500 to 650 nm for TMRDA and from 520 to 680 nm for RhoDA (bandwidth: 20 nm), with a step size of 0.2 nm. Excitation spectra were measured by fixing the emission wavelength at 650 nm for TMRDA and 590 nm for RhoDA, over excitation ranges of 400-570 nm and 400-630 nm, respectively. To avoid reabsorption effects, the probes were diluted to 2 μM in HEPES buffer (50 mM HEPES, 120 mM NaCl, pH 7.3) containing 0.2% SDS and incubated at room temperature (25 °C) for at least 20 minutes ^43^.

FRET measurements in cell wall–labeled cells, pre-fixed cell suspensions (100 µL, OD_600_ = 2.0) labeled with different probes (see Cell Wall Labeling section below) were applied to 96-well plates (In Vitro Scientific). Emission spectra were recorded using a Spark microplate reader (Spark 20M, Tecan) at room temperature (25 °C). For cells co-labeled with RhoDA (donor) and TMRDA (acceptor), donor emission spectra (500-700 nm, bandwidth: 20 nm) were measured under varying acceptor concentrations with excitation at 488 nm (bandwidth: 20 nm). For cells labeled only with TMRDA (acceptor), emission spectra were recorded using the same excitation wavelength, scanning from 500 to 800 nm. Corresponding 1× PBS negative controls were included for background subtraction.

### Sample preparation for Light microscopy imaging

#### Bacterial cell preparation

Glycerol stocks of *D. radiodurans* strains were streaked onto 2×TGY agar plates supplemented with appropriate antibiotics and incubated at 30 °C for 72 hours. Single colonies were then inoculated into 2×TGY liquid medium and cultured overnight at 30 °C. Once the cultures reached the OD_600_ ≈ 2.0 the culture was diluted back to OD_600_ = 0.5 and allowed to grow for 2-4 hours until OD_600_ reaches ~ 1.0.

Glycerol stocks of *S. aureus* RN4220 strains were streaked onto TSA plates supplemented with appropriate antibiotics and incubated at 37 °C for 24 hours. Single colonies were then inoculated into TSB medium and cultured overnight at 37 °C. Once the cultures reached the OD_600_ ≈ 4.0, the culture was diluted back to OD_600_ = 0.2 and allowed to grow for 1-2 hours until OD_600_ ≈ 0.6.

#### Cell wall labeling

For cell wall (PG) labeling of *D. radiodurans*, 1 mL *D. radiodurans* cell culture (OD_600_≈1.0) was harvested by centrifugation at 7000 × *g* for 3 minutes. The cell pellet was washed with 1 mL of 2×TGY medium and then resuspended in 200 µL fresh 2×TGY supplemented with the appropriate antibiotic if needed. 2 µL of TMRDA, RhoDA, or Cy5DA from the −20 °C stocks were added to the culture, then incubated with agitation at 30 °C for varying durations. Probe concentrations and incubation times for each condition are listed in Table S3. For the PG synthases inhibition experiments, the 200 µL culture was pretreated with aztreonam (32.0 µg/mL, 10×MIC), cefsulodin (62.5 µg/mL, 5×MIC), or moenomycin (4.0 µg/mL, 10×MIC) for 1 hour prior to labeling. For the FRET experiment, 200 µL of cell culture was labeled with a mixture of 2 µL RhoDA (donor, 5 µM final concentration) and varying concentrations (0-100 µM final concentration) of TMRDA (acceptor). Acceptor-only and donor-only control samples for spectrometric calibration were prepared similarly, with concentrations detailed in Table S3.

For cell wall (PG) labeling of *S. aureus*, 1 mL *S. aureus* RN4220 cell culture (OD_600_ ≈ 0.6) was centrifuged at 7000 × *g* for 3 minutes. The cell pellet was washed with 1 mL of TSB medium and then resuspended in 200 µL fresh TSB. 2 µL RhoDA from the −20 °C stocks were added to the culture and incubated with agitation at 37 °C for 10 minutes. For the PG synthases inhibition experiments, the 200 µL culture was pretreated with meropenem (4 µg/mL, 2×MIC) or mefotaxime (2 µg/mL, 2×MIC) for 30 minutes prior to probe addition.

Labeled *D. radiodurans* and *S. aureus* cells were fixed following a previously described protocol^43^. Briefly, 460 µL of prechilled 100% ethanol was added to the labeled culture and mixed thoroughly by pipetting to immediately terminate the labeling reaction. Fixed cells were incubated on ice for 15 minutes, then washed three times with 1×PBS to remove residual ethanol and unbound probes. Finally, cells were resuspended in 10-30 µL of 1×PBS to a final OD_600_ of approximately 4.0 for imaging.

For live-cell PG labeling and time-lapse imaging, RhoDA were pre-mixed with molten agarose in TSB medium to generate a 1.5% agarose gel pad with 1 μM FADA_560_ ^43^. ~ 1 μL *S. aureus* cells were applied on the gel pad for subsequent imaging experiment^86^. Antibiotics were supplemented in both the cell culture and agarose gel if necessary.

#### Halotag labeling

For HaloTag labeling in *D. radiodurans*, 1 mL *D. radiodurans* cell culture (OD_600_ ≈ 1.0) was centrifuged at 7000 × *g* for 3 minutes. The cell pellet was washed once with 1 mL of 2×TGY medium and then resuspended in 200 µL of fresh 2×TGY supplemented with the appropriate antibiotic supplement. In colocalization experiments (Figure 1; Figure S5), 1 µL of 200 µM JF552 or JF503 HaloTag-ligand^87^ (a gift from Dr. Luke Lavis) was added to the suspension to a final concentration of 2 µM, and incubated at 30 °C for 40 minutes in the dark. In SMT experiments, 1 µL of 1 nM JFX650 HaloTag-ligand^88^ was added to reach a final concentration of 0.02 nM. The mixture was incubated at 30 °C for 30 minutes, then washed twice with 1 mL of MM medium. Cells were resuspended in 20 µL of MM medium to a final OD_600_ of approximately 4.0. To inhibit PBP3 or PBP1B, MM medium was pre-mixed with aztreonam (32.0 µg/mL), cefsulodin (62.5 µg/mL), or moenomycin (4.0 µg/mL).

To calibrate the systematic drift during single-molecule tracking, cells were fixed after Halotag-ligand labeling with 2.6% paraformaldehyde and 0.05% glutaraldehyde at room temperature (25 °C) for 15 minutes, followed by 30 minutes on ice^89^. Fixed cells were washed three times with 0.5 mL 1×PBS and resuspended in 1×PBS to a final OD_600_ of approximately 4.0.

For HaloTag labeling in *S. aureus*, 1 mL *S. aureus* cell culture (OD_600_ ≈ 0.6) was centrifuged at 7000 × *g* for 3 minutes. The cell pellet was washed once with 1 mL of TSB medium and then resuspended in 200 µL of fresh TSB supplemented with the appropriate antibiotic. In colocalization experiments (Figure 6; Figure S11), 1 µL of 200 µM JFX650 HaloTag-ligand was added to a final concentration of 2 µM, incubated at 37 °C for 30 minutes in the dark, then washed twice with 1 mL TSB:1×PBS (1:1) medium ^18^. Cells were resuspended in 20 µL TSB:1×PBS (1:1) medium to a final OD_600_ of approximately 4.0 for imaging.

#### Agarose gel pad preparation

For the FLIM and time-lapse imaging, 2% agarose (LONZA, 50111) gel pad preparation was adapted from our previous protocol^43^. Briefly, 120 mg of low melting point agarose gel was mixed with 600 μL of 1×PBS. The mixture was heated on a metal heat block at 70 °C for 1 hour to ensure complete melting. Approximately 100 μL of the molten agarose was applied onto a clean glass slide and sandwiched between a clean coverslip and a 0.5 mm thick rubber gasket (Bioptechs). The gel pad was allowed to solidify for at least 2 hours at room temperature (25 °C). Subsequently, ~ 1 μL of cell culture was applied onto the prepared gel pad, then covered with another clean coverslip for imaging. For FtsZ treadmilling experiments, 2% agarose gel pad was similarly prepared using 2×TGY medium in place of 1×PBS. For the colocalization experiment for *S. aureus*, 1.5% agarose gel pad was similarly prepared using TSB:1×PBS (1:1) medium in place of 1×PBS.

For the SMT, Potomac Gold dye^50^ (a gift from Dr. Luke Lavis) was used for cell membrane labeling. A 100 nM Potomac Gold solution was prepared by diluting the DMSO stock into MM medium and pre-warmed to 50 °C. Antibiotics or aTc were added if necessary. Separately, 3% molten agarose in MM medium was prepared as above and maintained at 50 °C. Equal volumes of the dye solution and molten agarose were gently mixed to obtain a final 1.5% agarose gel. Approximately 100 μL of this gel mixture was solidified and assembled in a Bioptech FCS2 chamber together with the cell culture^43^. For time-lapse imaging experiments, 2×TGY medium was used instead of MM medium, with 200 nM Potomac Gold dye in the gel pad.

For the colocalization experiment for *D. radiodurans*, 1.5% agarose gel pads were prepared in MM medium, but containing 2 µM Potomac Red dye for membrane staining. In all experiments involving membrane dye labeling, cells were incubated on the gel pad for at least 20 minutes at room temperature (25 °C) in the dark prior to imaging.

For 3D-dSTORM experiments, 3% agarose in 20% D-glucose solution was melted and maintained at 50 °C. Meanwhile, 2×STORM imaging buffer (100 mM Tris-HCl pH 8.0, 100 mM NaCl, 0.2 M Cysteamine (Sigma-Aldrich/Merck, 30070-10G), 4.1 mg/mL glucose oxidase (Sigma-Aldrich/Merck, G2133-10KU) and 0.17 mg/mL catalase (Sigma-Aldrich/Merck, C1345-1G)) was pre-warmed in a 37 °C water bath. Finally, mix 300 µL of the pre-warmed 3% agarose solution with an equal volume of 2×STORM buffer, and vortex thoroughly to obtain 1.5% agarose gel (GLUX) for dSTORM imaging.

#### Fluorescent beads sample preparation

Multicolor fluorescent bead sample was prepared to calibrate chromatic aberrations across different fluorescence channels. TetraSpeck™ Microspheres (0.1 μm, Thermo Fisher) were diluted 1:100 in ddH2O by vortexing. 10 μL of the bead suspension was applied onto a D-lysine-pretreated glass slide (prepared by incubating slides in 0.1% Poly-D-lysine solution (Sigma-Aldrich/Merck, P6407-5MG)) at 25 °C for 6-8 hours and allowed to settle for 10 minutes. A square D-lysine-pretreated coverslip was carefully placed over the droplet and sealed with mounting resin to create a stable bead sandwich for imaging.

### Light microscopy

#### Fluorescence lifetime imaging (FLIM)

FLIM was performed using on a Leica STELLARIS 8 confocal microscope equipped with an HC PL APO 100×OIL CS2 objective (NA = 1.4, Refractive Index = 1.52), white light laser, and HyD S detectors. All images were acquired with a zoom factor of 3.00 and a pinhole size of 1.0 Airy unit at 580 nm (147.8 µm). Excitation wavelengths were set to 549 nm for TMRDA and 488 nm for RhoDA. Detailed imaging parameters are provided in Table S3.

#### HiLO- and TIRF-imaging

HiLO (Highly inclined and laminated optical sheet) and TIRF (total internal reflective fluorescence) imaging were performed on a Nikon ECLIPSE Ti2 inverted microscope equipped with a 100×oil-immersion TIRF objective (NA = 1.49) and either a BSI or a Prime95B sCMOS camera (Teledyne Photometrics). HiLO or TIRF illumination mode was achieved and switched using the MTW150 TIRF illuminator (Beijing Coolight Technology). Excitation was provided by a solid-state laser engine (MULT4, Beijing Coolight Technology) with three continuous-wave lasers at 488, 561, and 647 nm, targeting mNG and HaloTag ligands JF503 (488 nm), JF552, mCh, and Potomac Gold (561 nm), as well as Potomac Red (647 nm). Fluorescence signals were collected using a ZT405/488/561/647rpc dichroic mirror and a ZET405/488/561/647m emission filter set (Chroma Technology).

For two- or three-color imaging of *D. radiodurans*, the BSI camera was operated in HDR mode with an additional 1.5×magnification, yielding a pixel size of 43.3 nm. Sequential single-color images were collected with additional emission filters (ET525/50m, ET605/70m, and ET705/72m) associated with 488, 561, and 647 nm excitation, respectively. Brightfield images were captured alongside fluorescence images for reference. Excitation laser power and exposure times are detailed in Table S3. Z-stack imaging was performed with a 0.2 µm step size over a range of −0.5 µm to +0.5 µm relative to the cell center. For *S. aureus* imaging, single-slice imaging with TIRF illumination was used to minimize cytosolic background.

For single-color, time-lapse imaging, no extra magnification was applied, resulting in a 65 nm pixel size. Images were acquired at a single focal plane in the 561 nm channel (13 W/cm^2^ power and ET605/70m filter) with 50 ms exposure every 10 minutes for approximately 7 hours. To maintain physiological conditions, the FCS2 chamber and objective heater (Bioptech) were set to 30 °C. HiLO illumination was used for *D. radioduran*s while TIRF was used for *S. aureus*.

For SMT, fluorescence emission in the 561 and 647 nm channels was split by an Optosplit II (Cairn Research) equipped with a T647lpxr dichroic mirror plus ET605/50m and ET705/70m emission filters in each channel (Chroma Technology), projected onto the Prime95B camera (sensitivity mode). No additional magnification was used, resulting in a pixel size of 110 nm. The focal plane was set approximately 100 nm below the cell center, identified via brightfield images. Potomac Gold-labeled membranes were imaged for 60 frames with 50 ms exposure using the 561 nm laser at 90 W/cm^2^. Subsequently, single molecules tagged with JFX650 dye were tracked for 200 frames with 1-second exposure time without dark intervals, illuminated by 647 nm laser at 5 W/cm^2^. SMT experiments were conducted at 27 °C.

For every multi-color imaging experiment, A TetraSpeck beads sample was imaged sequentially across channels with 50 ms exposure for chromatic calibration. Multiple images were taken for each experiment to assure enough beads covering the entire field of view.

For FtsZ treadmilling imaging, the BSI camera was operated in HDR mode with an additional 1.5×magnification, yielding a final pixel size of 43.3 nm. FtsZ-mNG or FtsZ^D205G^-mNG was excited at a single focal plane near the bottom of the cells using the 488-nm laser at 21 W/cm^2^under TIRF illumination and wit h ET525/50m filter. The bright field and fluorescence images were acquired alternatively in 1-second intervals with an exposure time of 50 milliseconds for 100 frames. Experiments were conducted at 27 °C.

#### 3D-SMLM imaging

A 10 μL aliquot of fixed cell suspension (washed with 1×PBS, OD _600_ ≈ 0.5) was placed at the center of a 35-mm glass-bottom dish and allowed to settle for 5 minutes. Excess liquid was removed, and the sample was allowed to dry for 10 minutes. Subsequently, 600 μL of freshly prepared 1.5% agarose gel (GLUX) was applied on top, and a 22-mm square coverslip was gently placed on top and pressed to ensure close contact with the dish bottom. After solidification (~10 min at room temperature), the sample was transferred to the microscope stage for equilibration.

3D SMLM data were collected using a custom-built single-objective microscope^90^. A 642 nm laser (MPB Communications) was first passed through a clean-up filter (ZET640/10x, Chroma) and then expanded using a 10× achromatic beam expander (GBE-10A, Thorlabs). The laser beam was subsequently filtered by another laser filter (ZET405/488/561/635m, Chroma) before being focused onto the back focal plane of an oil-immersion objective lens (U Plan XApochromat 100×/1.45 NA, Olympus) via an achromatic lens (AC508-180-A, Thorlabs), enabling wide-field illumination. The focusing lens could be adjusted perpendicularly to the laser beam, allowing for illumination mode switching between epi-, TIRF-, and HiLO-illumination. Between the focusing lens and the objective, a 4-band dichroic mirror (ZET405/488/561/640rpcv2, Chroma) was installed to separate the excitation and emission paths. Additionally, a 405 nm laser (OBIS 405nm LX, Coherent) worked with the 642 nm laser to reactivate fluorophores from their radical (dark) state.

The emitted photons were collected by the same objective and focused via a tube lens (f = 200 mm) into the channel-splitter (OptoSplit II, Cairn). A cylindrical lens (f = 10,000 mm, CVI Laser Optics) was put in the infinite space between the objective and the tube lens to introduce astigmatism. Inside the channel-splitter, a 685 nm long pass dichroic (T685lpxr, Chroma) coupled with a 667/30 (ET667/30, Chroma) and 690/50 (ET690/50, Chroma) band-pass filter in the reflection and transmission channels, respectively, were placed to achieve the ratiometric multicolor imaging. The fluorescence was then focused into an sCMOS camera (Prime95B, Teledyne Photometrics). The intensity density of the 642 nm laser was calibrated to about 1.0 kW/cm^2^ specifically for the ratio-metric multicolor SMLM imaging. For each imaging, data acquisition consisted of 30,000 frames, acquired at 25 ms per frame, with persistent reactivation of the dyes facilitated by the 405 nm laser.

### Light microscopy data processing

#### Pre-processing

For multi-color imaging, images comprising four channels (Channel 1: brightfield, Channel 2: 647 nm, Channel 3: 561 nm, Channel 4: 488 nm) are pre-processed using a MATLAB–Fiji interface via Miji^91^. For each channel, image slices are combined into a hyperstack containing Z slices associated with four colors. The x-y plan drift was corrected using the *HyperStackReg* plugin^92^. Next, the chromatic aberration between channels were calibrated using the TetraSpeck bead images. Geometric transformation matrices for the 647-nm and 488-nm channels, relative to the 561-nm channel, were calculated using the MATLAB function *fitgeotrans* and applied to the experimental datasets to generate calibrated images (see Data and Code Availability). For single-molecule tracking, the 561-nm channel was registered to the 647-nm channel using the same procedure.

Except for FLIM and SMT images, all original images were then denoised using the Puredenoise plugin^93^ in ImageJ^85^.

#### Image segmentation and cell cycle classification

Cell segmentation was performed using Cellpose 3.0^94^. For fluorescence images of membrane or cell wall labeling, the pre-trained “cyto3” model was applied with manual refinement as necessary. For brightfield images acquired during SMT experiments, the “cyto3” model was re-trained using one or two representative images to improve segmentation accuracy. The resulting cell masks were then used for downstream analyses, including cell cycle classification and registration of singlemolecule trajectories.

For cell cycle classification, we constructed a convolution neuron network (CNN) based on ResNet^95^. A training dataset with approximate 1000 segmented *D. radiodurans* cells were manually classified into five stages over the cell cycle (Figure 1B). 30 epochs of training resulted a mode called DeCNN (**De**inococcus **c**lassification **n**euron **n**etwork), perform with ~92% accuracy of cell cycle classification. The mode and instruction are available on: https://github.com/BacImgLab/DeCNN. Cellpose segmented masks were used to outline individual cells in either bright field images or fluorescence images. Each cell can be classified to one of the five stages by DeCNN automatically.

#### Demograph construction

To construct septal demographs in *D. radiodurans*, cells at stages 3 and 4 were selected for profiling S1 septa, while cells at stages 2 through 4 were used for S0 analysis. Stage 1 cells were excluded as they are likely artifacts caused by mechanical disturbance during sample preparation, and stage 5 cells were omitted due to their low abundance.

To sort and align the cells across cytokinesis, septal morphology was first determined based on membrane or cell wall labeling (Figure S2A). To quantify S1 septal growth, a seven-pixel-wide (~ 300 nm) line was drawn across the constricting septum. The length of septum was recorded as ***D***_**S1**_. The septal gap (***d***_**S1**_) was calculated from the fluorescence intensity profile along this line, defined as the distance between two points where the signal dropped to half-maximum intensity on either side of the septum (Figure S2A). The ratio ***d***_**S1**_**/*D***_**S1**_ served as an indicator of septal constriction progression (Figure S2B). To quantify the development of S0, the diameter of the cell, denoted as ***D***_**S0**_, was measured as the width of the minimum bounding rectangle enclosing the segmented cell. The length of the closed septum (***d***_**S0**_) was manually defined based on the membrane signal. The ratio ***d***_**S0**_**/*D***_**S0**_ was used to estimate the progression of S0 hydrolysis and final septal separation (Figure S2B).

Normalized intensity profiles along S0 and S1 were sorted in descending order based on their respective ***d***_**S1**_**/*D***_**S1**_ and ***d***_**S0**_**/*D***_**S0**_ values, respectively. The profiles were then centered and aligned to produce raw demographs. These were subsequently binned into 80 rows to standardize across datasets and enable comparisons between experiments. For FLIM data, demographs were constructed using the same method, except that the profiles represent average fluorescence lifetimes rather than intensity, and no normalization was applied. Scripts for demograph construction are available at https://github.com/BacImgLab/DRsPGsynthesisCode.

Demographs for *S. aureus* were generated using a similar approach. Five-pixel-wide lines (~ 210 nm) were drawn across septa to extract Z-ring intensity profiles. Each profile was fit was fitted with a pair of Voigt functions to obtain the centers of Z-ring on each side of the constricting septum. The distance between the centers decreased as constriction progressed and was used to sort intensity profiles of *Sa*FtsW and *Sa*PBP2, which were normalized to generate demographs (Figure S11B, D). Demographs were grouped into six stages, producing the averaged profiles of the protein distribution over constriction process.

#### Pearson’s correlation coefficient (PCC) calculation

The colocalization level between channels was demonstrated by PCC values calculated from individual cells, using the equation below:

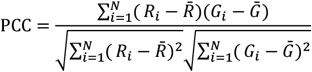

Where *R*_i_ and *G*_i_ are the intensity values of the i-th pixel in the two channels, respectively; 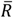 and 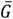 are the mean intensity values of all pixels in the two channels; N is the total number of pixels. To eliminate the influence of background noise, all images were corrected by subtracting the mean value of dark images (approximately 100). Cells with very low or very high overall intensity (below or above the 95%) were excluded to avoid false-negative or false-positive colocalization results.

#### FLIM data analysis

Each FLIM image was exported as a fluorescence intensity image (intensity range: 0-65535) and a fluorescence lifetime image (lifetime range: 0-10 ns), and subsequent statistical analysis was conducted in MATLAB. To remove background, the exported fluorescence intensity images were binarized using the built-in local thresholding algorithms in Fiji (Bernsen for *D. radiodurans* and Median for *S. aureus*). The generated mask was then applied to the fluorescence lifetime images to remove all the pixels not on the cell envelop. Lifetime and intensity values of all remaining pixels were extracted for subsequent distribution analysis.

#### FRET efficiency calculation

To quantify FRET efficiency based on fluorescence intensity, the emission spectra of the sample with donor and acceptor were subtracted by the spectra with only acceptor, generating the *FI*_DA_ curves in Figure 1E. The apparent FRET efficiency (Figure 1F) was then calculated as:

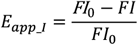

where *FI*_0_is the donor emission intensity at 530 nm in the absence of acceptor, and *FI* is that in the presence of acceptor.

For lifetime-based FRET (FLIM-FRET) efficiency, pixels with intensity above threshold in the donor channel were used to calculate the FRET efficiency as:

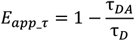

where *τ*_DA_ is the fluorescence lifetime of the donor in the presence of the acceptor, and *τ*_D_ is the donor lifetime in the absence of acceptor. Lifetimes were extracted from segmented septal regions of individual cells using a biexponential fitting model when applicable.

To calculate the average distance between donor and acceptor under different conditions. The Förster distance (*R*_0_) between the donor (RhoDA) and acceptor (TMRDA) was first calculated based on their spectral properties using the equation:

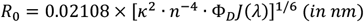

where *κ*^2^ is the orientation factor (assumed to be 2/3 here), *n* is the refractive index of the medium (assumed to be 1.33 here), *Φ*_D_ is the quantum yield of the donor (0.95 for RhoDA), and J (λ) is the spectral overlap integral between the donor emission and acceptor absorption spectra, measured in Figure S1D. This yielded an *R*_0_ of approximately 6.14 nm for the RhoDA–TMRDA pair.

#### FtsZ treadmilling speed analysis

The analysis was performed as previously described^60^. Briefly, Raw time-lapse fluorescence image stacks were first denoised using the PureDenoise plugin in Fiji^96^ and then enlarged to a pixel size of 21.7 nm using the bicubic interpolation method. Drift correction was applied using the StackReg plugin^97^. The denoised stacks were then smoothed through a moving average over a 3-frame window. Kymographs were generated using the kymograph plugin (written by J. Rietdorf and A. Seitz) with a nine-pixel-wide (~ 195 nm) line along the Z ring. The polymerization or depolymerization speeds of the FtsZ clusters were calculated by manually measuring the slopes of the leading or trailing edges of the fluorescence zigzags in the kymographs.

#### SMLM data analysis

3D SMLM images were localized using a multi-emitter fitting strategy as previously described^90^. Briefly, raw camera frames were first converted to electron count maps based on the manufacturer’s specifications. Local maxima in each electron count map were identified using a Difference-of-Gaussian (DoG) filter. Around each local maximum, a 13 × 13 pixel^2^ region of interest (ROI) was cropped and sent to a GPU (NVIDIA RTX 3070) for fast multi-emitter fitting.

Each ROI was fit to a *K*-component point spread function (PSF) mixture model (PMM). The 3D PSF was constructed by sub-pixel alignment and robust normalization of several bright fluorescent beads (T7279, ThermoFisher), and then represented using cubic spline interpolation. The *K*-component PMM was parameterized by **Θ** = {**θ**_1_, … **θ**_*K*_}, where each component **θ**_*k*_ = {*N*_*k*_, *x*_*k*_, *y*_*k*_, *z*_*k*_} includes the photon count *N*_*k*_ and 3D position *x*_*k*_, *y*_*k*_, *z*_*k*_ of the *k*-th emitter. The expected photon count at pixel *i* was modeled as *μ*_*i*_ = ∑_*k*_ *N*_*k*_PSF(**r**_***i***_|*x*_*k*_, *y*_*k*_, *z*_*k*_), where **r**_***i***_ is the position of pixel *i* relative to emitter *k*. The likelihood of each ROI was modeled using a Poisson distribution: *p* = ∏_*i*_ Poisson(*d*_*i*_|*μ*_*i*_) and the parameters **Θ** were optimized by minimizing the negative log-likelihood using the Levenberg–Marquardt algorithm.

The number of components *K* was determined via Maximum Likelihood Estimation (MLE) penalized by the Bayesian Information Criterion (BIC) to avoid overfitting. The resulting localizations were corrected for 3D drift using Redundant Cross-Correlation (RCC), followed by a consecutive-frame grouping protocol to merge localizations that appeared at the same position across sequential frames.

#### SMT data analysis

Each SMT dataset consisted of three channels: brightfield (BF), membrane fluorescence (561 nm), and single-molecule localizations (647 nm). Datasets with drift exceeding 3 pixels in the brightfield channel over the 200-second imaging period were excluded to prevent false interpretation of molecular movement due to sample or stage instability. Membrane fluorescence frames were averaged to generate a high signal-to-noise image for accurate cell segmentation, stage classification, and trajectory registration.

Single-molecule localization and tracking in 2D were performed following previously described procedures^98^. Briefly, a Gaussian sigma filter (< 250 nm) and intensity filter (range: 120-700) were applied to remove single-pixel noise and out-of-focus fluorescence after singlemolecule localization by ThunderStorm in Fiji^99^. Localizations were connected into trajectories using a maximum distance threshold of 300 nm per frame and a dark interval threshold of 6 frames.

Individual cells were segmented based on membrane labeling and assigned to different cell cycle stages. Single-molecule trajectories were overlaid on the corresponding cell outlines and manually categorized as localizing to the leading or lagging edge of the unclosed septum (S1), or to the closed septum (S0).

To analyze molecular dynamics, we applied a previously established motion classification algorithm^14^ to segment each trajectory into directionally moving or stationary states. Directional segments with speeds below 1 nm/s were excluded to eliminate potential artifacts from residual drift. For each condition, the proportions of trajectories localized to the leading edge of S1, the lagging edge of S1, and S0 were calculated based on the number of segments in these regions.

The percentage of molecules stay in the directional movement state was averaged across three to five biological replicates. Speed distributions of the directionally moving segments were analyzed using cumulative probability plots and fitted with single-or double-population lognormal functions, as described^14^. Standard error of the mean (s.e.m.) for the average speed and directional fraction was estimated by bootstrapping (2,000 iterations).

2D-projection correction (unwrapping) of the single-molecule trajectories were processed using a similar method to our previously study on *Ec*FtsW^14^. Instead of using super-resolution imaging to calibrate the radius of the constriction ring, we used the membrane staining channel to estimate the radius (Figure S10I, J). Considering the low resolution, a 150% larger ring was also used to demonstrate how the ring size affects the velocity estimation. Due to the lack of clear spatial landmarks of the closed septum S0, no unwrapping process was performed on the PBP1B trajectories located in S0. Scripts for the 2D-projection correction in this study are available at https://github.com/BacImgLab/DRsPGsynthesisCode.

### Room temperature electron tomography

#### Ultrathin Section Sample Preparation

The overnight culture of wild-type *D. radiodurans* was diluted to OD_600_ ~ 0.2 and incubated at 30 °C with shaking for another 2-4 hours until the OD_600_ reach ~ 1.0. Then, 1 mL of the culture was harvested by centrifugation at 7000 × *g* for 3 minutes. The bacterial pellet was resuspended in 200 µL of fresh 2×TGY medium supplemented with necessary antibiotics to inhibit specific PG synthase and incubated at 30 °C for 1 hour. The final concentration of each antibiotic was kept at 5×MIC (16 µg/mL aztreonam, 62.5 µg/mL cefsulodin, or 2 µg/mL moenomycin. After washing by 1×PBS buffer, the cells were then resuspended in 1 mL of fixation buffer (5% glutaraldehyde and 4% paraformaldehyde in 1×PBS) and rotated for 2 hours at room temperature, followed by overnight fixation at 4 °C. After fixation, the cells were washed three times with 1×PBS and resuspended in 4% osmium tetroxide for another overnight incubation at room temperature. The cells were then washed three times with 1×PBS and dehydrated using a graded ethanol series (20%, 50%, 75%, 80%, 95%, 100%), each step lasting 20 minutes with gentle mixing. Next, the cells were washed three times with acetone and embedded in Spurr’s resin in a graded series (25% for 24 hours, 50% for 24 hours 75% for 24 hours, and 100% for 120 hours), replacing the resin with fresh room temperature balanced resin every 7-8 hours. Polymerization was carried out in a 70 °C incubator for 48 hours^100^. Ultrathin sections (200 nm) were generated with an EM UC7 ultramicrotome (Leica), mounted on copper palladium slot grids coated with 1% formvar (Agar Scientific) in chloroform and post-stained with uranyl acetate (4%, w/v) and Reynold’s lead citrate (3%, w/v) for 5 minutes each.

#### Electron tomography imaging

Transmission electron microscopy imaging was performed at 200 kV using a Thermo Scientific Glacios Cryo-Transmission Electron Microscope equipped with a Falcon3EC camera. A single-axis tilt scheme with alternating tilting directions was employed, covering an angular range from −60° to +60° at 3° intervals. Each image was acquired at 17,500 ×magnification with a 1-second exposure time.

#### Electron microscopy image data analysis

Raw tomographic tilt series from different angular views were first aligned and drift-corrected using internal cellular landmarks with AreTomo^101^, which enables accurate motion correction based on fiducial-free registration. From each reconstructed tomographic tilt series, the most focused and structurally informative 2D slice—typically corresponding to the central plane of the septum—was manually selected for morphological analysis.

Within these selected tomographic slices, S0 and S1 septa were identified based on their morphology. Septal contours were segmented by tracing the edges of highest electron density, corresponding to the peptidoglycan cell wall. A centerline was then fitted along the long axis of each septum in1-nm steps.

At each centerline position, septal width was calculated by measuring the perpendicular distance between the inner edges of the septal wall using a custom MATLAB script *TomowidthCaclu*. This process generated a one-dimensional profile of septal thickness along the septum length. The average septal width was determined by averaging all thickness values along the centerline. These values were used for quantifying differences in septal architecture under different experimental conditions (e.g., drug treatments).

#### Structural prediction of sPG synthases

Full-length PBP1A, PBP1B, PBP3, PBP4, and FtsW were modeled using the AlphaFold3 server with default settings^102^, providing the amino acid sequences as input and specifying the monomeric form. For comparison, *Ec*PBP1A, *Ec*PBP1B, *Ec*PBP3, *Ec*PBP4, and *Ec*FtsW were modeled in the same manner.

## End Matter

### Author Contributions and Notes

Conceptualization: J.W., X.Y.

Methodology: J.W., J.H., Y. X., L. D., X. W., J. P., Y. L., L. Z., C.Y., Y. G., T. X., Y. Y., J. Y., L. X., X. Y.

Investigation: J.W., J.H., H. W., X.Y.

Software: J.W., X. O., X.Y.

Formal analysis: J.W., J.H., Y. X., J. P., S. Z., L. Z., Y. Y., X. Y.

Writing – original draft: J.W., X. Y.

Writing – review & editing: J.W., J.H., Y. Y., L. X., X. Y.

Supervision: Y. Y., J. Y., L. X., X. Y.

Funding acquisition: Y. Y., J. Y., L. X., X. Y.

## Acknowledgments

The authors thank other members in Yang, Xue, and Yin’s laboratory for their valuable advice and discussions. The authors thank Dr. Mariana G. Pinho, Dr. Hongwu Qian, and Dr. Yuejing Hua for providing bacteria strains and plasmids. The authors thank Dr. Luke D. Lavis for sharing JF dyes and Potomac dyes for the protein and membrane labeling. The authors thank Dr. Jie Xiao, Dr. Shishen Du, Dr. Mariana G. Pinho, and Dr. Amilcar Perez for helpful suggestions and discussions. The authors would like to acknowledge the Confocal Imaging Unit at the Core Facility Centre for Life Science, University of Science and Technology of China. The Laboratory research staff (Zhenbang Liu, M.S.) contributed valuable technical expertise and assistance to this project.

This work was partially carried out at the Instruments Center for Physical Science, University of Science and Technology of China. This work was supported by the National Natural Science Foundation of China (32270035 to X.Y., 22377115 to L.X.), Anhui Provincial Natural Science Foundation (2208085MC49 to L.X., 2208085MC40 to X.Y., and 2008085QC98 to J.Y.), the Center for Advanced Interdisciplinary Science and Biomedicine of IHM (QYPY20220016 to L.X.), the Fundamental Research Funds for the Central Universities WK9100000063 to X.Y., Basic and Applied Basic Research Foundation of Guangdong Province (2024A1515012105) to Y.Y., Shenzhen Medical Research Fund (B232018) and Shenzhen Bay Laboratory Startup Fund (21230101) to Y.Y., and the USTC start-up funding (KY9100000033, KY9100000068 KJ2070000081 to L.X. and KY9100000035, KJ2070000083 to X.Y.).

## Resource availability

### Lead contact

Information and requests for reagents may be directed to, and will be fulfilled by, the lead contact, Xinxing Yang.

### Materials availability

Strains, plasmids, and probes are available on request from the lead contact with a materials transfer agreement.

### Data and code availability

Code used in this study can be accessed at the following GitHub repository: https://github.com/BacImgLab/DRsPGsynthesisCode.

