## Supplemental File for "Class A PBPs reinforce the septal cell wall following initial synthesis by SEDS–bPBP pairs during bacterial cytokinesis"

**This PDF file includes:**

**Figures S1 to S11**

**Supplementary Table 1 to 7**

**General considerations of chemistry**

**Captions for Movies S1 to S14**

**References**

**NMR spectra**

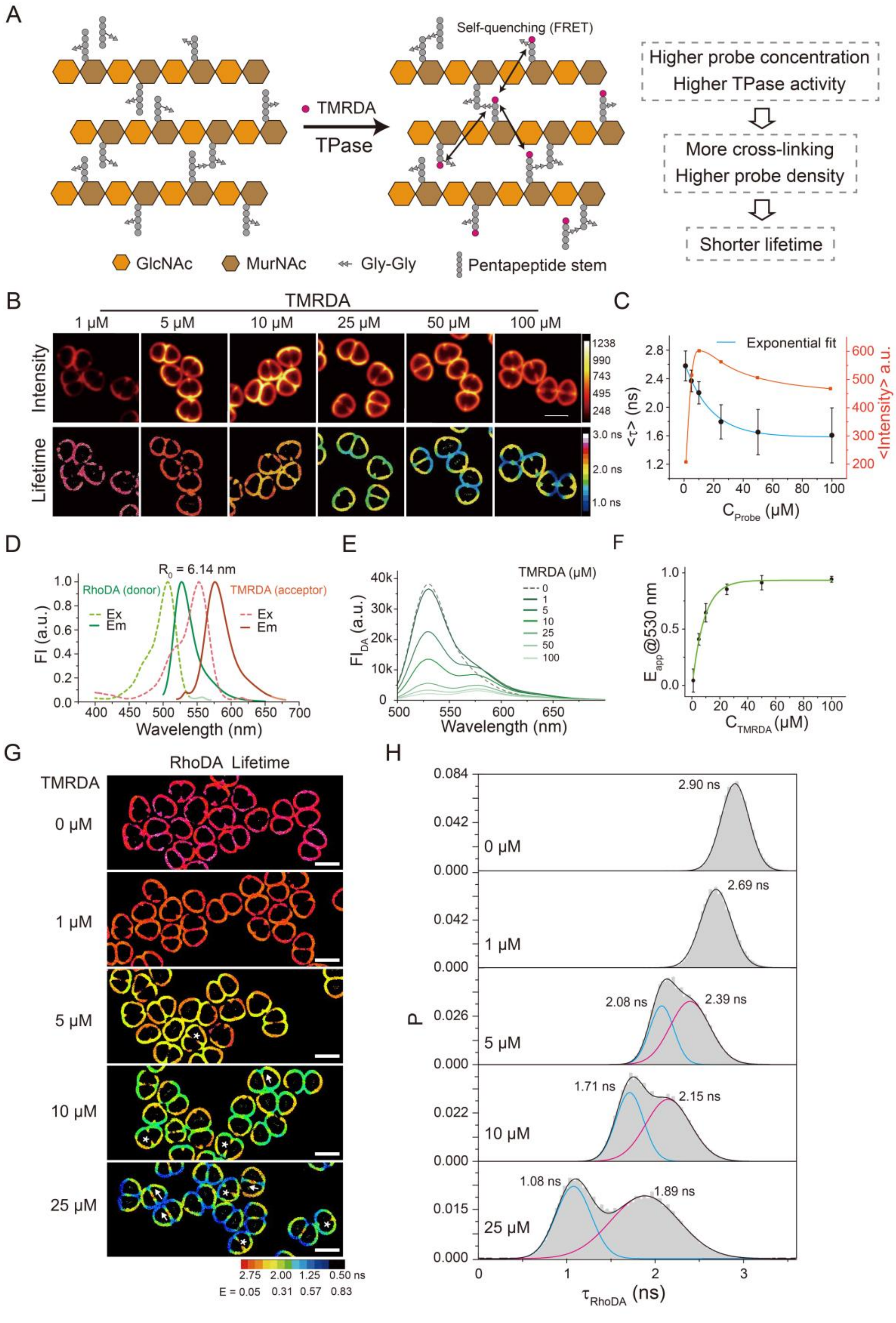

A
Self-quenching (FRET)
TMRDA
TPase
Higher probe concentration
Higher TPase activity
More cross-linking
Higher probe density
Shorter lifetime
GlcNAc
MurNAc
Gly-Gly
Pentapeptide stem
B
TMRDA
1 μM
5 μM
10 μM
25 μM
50 μM
100 μM
Intensity
Lifetime
C
Exponential fit
<τ> (ns)
<Intensity> a.u.
C_Probe (μM)
D
R_0 = 6.14 nm
RhoDA (donor)
TMRDA (acceptor)
Ex
Em
FI (a.u.)
Wavelength (nm)
E
TMRDA (μM)
FI_DA (a.u.)
Wavelength (nm)
F
E_app@530 nm
C_TMRDA (μM)
G
RhoDA Lifetime
TMRDA
0 μM
1 μM
5 μM
10 μM
25 μM
2.75 2.00 1.25 0.50 ns
E = 0.05 0.31 0.57 0.83
H
2.90 ns
2.69 ns
2.08 ns
2.39 ns
1.71 ns
2.15 ns
1.08 ns
1.89 ns
P
τ_RhoDA (ns)

**Figure S1 | Fluorescence lifetime imaging reveals incorporation density of fluorescent D-amino acids (TMRDA and RhoDA) and peptidoglycan cross-linking.**

**(A)** Schematic illustrating how fluorescence lifetime of fluorescent D-amino acid (TMRDA or RhoDA) reports cross-linking of PG. TMRDA or RhoDA is incorporated into the pentapeptide stem of PG by transpeptidases (TPase), which also catalyze the cross-linking reaction between the pentapeptide stems. High TMRDA/RhoDA incorporation therefore indicates increased cross-linking. At elevated labeling density, probes are brought into close proximity, leading to self-quenching or, when an acceptor is co-labeled, Förster Resonance Energy Transfer (FRET), both of which result in shorter fluorescence lifetimes.

**(B)** Representative fluorescence intensity (top) and lifetime (bottom) images of cells labeled with increasing concentrations of TMRDA (1–100 μM, 2 hours). Lifetimes become progressively shorter with higher probe concentrations, reflecting increased labeling density. Above 25 μM, lifetimes also display heterogeneous subcellular distributions, indicating variable TPase activity and different local cross-linking. Scale bar, 2 μm.

**(C)** Quantification of average lifetime (black) and intensity (orange) as a function of probe concentration. Lifetimes decrease exponentially with concentration (cyan), yielding a quenching constant ($k_q$) of 17.4 $\mu M^{-1}$. Intensity increases up to ~10 μM but declines at higher concentrations, consistent with self-quenching at high probe density. Data are mean ± s.e.m. from >100 cells.

**(D)** Excitation (dashed lines) and emission (solid lines) spectra of RhoDA (donor, in green) and TMRDA (acceptor, in red). The Förster distance between the RhoDA-TMRDA pair is 6.14 nm. Ex: excitation; Em: emission; FI: normalized fluorescence intensity.

**(E)** FRET efficiency measured by donor (RhoDA) fluorescence intensity decrease. *Deinococcus radiodurans* (*D. radiodurans*) cells were dual-labeled with 5 μM RhoDA and increasing concentrations (0–100 μM) of TMRDA, and emission spectra were acquired using 480 nm excitation.

**(F)** Apparent FRET efficiency calculated from the fluorescence intensity decay at 530 nm. $E_{app} = \frac{I_0 - I}{I_0}$, where the $I_0$ and $I$ represent the fluorescence intensity of RhoDA at 530 nm in the absence and presence of TMRDA, respectively.

**(G)** Representative fluorescence lifetime images (RhoDA channel) of cells dual-labeled with 5 μM RhoDA and varying TMRDA concentrations (0–25 μM). Increasing TMRDA concentration resulted in reduced donor lifetime and increased FRET efficiency. FRET efficiency values at the leading edge of constricting septa were lower than those at the lagging edge near sidewall (asterisks). In closed septa, FRET efficiency decreased progressively from the sidewall toward the center (arrows). LUT representing the lifetime with FRET efficiency (E) shown at the bottom. Scale bars, 2 μm.

**(H)** Fluorescence lifetime distributions corresponding to **(G).** At low TMRDA concentrations (1–5 μM), a single long-lifetime population of RhoDA was observed. At concentrations above 5 μM, the distributions became bimodal, with reduced lifetimes and increased contribution from the short-lifetime population, indicating a new population with high FRET efficiency. Black curves represent total fits; orange and blue curves show Gaussian fits for the long- and short-lifetime components, respectively. Peak values are indicated.

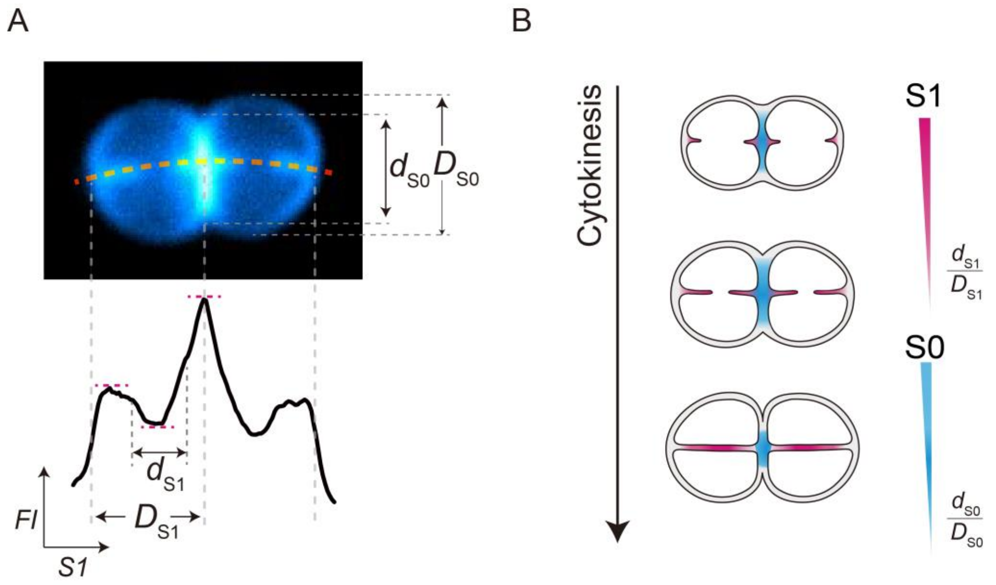


**Figure S2 | Schematic of demograph construction to represent septum formation and development in *D. radiodurans*.**
**(A)** Schematic defining septal morphology in *D. radiodurans*. Diad cells typically contain two sequential septa: constricting septa (S1) and a closed septum (S0). For S1, a curve was drawn along the septum and the intensity profile of membrane labeling was extracted. The full septal length ($\boldsymbol{D}_{\mathbf{S1}}$) was measured between the point of highest contrast at the cell periphery and the maximum signal in the central region (also the center of S0). The gap length ($\boldsymbol{d}_{\mathbf{S1}}$) between the two leading edges was determined from the distance between half-maximum points on either side of the septum. For S0, the cell width ($\boldsymbol{D}_{\mathbf{S0}}$) was obtained from the smallest bounding rectangle of the cell, representing the "full" septal length prior to hydrolysis. The actual length of S0 ($\boldsymbol{d}_{\mathbf{S0}}$) was then measured manually between its two endpoints.
**(B)** Cytokinesis in *D. radiodurans* involves sequential but overlapping septum formation processes: S1 constriction and S0 separation. To capture both, separate demographs were constructed for S1 and S0. During constriction, S1 elongates due to cell growth (increasing $\boldsymbol{D}_{\mathbf{S1}}$) while the leading-edge gap ($\boldsymbol{d}_{\mathbf{S1}}$) narrows; cells were therefore sorted by decreasing $\boldsymbol{d}_{\mathbf{S1}}/\boldsymbol{D}_{\mathbf{S1}}$ ratio to represent constriction progression. During S0 separation, the septum thickens while its length ($\boldsymbol{d}_{\mathbf{S0}}$) shrinks until final separation. Cells were sorted by decreasing $\boldsymbol{d}_{\mathbf{S0}}/\boldsymbol{D}_{\mathbf{S0}}$ ratio. This approach enables demograph-based analysis of protein localization and PG cross-linking throughout cytokinesis.

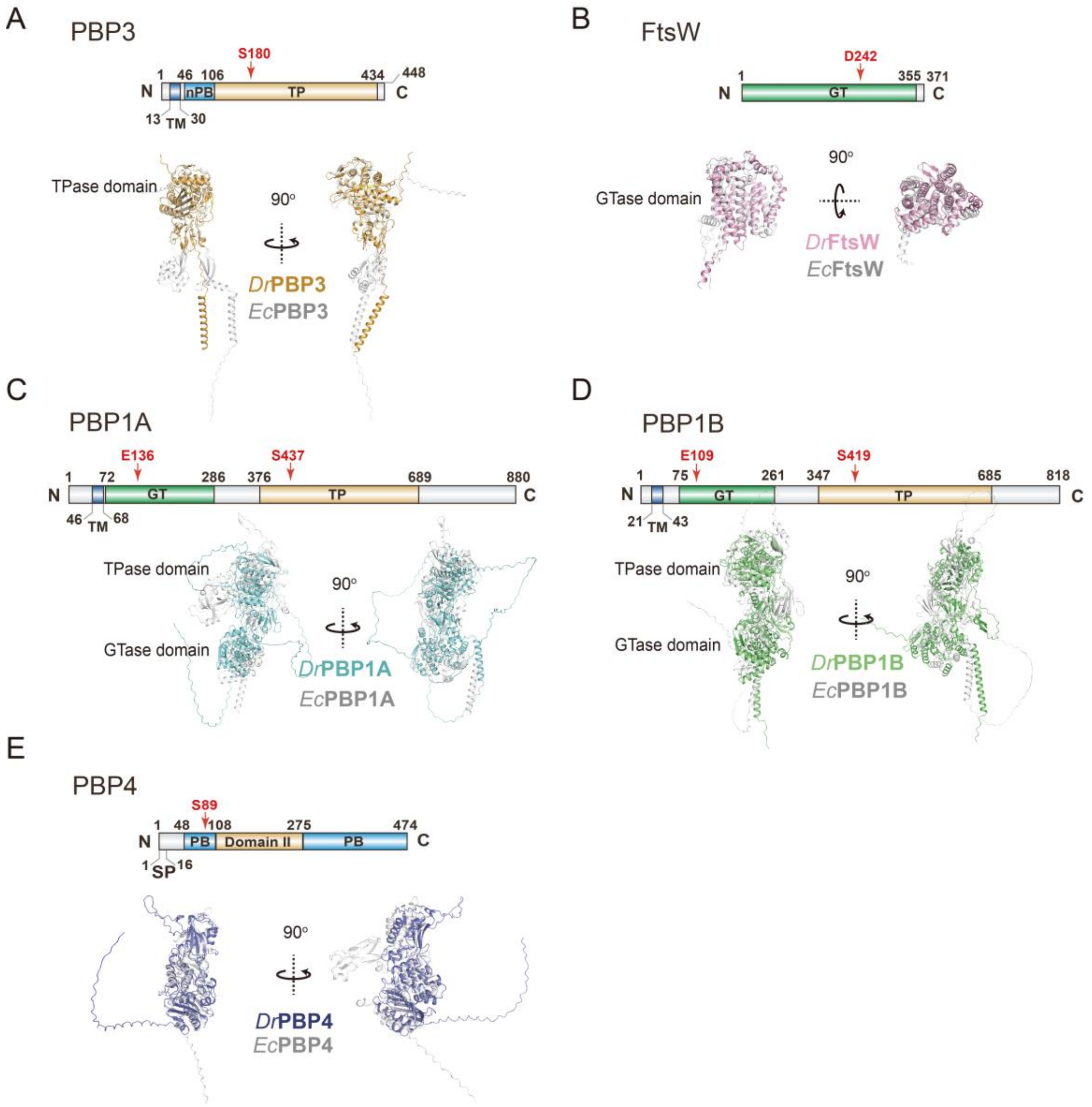


**Figure S3 | Predicted domains and structures of putative SEDS proteins and PBPs in *D. radiodurans*.** Domain architectures and active site annotations of predicted SEDS-family proteins and PBPs in *D. radiodurans* are shown alongside their *E. coli* homologs. PBP1A, PBP1B, PBP3, PBP4, and FtsW were identified by BLAST against the *D. radiodurans* genome using homologous SEDS proteins and PBPs from *Escherichia coli* (*E. coli*), *Bacillus subtilis* (*B. subtilis*) and *Staphylococcus aureus* (*S. aureus*) (see Methods). All the structures were predicted using AlphaFold3[1] and aligned in PyMOL, presenting at two viewing angles. In the models, ***E. coli*** structures are shown in gray, while *D. radiodurans* proteins are colored blue, green, yellow, purple, and pink, respectively.

**(A)** PBP3 possesses a transmembrane helix (TM) domain, a PBP dimerization domain (nPB), and a TPase domain with the catalytic serine located at S180.

**(B**) FtsW contains a glycosyltransferase (GTase) domain with its putative active site at D242 but lacks an associated TPase domain.

**(C**) PBP1A feature a TM domain, a GTase domain, and a TPase domain. The active site residues for DrPBP1A are E136 (GTase) and S437 (TPase). DrPBP1A contain C-terminal intrinsically disordered regions (IDRs).

**(D**) PBP1B feature a TM domain, a GTase domain, and a TPase domain. The active site residues for PBP1B are E109 (GTase) and S419 (TPase). PBP1B contain C-terminal IDRs.

**(E)** PBP4 comprises a signal peptide (SP), a penicillin-binding domain (PB) with an active site at S89, a Domain II homologous to *E. coli* DacB, and a Domain III of unknown function.

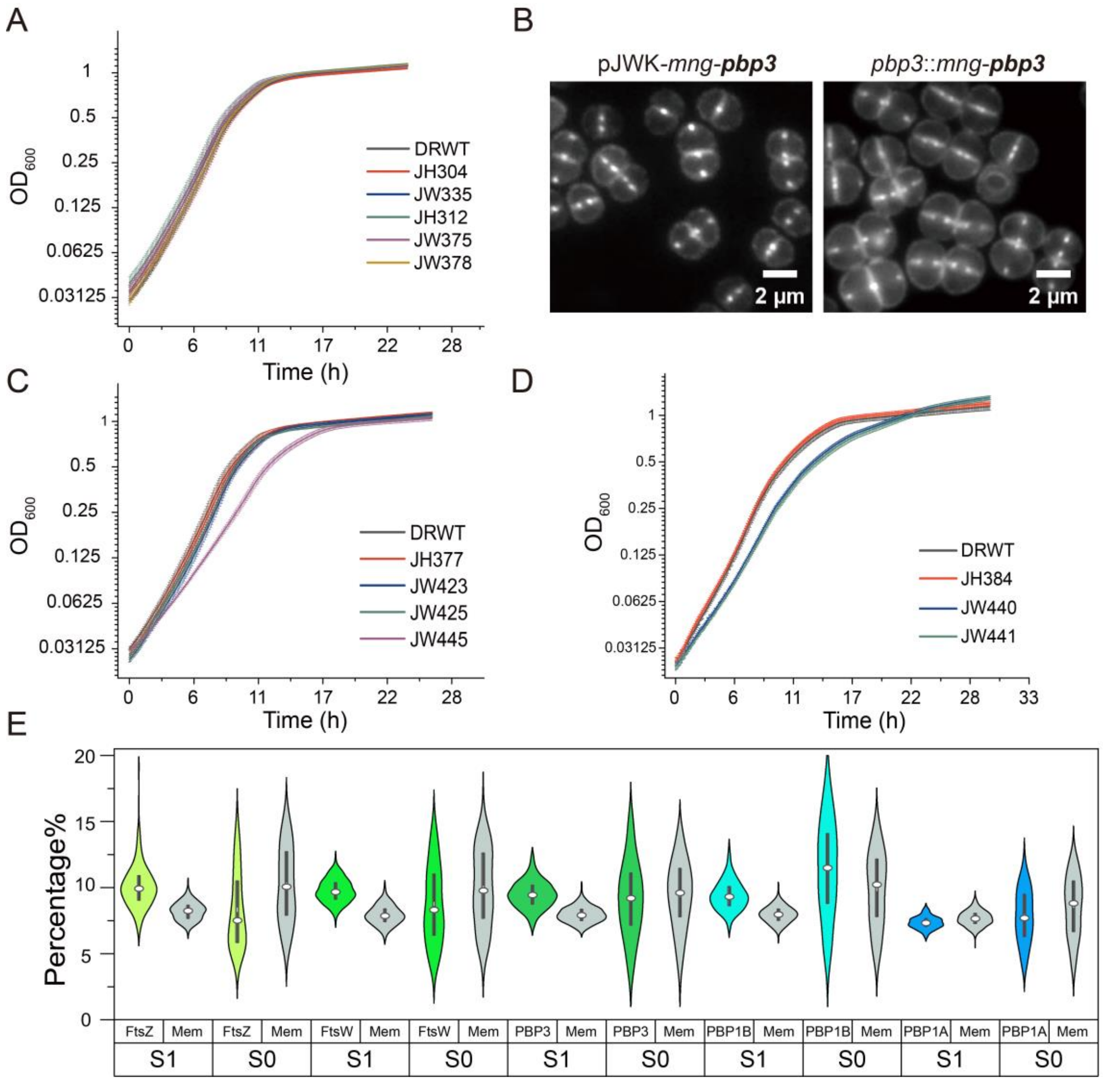


**Figure S4 | Evaluation growth rate, cell morphology, and protein localizations of *D. radiodurans* expressing fluorescently tagged PG synthases.**

**(A)** Growth curves of *D. radiodurans* strains expressing fluorescently tagged FtsZ and PG synthases, including endogenously expressed mNeonGreen[2]-PBP1B (mNG-PBP1B, JW378), FtsW-mNG (JW335) and HaloTag[3]-PBP1A (Halo-PBP1A, JW375), or ectopically expressed mNG-PBP3 (JH312) and FtsZ-mNG (JH304) from plasmids. All strains exhibited comparable growth rates to wild-type *D. radiodurans* (DRWT) in 2×TGY medium at 30 °C. Data represent mean ± s.d. from three independent biological replicates.

**(B)** Representative fluorescence images of cells expressing mNG-PBP3 either from a plasmid or through chromosomal replacement of the native *pbp3* locus. Cells expressing the fusion protein from the chromosome display increased cell size and a higher frequency of forming diads or tetrads, indicating partial loss of function. Scale bars, 2 μm.

**(C)** Growth curves of strains co-expressing mNG- or Halo-tagged PG synthases (FtsW, PBP1A, or PBP1B) and mNG or mCherry (mCh)[4] labeled FtsZ. All strains showed similar growth curves to DRWT, with the exception of JW445 (mNG-PBP1B; FtsZ-mCh), which exhibited a modestly reduced growth rate. JH377, mNG-PBP3, FtsZ-mCh; JW423, Halo-PBP1A, FtsZ-mNG; JW425, FtsW-Halo, FtsZ-mNG. Data represent mean ± s.d. from three independent biological replicates.

**(D)** Growth curves of strains co-expressing mNG-PBP3 with mCh-PBP1B (JH384), mNG-PBP3 with FtsW-Halo (JW440) and mNG-PBP1B with FtsW-Halo (JW441). JW440 and JW441 exhibited a modestly reduced growth rate. Data represent mean ± s.d. from three independent biological replicates.

**(E)** Percentages of the fluorescence intensity of protein of interest (FtsZ, FtsW, PBP3, PBP1B, PBP1A) and the membrane probe (Mem, labeled Potomac Red) at the constricting septum S1 or closed septum S0 over the total intensity of the cell. The circle indicates the median while the line indicates the s.e.m from more than 100 cells.

The genotype of all the strains are listed in Table S1.

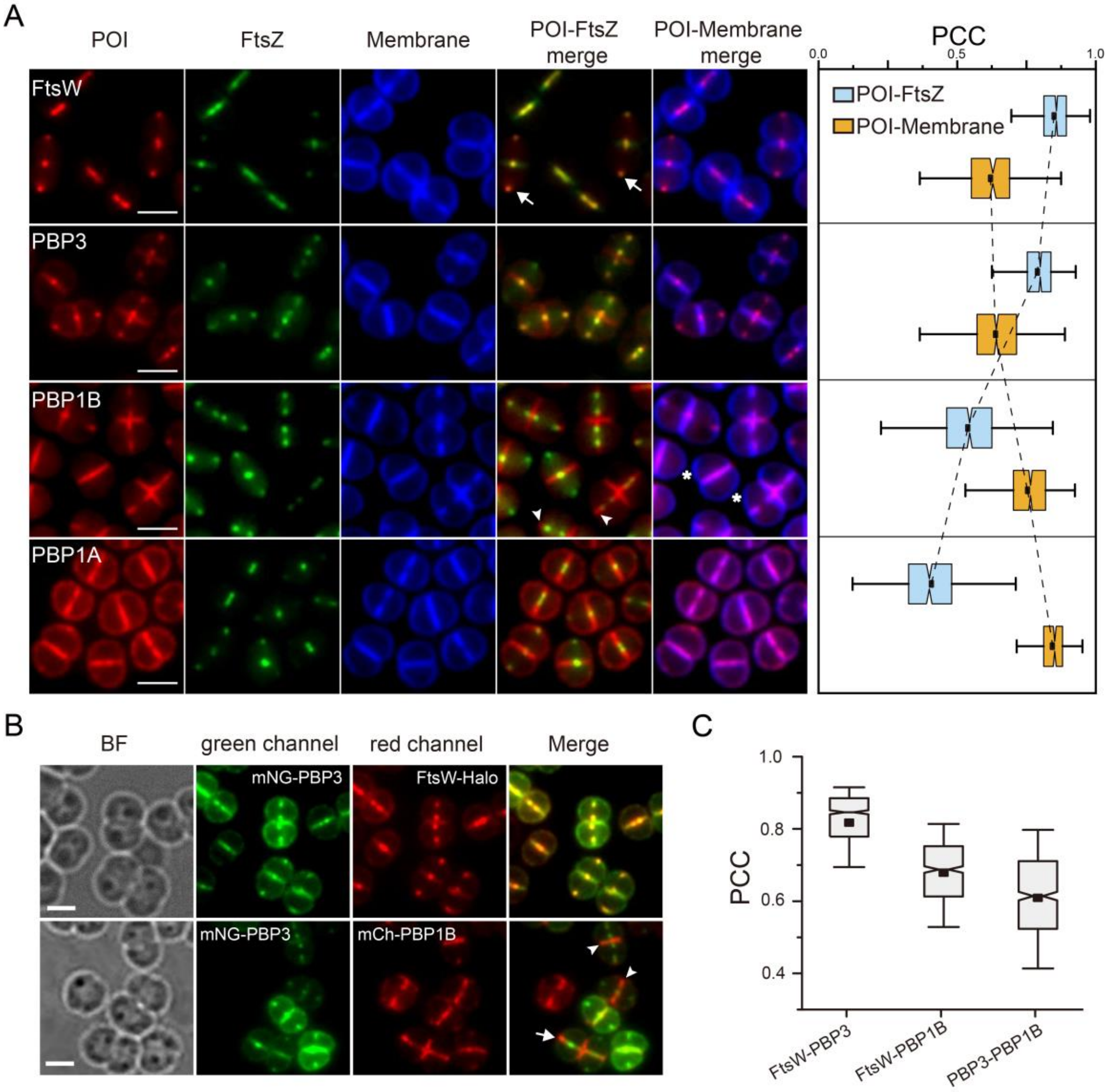


**Figure S5 | Colocalization analysis of PG synthases with FtsZ and membrane in *D. radiodurans*.**

**(A)** Representative fluorescence images showing subcellular localization of proteins of interest (POI) (FtsW, PBP3, PBP1B, and PBP1A; red), FtsZ (green), and membrane (blue). The right panel shows Pearson's correlation coefficients (PCC; see Methods) between each POI and FtsZ (light blue) or membrane (orange). Box plots indicate mean (squares), interquartile ranges (25%-75%, boxes), and 1.5× IQR (whiskers). Arrows, FtsZ and FtsW enrichment at the leading edge of S1; arrowheads, PBP1B accumulation at the lagging edge of S1, outside the FtsZ clusters; asterisks, PBP1B exclusion from peripheral membrane. Scale bars, 2 μm.

**(B)** Representative two-color fluorescence images showing the subcellular distribution of FtsW–PBP3 and PBP3–PBP1B. Top row: Cells co-expressing mNG-PBP1B (green) and FtsW-Halo labeled with JF552[5] (red) (strain JW441). Bottom row: Cells co-expressing mNG-PBP3 (green) and mCh-PBP1B (red) (strain JH384). Arrowheads highlight regions where PBP1B is enriched at S0 with weaker or absent PBP3 signals. Arrows indicate PBP1B accumulation at the lagging edge of S1, while PBP3 localize to the leading edge. Scale bars, 2 μm.

**(C)** Quantification of colocalization between FtsW–PBP3, FtsW–PBP1B, and PBP3–PBP1B using PCC. FtsW and PBP3 show the highest correlation, while PBP1B displays reduced spatial overlap with FtsW and PBP3. Data represent mean ± s.e.m. from more than 600 cells across three independent experiments. Box plots show medians (squares), interquartile ranges (boxes), and whiskers indicating 1.5× IQR.

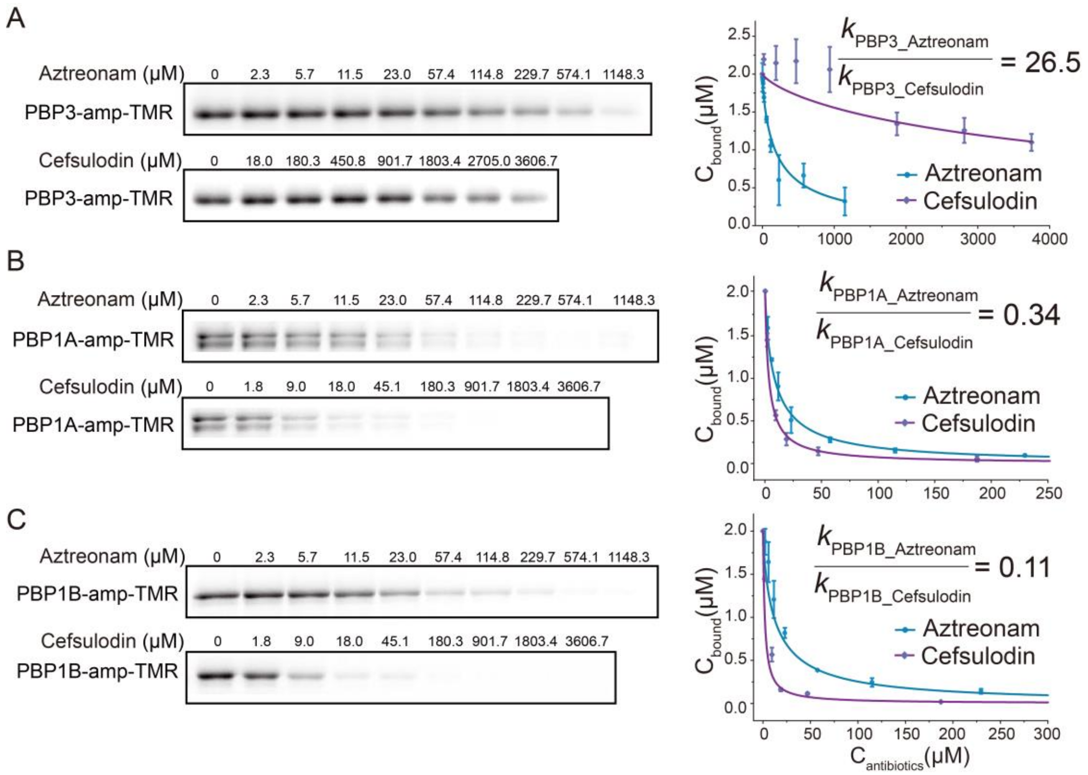


**Figure S6 | Competition assays of aztreonam and cefsulodin with fluorescent ampicillin.**

**(A–C)** End-point competition assays using ampicillin-TMR to evaluate the binding specificity of PBPs in *D. radiodurans*. Left panels show representative SDS-PAGE gels of purified PBP3 **(A)**, PBP1A **(B)**, and PBP1B **(C)** labeled with ampicillin-TMR (2 μM) in the presence of increasing concentrations of aztreonam (top) or cefsulodin (bottom) for 1 hour (amp-TMR indicates ampicillin-TMR). Reactions were terminated with SDS and resolved by SDS-PAGE (see Methods). Right panels are the quantifications of aztreonam- or cefsulodin-mediated inhibition of ampicillin-TMR binding. Blue circles and purple diamonds represent the mean signal intensity corresponding to ampicillin-TMR-labeled PBPs ($C_{bound}$) at different concentrations of antibiotics. Error bars indicate standard deviations from three independent experiments. Binding inhibition curves were fitted to extract the ratios of kinetic rate constants of aztreonam or cefsulodin compared to ampicillin-TMR (see Methods). The ratio of inhibition rates (aztreonam/cefsulodin) for each PBP is indicated. PBP3 exhibited higher affinity for aztreonam, while PBP1B showed stronger inhibition by cefsulodin.

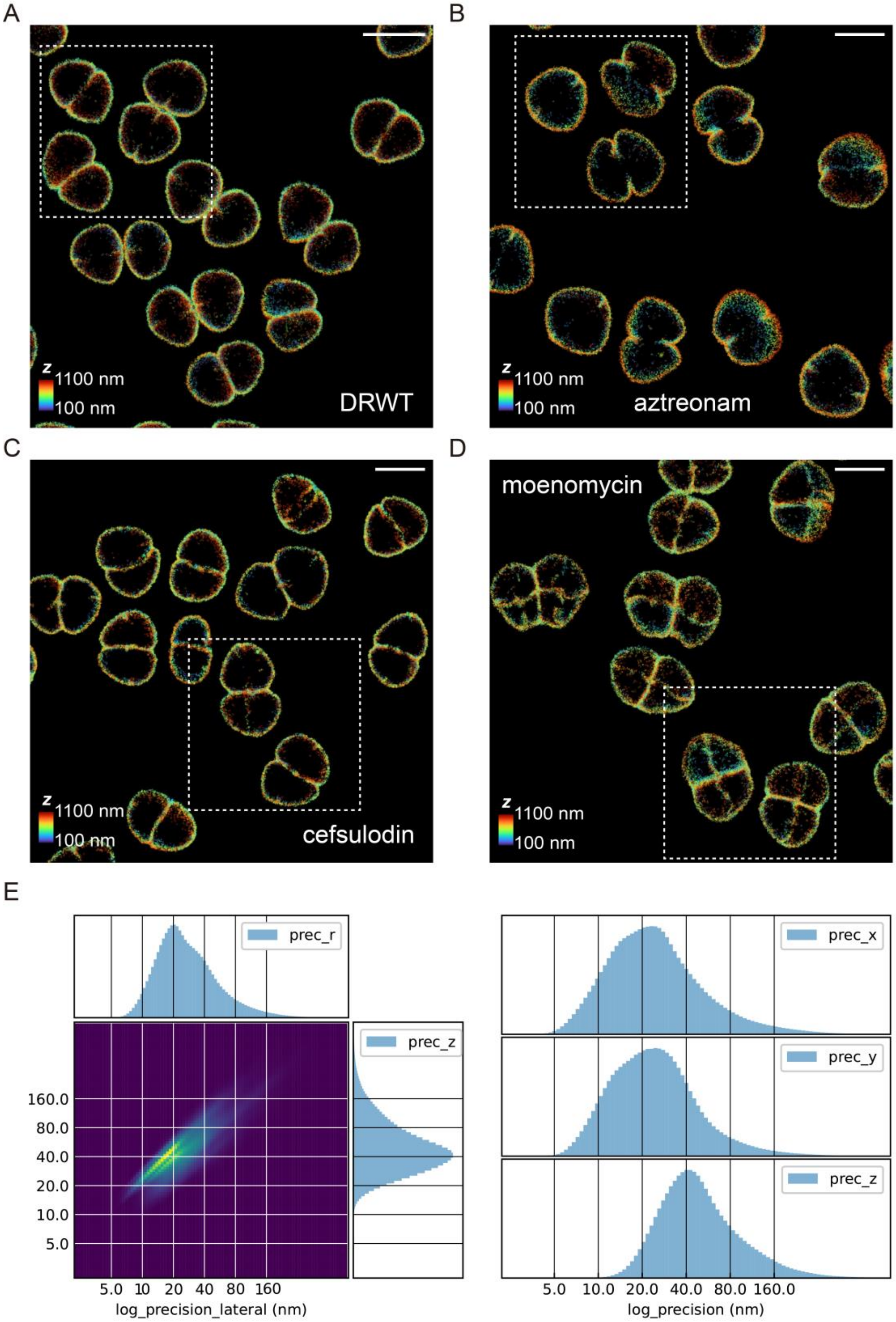


**Figure S7 | Three-dimensional single-molecule localization microscopy (3D-SMLM) images of Cy5DA labeled *D. radiodurans*.**

**(A–D)** Reconstructed 3D-SMLM images of untreated **(A)**, aztreonam-treated **(B)**, cefsulodin-treated **(C)**, and moenomycin-treated **(D)** DRWT cells labeled with Cy5DA. Photon based gauss rendering was used to render the images. The position of localizations along Z axis is demonstrated by color in each image. Rectangles highlight the regions presented in Figure 3A. Scale bars, 2 μm.

**(E)** The distribution of localization precision (uncertainty) of x (proc_x), y (proc_y), x-y (prec_r) and z (prec_z) estimated by the Cramer-Rao lower bound using our previously reported method[6].

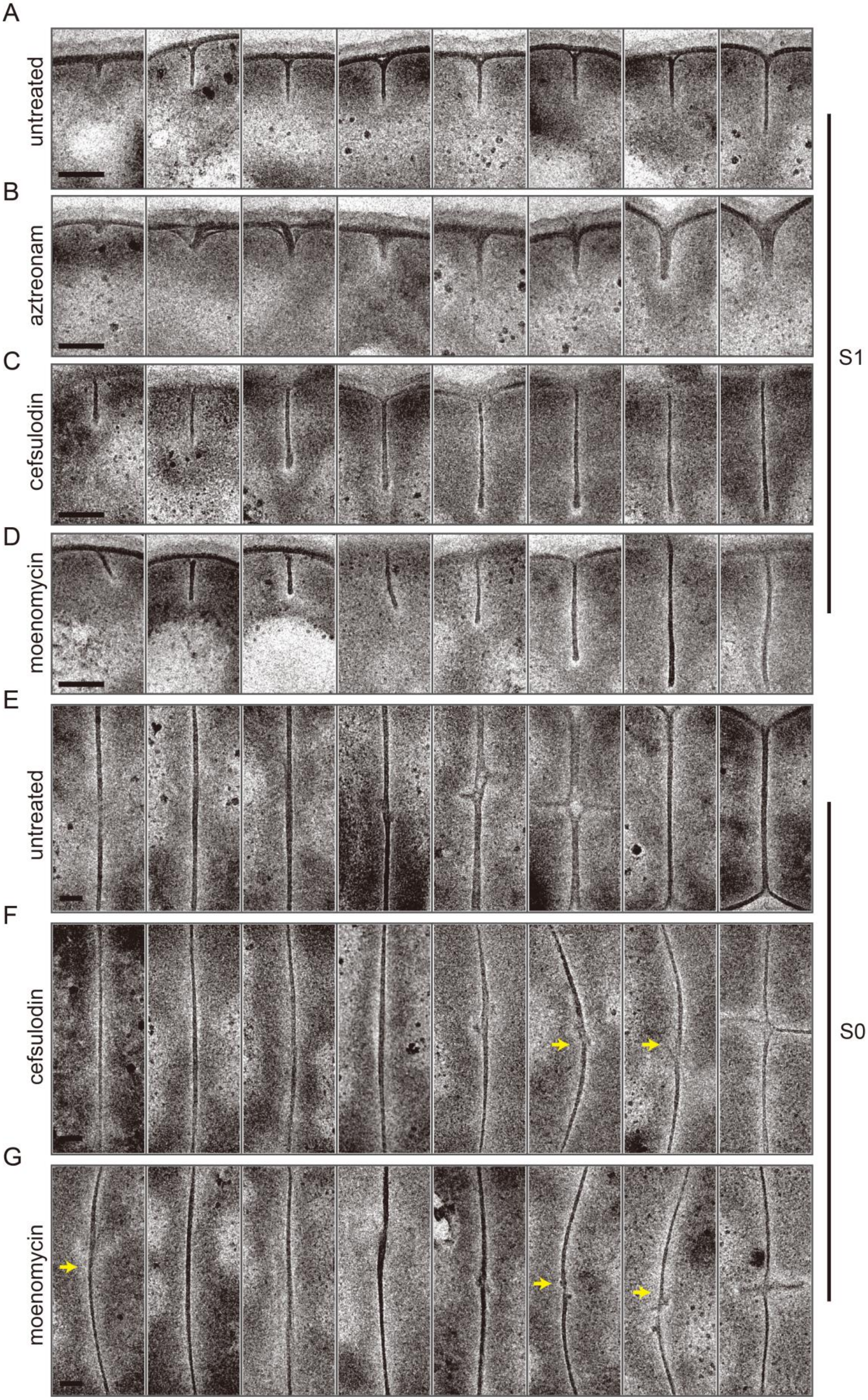

A
untreated
B
aztreonam
C
cefsulodin
D
moenomycin
E
untreated
F
cefsulodin
G
moenomycin
S1
S0

**Figure S8 | Additional electron microscope images of S1 and S0 septa.**
**(A–D)** Representative electron tomographic slices showing S1 septa from DRWT cells under different conditions: untreated control (**A**), aztreonam-treated (**B**), cefsulodin-treated (**C**), and moenomycin-treated (**D**). In aztreonam-treated cells, S1 septa often exhibited thickened, triangular shapes. In contrast, cells treated with cefsulodin or moenomycin showed noticeably thinner or absence of bases of the S1 septa compared to untreated cells, although the leading edge of the septum remained relatively unaffected. All images were selected from the focused slice from tomographic tilt series of individual cells. Scale bars, 100 nm.
**(E–G)** Representative electron tomographic slices showing S0 septa from DRWT cells under different conditions: untreated control (**E**), cefsulodin-treated (**F**), and moenomycin-treated (**G**). Compared to untreated cells, both cefsulodin and moenomycin treatments led to visibly thinner, with some appearing curved or irregular structure (arrows). All images were selected from the focused slice from tomographic tilt series of individual cells. Scale bars, 100 nm.

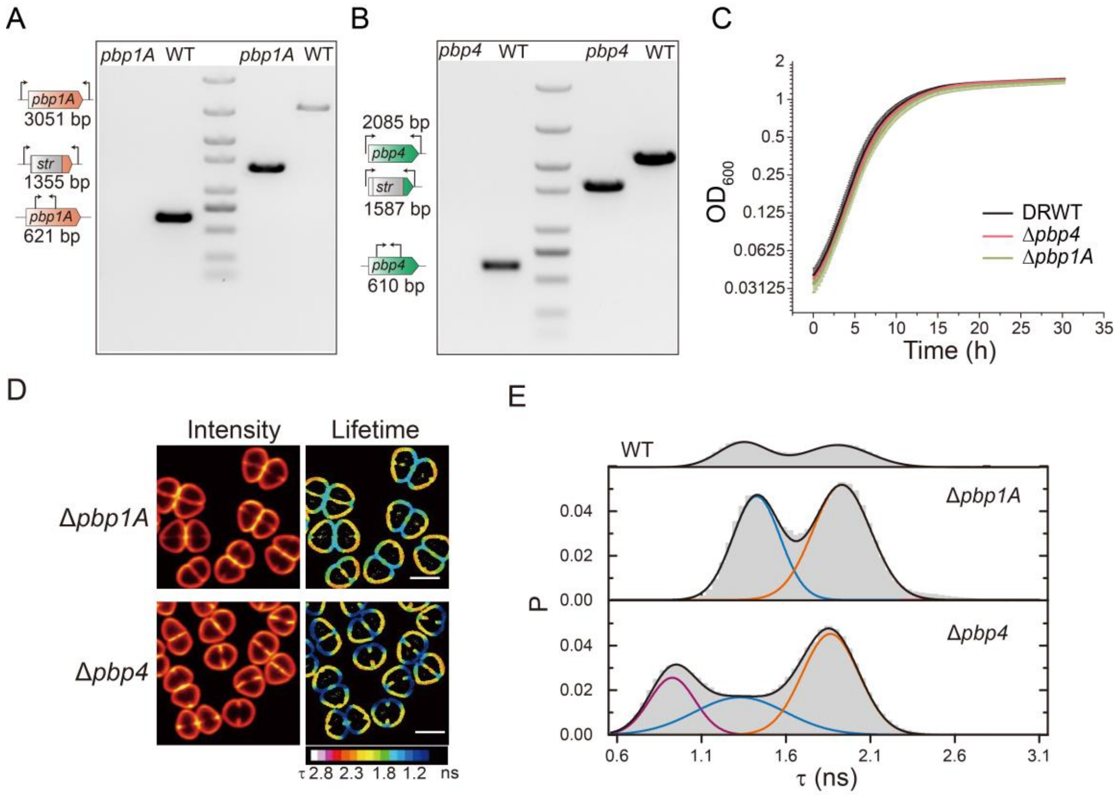


**Figure S9 | Effects of *pbp1A* and *pbp4* knockout on cell growth, morphology, and septal PG cross-linking in *D. radiodurans*.**
**(A–B)** PCR validation of *pbp1A* and *pbp4* knockout strains. Genomic DNAs from knockout and DRWT cells were used as templates for PCR amplification with inside and outside primers (Table S2). Expected band sizes and loci are indicated. All products were verified by Sanger sequencing. Gel contrast was adjusted in ImageJ[7] for visualization.
**(C)** Growth curves of *pbp1A* and *pbp4* knockout strains in 2×TGY medium at 30 °C. Data represent mean ± s.d. from three independent biological replicates.
**(D)** Representative fluorescence intensity (left) and fluorescence lifetime (right) images of TMRDA-labeled cells. Lifetime maps use a rainbow LUT (0.7–2.8 ns). Scale bars, 2 μm.
**(E)** Fluorescence lifetime distributions corresponding to **(D)**, from three independent biological replicates. Data were fitted with two or three Gaussian populations. Deletion of *pbp1A* produced no detectable change compared to DRWT (top), whereas deletion of *pbp4* introduced an additional short-lifetime population (~0.8 ns)

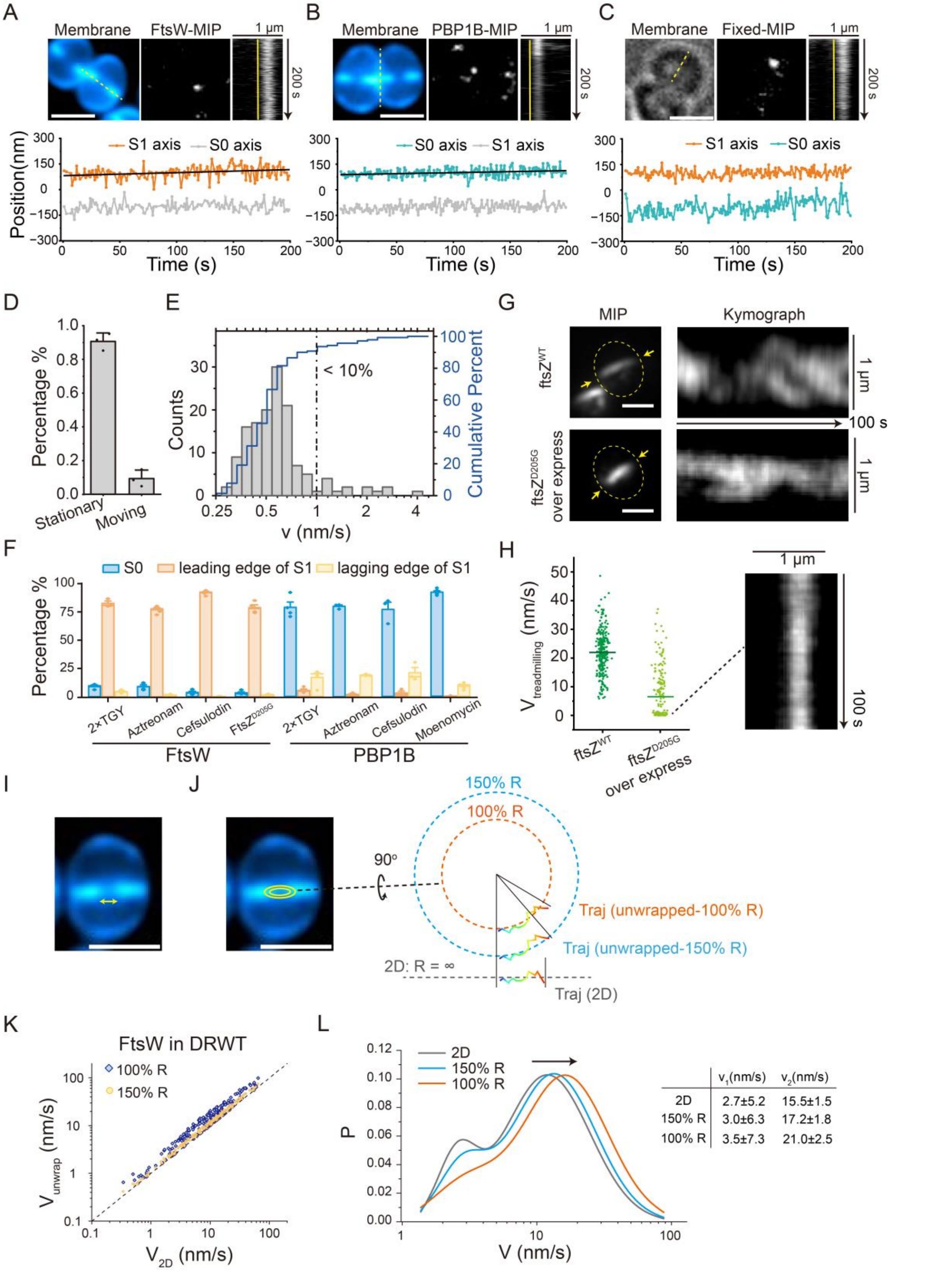


**Figure S10 | Additional examples and analyses of FtsW, PBP1B, and FtsZ dynamics.**

**(A–B)** Representative trajectories of stationary FtsW-Halo (**A**, at S1) and Halo-PBP1B (**B**, at S0) labeled with JFX650[8]. Top panels: membrane staining (left), maximum intensity projection (MIP, middle), and kymographs of septal line scans (right; scan positions marked in membrane images). Bottom panels: extracted 1D trajectories along S1 or S0. Scale bars, 2 μm.

**(C)** A representative trajectory of FtsW-Halo in paraformaldehyde (PFA)-fixed cells. Scale bar, 2 μm.
**(D)** Mobility classification of FtsW molecules in fixed cells. ~90% were stationary using our established method[9]. Data are mean ± s.d. from three independent experiments.
**(E)** Cumulative probability distribution of directional FtsW speeds in fixed cells. Fewer than 10% moved faster than 1 nm/s, confirming that motile populations in live cells (fewer than 1% with speed faster than 1 nm/s) are not fixation artifacts.
**(F)** Spatial distribution of FtsW and PBP1B trajectories under perturbations. In aztreonam-, cefsulodin-treated, and FtsZ$^{D205G}$-overexpressing cells, ~75% of FtsW trajectories localized to the leading edge of S1. In aztreonam-, cefsulodin-, and moenomycin-treated cells, ~75% of PBP1B trajectories remained enriched at S0, with ~10% at the lagging edge of S1. Data are mean ± s.d. from three to five replicates.
**(G)** FtsZ-mNG treadmilling in DRWT (top) and FtsZ$^{D205G}$-overexpressing (bottom) cells. Left: MIP images overlaid with outlines (dashed circles) and kymograph scan positions (arrows). Right: kymographs showing treadmilling dynamics.
**(H)** FtsZ treadmilling speed distributions. Overexpression of FtsZ$^{D205G}$ reduced mean speed from 22.0 nm/s (dark green) to 6.5 nm/s (light green). Many mutant cells displayed stalled FtsZ clusters; an example kymograph is shown (right).
**(I–L)** Correction of apparent FtsW speed underestimation due to 3D-to-2D projection. When septa are highly constricted, FtsW molecules move along the constricting ring (enclosed by the leading edge of S1, arrow marked region in **(I)**), but trajectories are recorded only in 2D projections of this circular path **(J)**. This projection compresses arc length into a shorter linear distance, underestimating the true velocity. To correct this bias, we estimated the radius of the ring from membrane staining and recalculated FtsW speeds assuming circular motion ("unwrapping the trajectories"), also testing a 150% wider radius to account for errors introduced by limited spatial resolution **(J)**. After correction, the velocity of individual FtsW molecules increased **(K)** and the fast-moving population shifted to ~21.0 nm/s **(L)**, closely matching the FtsZ treadmilling speed (22.0 nm/s) measured by TIRF microscopy in less constricted septa **(H)**. This analysis supports that the fast-moving fraction of FtsW is coupled to FtsZ treadmilling. Scale bars, 2 μm.

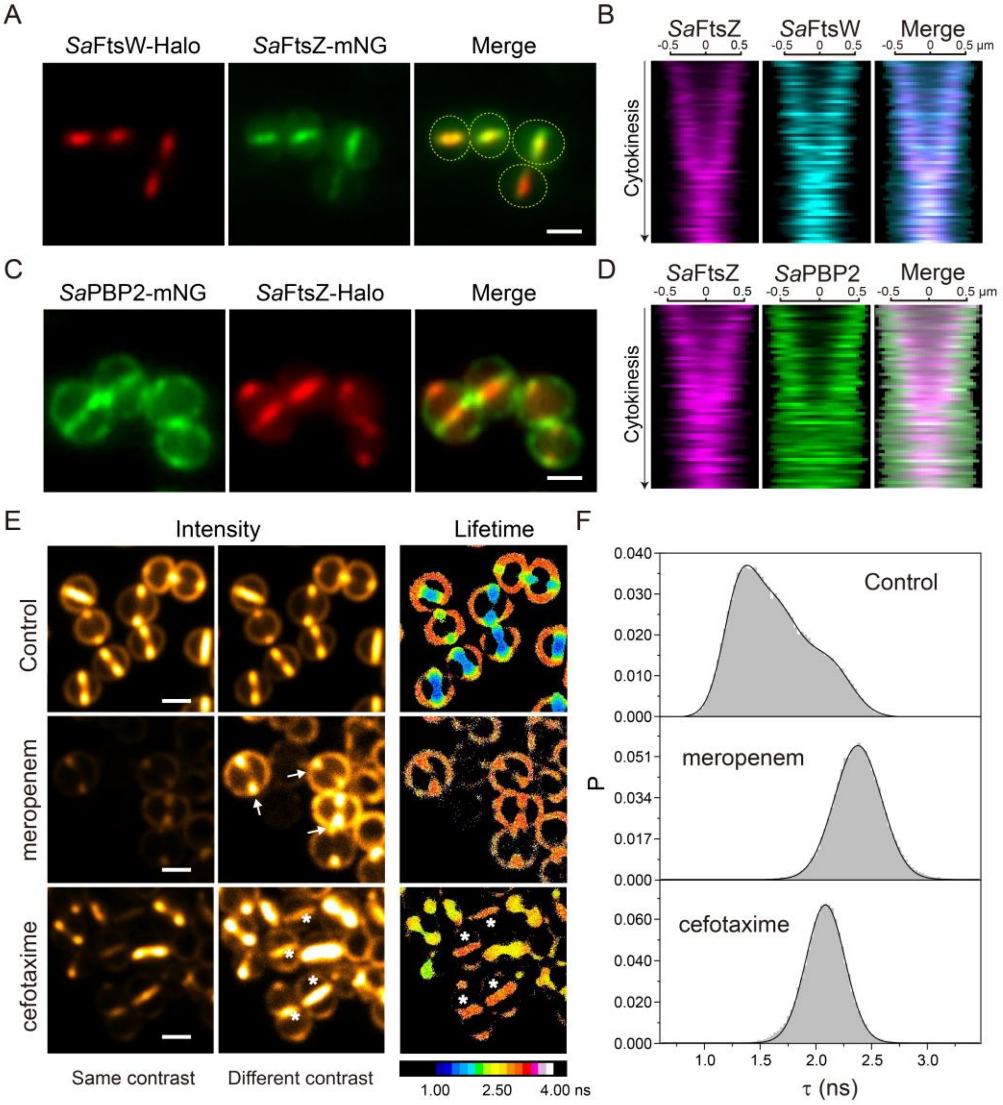


**Figure S11 | Spatial organization and functional roles of *Sa*FtsW–*Sa*PBP1 and *Sa*PBP2 in septal PG synthesis of *S. aureus*.**
(**A**) Representative fluorescence images of ***S. aureus*** cells co-expressing *Sa*FtsW–Halo (JF552, red) and *Sa*FtsZ–mNG (green). *Sa*FtsW colocalizes to the same regions at the septum as *Sa*FtsZ. Dashed curves mark cell boundaries. Scale bar, 1 μm.
(**B**) Demograph analysis of *Sa*FtsZ (magenta) and *Sa*FtsW (cyan) signals along the septal axis across the constriction process. Septa were aligned and sorted by the distance between *Sa*FtsZ intensity peaks (Methods). Merged demographs show consistent overlap of *Sa*FtsW with *Sa*FtsZ throughout septal ingrowth.
(**C**) Representative fluorescence images of cells co-expressing mNG–*Sa*PBP2 (green) and *Sa*FtsZ–Halo (JF552, red). *Sa*PBP2 is broadly distributed on the envelope, enriched at septa but positioned behind FtsZ. Scale bar, 1 μm.
(**D**) Demograph analysis of *Sa*FtsZ (magenta) and *Sa*PBP2 (green), aligned and sorted as in (**B**). The merged demograph shows that *Sa*PBP2 follows the constriction trajectory of FtsZ but is spatially displaced toward the lagging edge of the septum.
(**E**) Representative fluorescence images of *S. aureus* cells labeled with RhoDA under selective antibiotic treatments. First column: fluorescence intensity images (same contrast). Second column: intensity images with enhanced contrast. Third column: fluorescence lifetime maps with a rainbow LUT (1.0–4.0 ns). In untreated cells, septal centers display short lifetimes, consistent with active PG synthesis. Meropenem (*Sa*PBP1 TPase inhibitor[10,11]) blocked septal ingrowth, leaving many unclosed, low–cross-linked septa (arrows). Cefotaxime (*Sa*PBP2 TPase inhibitor[10,12]) allowed septal closure (asterisks) but yielded uniformly long lifetimes across the septum, indicating reduced cross-linking. Scale bars, 1 μm.
(**F**) Fluorescence lifetime distributions of septa under the conditions shown in (**E**).

**Supplementary Table 1: Strains and plasmids.**

| Bacteria | Strain | Genotype | Source |
|---|---|---|---|
| *D. radiodurans* | R1 | *Deinococcus radiodurans* wild-type ATCC 13939 | A gift from Dr. Yuejing Hua[13] |
| *D. radiodurans* | JW116 | R1, Δ*dr_0176 (pbp4)::str* | This study |
| *D. radiodurans* | JW117 | R1, Δ*dr_0479 (pbp1A)::str* | This study |
| *D. radiodurans* | JW335 | R1, *dr_2497 (ftsW)::ftsW-linker0-mng str* | This study[a] |
| *D. radiodurans* | JW337 | R1, *ftsW::ftsW-linker0-halo str* | This study |
| *D. radiodurans* | JW375 | R1, *pbp1A::N1-halo-linker0-pbp1A str* | This study |
| *D. radiodurans* | JW378 | R1, *pbp1B::N-mng-linker0-pbp1B str* | This study |
| *D. radiodurans* | JW423 | R1, *pbp1A::N1-halo-linker0-pbp1A str*, pJWK-$P_{tetdr}$*-ftsZ-linker0-mng* | This study |
| *D. radiodurans* | JW425 | R1, *ftsW::ftsW-linker0-halo str*, pJWK-$P_{tetdr}$*-ftsZ-linker0-mng* | This study |
| *D. radiodurans* | JW440 | R1, *ftsW::ftsW-linker0-halo str*, pJWK-$P_{tetdr}$*-N-mng-linker0-pbp3* | This study |
| *D. radiodurans* | JW441 | R1, *ftsW::ftsW-linker0-halo str*, pJWK-$P_{tetdr}$*-N-mng-linker0-pbp1B* | This study |
| *D. radiodurans* | JW443 | R1, *pbp1A::N1-halo-linker0-pbp1A str*, pJWK-$P_{tetdr}$*-N-mng-linker0-pbp3* | This study |
| *D. radiodurans* | JW445 | R1, *pbp1B::N-mng-linker0-pbp1B str*, pJWK-$P_{tetdr}$*-ftsZ-linker0-mch* | This study |
| *D. radiodurans* | JW449 | R1, pJWK-$P_{spac1}$*-N-halo-pbp1B* | This study |
| *D. radiodurans* | JH304 | R1, pJWK-$P_{tetdr}$*-ftsZ-linker0-mng* | This study |
| *D. radiodurans* | JH312 | R1, pJWK-$P_{tetdr}$*-mng-linker0-pbp3* | This study |
| *D. radiodurans* | JH377 | R1, pJWK-$P_{tetdr}$*-N-mng-pbp3-ftsZ-linker0-mch* | This study |
| *D. radiodurans* | JH379 | R1, *ftsW::ftsW-linker0-halo str*, pJWK-$P_{tetdr}$-$ftsZ^{D205G}$*-linker0-mng* | This study |
| *D. radiodurans* | JH384 | R1, *pbp1B::N-mch-linker0-pbp1B str*, pJWK-$P_{tetdr}$*-N-mng-linker0-pbp3* | This study |
| *D. radiodurans* | JH385 | R1, *pbp1B::N-mng-linker0-pbp1B str*, pJWK-*N1-mch-linker0-pbp1A* | This study |
| *D. radiodurans* | JH386 | R1, *ftsW::ftsW-linker0-mng str*, pJWK-*N1-mch-linker0-pbp1A* | This study |
| *E. coli* | DH5α | supE44 **Δ**lacU(Φ80lacZ1 M15) hsdR17 recA1 endA1 gyrA96 thi-1 relA1 | Laboratory stock |
| *E. coli* | BL21 | F– ompT hsdSB (rB– mB–) gal dcm | Laboratory stock |
| *S. aureus* | RN4220 | Restriction-deficient derivativeof NCTC8325-4 | Laboratory stock |
| *S. aureus* | JW460 | RN4220, pCN5-*N-mng-linker1-sapbp2*-$saftsZ^{55\text{-}56}$*-halo* | This study[b] |
| *S. aureus* | JW461 | RN4220, pCN5-*saftsW-linker2-halo*-$saftsZ^{55\text{-}56}$*-mng* | This study[c] |
| *S. aureus* | JW462 | RN4220, pCN5-*N-mng-linker1-sapbp2-saftsW-linker2-halo* | This study |

| Plasmid | Genotype | Source |
|---|---|---|
| pRADK | ColE1 (*E. coli*), DrOri (*D. radiodurans*), *bla* (*E. coli*), *cat* (*D. radiodurans*), *repU*, $P_{lac}$*, AT-rich box, *orfB******, *orfH******, $P_{groes}$::*aph* | Laboratory stock |
| pCNX | ColE1 (*E. coli*), pT181cop-wt repC (*S. aureus*), *bla* (*E. coli*), *aph* (*S. aureus*), *LacI*, $P_{cad\text{-}cadC}$:: | A gift from Dr. Mariana G. Pinho[14] |
| pCN1 | ColE1 (*E. coli*), pT181cop-wt repC (*S. aureus*), *bla* (*E. coli*), *cat* (*S. aureus*), *LacI*, $P_{cad\text{-}cadC}$:: | Laboratory stock |
| pCN5 | ColE1 (*E. coli*), pT181cop-wt repC (*S. aureus*), *bla* (*E. coli*), *cat* (*S. aureus*), *LacI*, $P_{spac}$:: | Laboratory stock |
| pXY027 | $P_{T5lac}$::*ftsZ-gfpmut2* | Laboratory stock[15] |
| pXW001 | $P_{tet}$::$hpccma^{I55A}$*-gph-mng* | Laboratory stock |
| pJH024 | $P_{tet}$::*hfq-gpg-halo* | Laboratory stock |

| | | |
|---|---|---|
| pJH032 | $P_{tet}$::*mch-ggs-hfq* | Laboratory stock |
| pJW18 | pRADK, $P_{groes}$::*gfpmut2* | This study |
| pJW31 | pJW18, $P_{1261}$::*gfpmut2* | This study |
| pJW32 | ColE1 (*E. coli*), DrOri (*D. radiodurans*), *cat*, *repU*, $P_{lac}$*, AT-rich box, $P_{1261}$::*gfp* | This study |
| pJW33 | ColE1 (*E. coli*), DrOri (*D. radiodurans*), *cat*, *repU*, AT-rich box*, $P_{1261}$::*gfp* | This study |
| pJWK | ColE1 (*E. coli*), DrOri (*D. radiodurans*), *aph*, *repU*, AT-rich box*, $P_{1261}$::*gfp* | This study |
| pJWK1 | pJWK, $P_{1261}$::*mch* | This study |
| pJWK7 | pJWK, $P_{1261}$::*N1-mch-linker0-pbp1A* | This study |
| pJWK64 | ColE1 (*E. coli*), DrOri (*D. radiodurans*), *aph*, *repU*, AT-rich box*, *LacI*, $P_{spac1}$::*mch* | This study |
| pJWK65 | ColE1 (*E. coli*), DrOri (*D. radiodurans*), *aph*, *repU*, AT-rich box*, *LacI*, $P_{spac2}$::*mch* | This study |
| pJWK67 | ColE1 (*E. coli*), DrOri (*D. radiodurans*), *aph*, *repU*, AT-rich box*, *TetR*, $P_{tetdr}$::*mch* | This study |
| pJWK81 | pJWK67, $P_{tetdr}$::*N-mng-linker0-pbp1B* | This study |
| pJWK84 | pJWK67, $P_{tetdr}$::*ftsZ-linker0-mch* | This study |
| pJWK87 | pJWK64, $P_{spac1}$::*N-halo-pbp1B* | This study |
| pJWK88 | pCN5, $P_{spac}$::*N-mng-linker1-sapbp2-saftsZ*$^{55\text{-}56}$*-halo* | This study |
| pJWK89 | pCN5, $P_{spac}$::sa*ftsW-linker2-halo-saftsZ*$^{55\text{-}56}$*-mng* | This study |
| pJWK90 | pCN5, $P_{spac}$::*N-mNG-linker1-sapbp2-saftsW-linker2-halo* | This study |
| pJH120 | pJWK, $P_{1261}$::*ftsZ-linker0-mng* | This study |
| pJH150 | pJWK67, $P_{tetdr}$::*ftsZ-linker0-mng* | This study |
| pJH158 | pJWK67, $P_{tetdr}$::*mng-linker0-pbp3* | This study |
| pJH180 | pJWK67, $P_{tetdr}$::*ftsZ*$^{D205G}$*-linker0-mng* | This study |
| pJH195 | pJWK67, $P_{tetdr}$::*N-mng-pbp3-ftsZ-linker0-mch* | This study |
| pCAG-Hrd1-CS | *SV40* (*mammalian*), pMB1(*E. coli*), *bla* (*E. coli*), $P_{CAG}$::*hrd1-Twin-Strep-tag* | A gift from Dr. Hongwu Qian[16] |
| pYY302 | pET-28a(+), $P_{T7}$::*ferritn-mng* | Laboratory stock |
| pJH130 | pET-28a(+), $P_{T7}$::*pbp1A*$^{\Delta 1\text{-}73}$*-His*$_{10}$*-Twin-Strep-tag* | This study |
| pJH131 | pET-28a(+), $P_{T7}$::*pbp1B*$^{\Delta 1\text{-}48}$*-His*$_{10}$*-Twin-Strep-tag* | This study |
| pJH132 | pET-28a(+), $P_{T7}$::*pbp3*$^{\Delta 1\text{-}35}$*-His*$_{10}$*-Twin-Strep-tag* | This study |

Note:
a. DNA sequence of linker0: ggtggcggtggtagcccggcaccggcaccgggcggtggcggtagc
Peptide sequence of linker0: GGGGSPAPAPGGGGS
b. DNA sequence of linker1: ggcggtggtgggtctggtggcggcggatct
Peptide sequence of linker1: GGGGSGGGGS
c. DNA sequence of linker2: ttggaaggatcaggacaaggaccaggatctggtcaaggttctggt
Peptide sequence of linker2: LEGSGQGPGSGQGSG

**Supplementary Table 2: DNA Oligos.**

| Primer | Sequence[a] | Purpose |
|---|---|---|
| Primer001 | ACTGTCTTCCAAGCTTGATATCGAATTCGAGCTCG | Amplification of *aph* for JW116 |
| Primer002 | ACAATTCATCGGATCCTTATTTGCCGACTACCTT | |
| Primer003 | CTTCCGTGAAACCGGCAC | Amplification of the upstream homologous arm of *pbp4* for JW116 |
| Primer004 | TATCAAGCTTGGAAGACAGTTTAAGGCGTCCCGCC | |
| Primer005 | ATAAGGATCCGATGAATTGTGATTGACGATGGAGG | Amplification of the downstream homologous arm of *pbp4* for JW116 |
| Primer006 | GCTCCCAGCAGGAGGTG | |
| Primer007 | CACCATGAAACTGGTCACGG | Knockout strain JW116 validation |
| Primer008 | CGACCTTTCTCGTCCTGAC | |
| Primer009 | CTGTGCAGGTAAGCTTGATATCGAATTCGAGCTCG | Amplification of *aph* for JW117 |
| Primer010 | GGCGGCAGCGGGATCCTTATTTGCCGACTACCTTG | |
| Primer011 | GTCGGGTTGAGCAGTGTTG | Amplification of upstream homologous arm of *pbp1A* for JW117 |
| Primer012 | TATCAAGCTTACCTGCACAGCATAGACGGCGAAT | |
| Primer013 | ATAAGGATCCCGCTGCCGCCGAGGCCATAAGT | Amplification of downstream homologous arm of *pbp1A* for JW117 |
| Primer014 | CATCCGCTCGGAGGTGG | |
| Primer015 | ACCGGCGCTTTTACACCCACA | Knockout strain JW117 validation |
| Primer016 | ATTTGCCCGTCTTCTACCATGC | |
| Primer017 | GCTGGGAATACTTCGCGG | Amplification of *ftsw*$^{698-1113}$ for JW335 |
| Primer018 | GCCCGGTGCCGGTGCCGGGCTACCACCGCCACCGTCGT CCGCCGCGGC | |
| Primer019 | AGCCCGGCACCGGCACCGGGCGGTGGCGGTAGCGTGA GCAAAGGCGAAGAAG | Amplification of *mng* for JW335 |
| Primer020 | ATAAGGATCCTCATTTATACAGTTCATCCATGCCC | |

| | | |
|---|---|---|
| Primer021 | ACTGTATAAATGAGGATCCTTATTTGCCGACTA | Amplification of *aph* for JW335 |
| Primer022 | CTATTCCGGGAAGCTTGATATCGAATTCGAG | |
| Primer023 | TATCAAGCTTCCCGGAATAGGAAGAAAAGA | Amplification of downstream homologous arm of *ftsw* for JW335 |
| Primer024 | GTCAACGACGTCCACGG | |
| Primer025 | TGCTCAATCATCCGGTAATCCCCGA | Amplification of upstream homologous arm of *pbp1A* for JW375 |
| Primer026 | CCAGTACCGATTTCGCTACCCATGGATCCTTATTTGCCGACTAC | Amplification downstream homologous arm of *aph* for JW375 |
| Primer027 | ATGGGTAGCGAAATCGGTACTGGCTTTCC | Amplification of *halo* for JW375 |
| Primer028 | GCCCGGTGCCGGTGCCGGGCTACCACCGCCACCACCGGAAATCTCCAGAGTAG | |
| Primer029 | GCCCGGCACCGGCACCGGGCGGTGGCGGTAGCTCGCGCTCGTCCCAGG | Amplification of *pbp1A*$^{1-382}$ for JW375 |
| Primer030 | GCAGCCAGGAGCTGATCT | |
| Primer031 | ACGAAATCGTGGCGGTGC | Amplification of upstream homologous arm of *pbp1B* for JW378 |
| Primer032 | GCTACCTCCTGGGGCGGGCGCATACCACATGGATCCTTATTTGCCGACTAC | Amplification downstream homologous arm of *aph* for JW378 |
| Primer033 | CGCCCGCCCCAGGAGGTAGCGTGAGCAAAGGCGAAGAAG | Amplification of *mng* for JW378 |
| Primer034 | GCCCGGTGCCGGTGCCGGGCTACCACCGCCACCTTTATACAGTTCATCCATGCCC | |
| Primer035 | GCCCGGCACCGGCACCGGGCGGTGGCGGTAGCGGCCCGGCATTGATTTTC | Amplification of *pbp1B*$^{25-425}$ for JW378 |
| Primer036 | TCGAGCCGCCTTGCACCT | |
| Primer037 | AGCCCGGCACCGGCACCGGGCGGTGGCGGTAGCGAAATCGGTACTGGCTTTCC | Amplification of *halo* for JW337 |
| Primer038 | ATAAGGATCCTCAACCGGAAATCTCCAGAGTAG | |
| Primer039 | GATTTCCGGTTGAGGATCCTTATTTGCCGACTA | Amplification of *aph* for JW337 |
| Primer040 | TCACTCACAGGAGGACCCCATATGAGTAAAGGAGAAGAACTTTTCA | Amplification of *gfpmut2* for pJW18 |
| Primer041 | TGCAGGTCGAATCGGATCCTATTATTTGTATAGTTCATCCATGCC | |
| Primer042 | TAGGATCCGATTCGACCTGCAGGCAT | Amplification of pRADK backbone for pJW18 |

| | | |
|---|---|---|
| Primer043 | ATGGGGTCCTCCTGTGAGTGAGATGG | |
| Primer044 | AGAATTCGGCTCCACGCACCATCTCTAACGAC | Amplification of $P_{1261}$ for pJW31 |
| Primer045 | CTTTACTCATGGAAGTTACCCTAAGTCCCGTC | |
| Primer046 | GGTAACTTCCATGAGTAAAGGAGAAGAACTTTTC | Amplification of pJW18 backbone for pJW31 |
| Primer047 | ATGGTGCGTGGAGCCGAATTCTGCAGATGACGCGT | |
| Primer048 | GCGGGCCAATTGGAAGCACGTATTGTCGCCCT | Amplification of $P_{groes}$ for pJW32 |
| Primer049 | TTTTTCTCCATTGGGGTCCTCCTGTGAGTGAG | |
| Primer050 | GAGGACCCCAATGGAGAAAAAATCACTGGATATACCACC | Amplification of pJW31 backbone for pJW32 |
| Primer051 | ACGTGCTTCCAATTGGCCCGCCGCCTGTTTC | |
| Primer052 | GGCCTTTTGCTAAGCTTCTGCAGACGCGTCATCT | Amplification of $P_{1261}$-*gfpmut2* from pJW32 for pJW33 |
| Primer053 | CCCCGGATTGCCACCGGGGGGATTTTTATGCCTCTAGCACGCGTACCAT | |
| Primer054 | CCCCCGGTGGCAATCCGGGGGGTTTTTTCGCTTCTCTATTTCAATATCTAAA | Amplification of pJW32 backbone gene for pJW33 |
| Primer055 | CTGCAGAAGCTTAGCAAAAGGCCAGCAAAAGG | |
| Primer056 | TCACTCACAGGAGGACCCCAATGAGCCATATTCAACGGGAAAC | Amplification of *aph* for pJWK |
| Primer057 | GAGTAAACTTGGTCTGACAGCTAGAAAAACTCATCGAGCATC | |
| Primer058 | CTGTCAGACCAAGTTTACTC | Amplification of pJW33 backbone for pJWK |
| Primer059 | TGGGGTCCTCCTGTGAGT | |
| Primer060 | GGGACTTAGGGTAACTTCCTATGGTGAGCAAGGGCGAGGA | Amplification of *mch* for pJWK1 |
| Primer061 | TGGGATCCCCCGGGCTGCAGTTACTTGTACAGCTCGTCCATGC | |
| Primer062 | CTGCAGCCCGGGGGATC | Amplification of pJWK backbone for pJWK1 |
| Primer063 | GGAAGTTACCCTAAGTCCCG | |
| Primer064 | GTCATCTGCAGAATTCGGCTGCTAAAATTCCTGAAAAATTTTGC | Amplification of $P_{spac1}$ for pJWK64 |

| | | |
|---|---|---|
| Primer065 | ATCTAGTATTTCTCCTCTTTAAGCTTAATTGTGAGCGGCT | |
| Primer066 | TTAAAGAGGAGAAATACTAGATGGTGAGCAAGGGCGAG | Amplification of pJWK1 backbone for pJWK64 |
| Primer067 | AGCCGAATTCTGCAGATGAC | |
| Primer068 | GAATTCGGCTTCCCTATCAGTGATAGAGATTG | Amplification of $P_{tetdr}$ for pJWK67 |
| Primer069 | GCGGTACCTTTCTCCTCTTTAATGAATTCGGTCAGTGCGT | |
| Primer070 | AAAGAGGAGAAAGGTACCGCATGGTGAGCAAGGGCGAG | Amplification of *mch* for pJWK67 |
| Primer071 | TTTGCCCGCTAGATTACTTGTACAGCTCGTCCA | |
| Primer072 | CAAGTAATCTAGCGGGCAAAACCCAGGAGGAAAGACTCATGTCTAGATTAGATAAAAGTAAAG | Amplification of *TetR tn10* for pJWK67 |
| Primer073 | GCTGCAGTTATTAAGACCCACTTTCACATTTAAG | |
| Primer074 | TGGGTCTTAATAACTGCAGCCCGGGGG | Amplification of pJWK1 backbone for pJWK67 |
| Primer075 | CTGATAGGGAAGCCGAATTCTGCAGATGAC | |
| Primer076 | CGGGACTTAGGGTAACTTCCTATGCAAGCAGCCAGAATTCGCG | Amplification of *ftsZ* for pJH120 |
| Primer077 | GCTACCGCCACCGCCCGG | |
| Primer078 | CACCGGGCGGTGGCGGTAGCGTGAGCAAAGGCGAAGAAGAT | Amplification of *mng* for pJH120 |
| Primer079 | CCCCCGGGCTGCAGTTATTATTTATACAGTTCATCCATGCCCA | |
| Primer080 | TAATAACTGCAGCCCGGGGGATC | Amplification of pJWK backbone for pJH120 |
| Primer081 | AGGAAGTTACCCTAAGTCCC | |
| Primer082 | GAGGAGAAAGGTACCGCATGCAAGCAGCCAGAATTCGC | Amplification of *ftsZ-linker0-mng* for pJH150 |
| Primer083 | TGGGTTTTGCCCGCTAGATTATTTATACAGTTCATCCATGCCCA | |
| Primer084 | TAATCTAGCGGGCAAAACCCAG | Amplification of pJWK67 backbone for pJH150 |
| Primer085 | CATGCGGTACCTTTCTCCTCTTT | |
| Primer086 | GAGGAGAAAGGTACCGCATGGTGAGCAAAGGCGAAGAAGA | Amplification of *mng* for pJH158 |

| | | |
|---|---|---|
| Primer087 | CACCGGGCGGTGGCGGTAGCATGGAACTGAAGATTCGGAACC | Amplification of *pbp2* for pJH158 |
| Primer088 | CTGGGTTTTGCCCGCTAGATTACTTCGCCGGTTCGCTG | |
| Primer089 | GCGCACGCCGGCGAAGTCGAGGTTGATCATCC | Amplification of $\mathit{ftsZ}^{1\text{-}621}$ for making pJH180 containing $\mathrm{FtsZ}^{\mathrm{D205G}}$ |
| Primer090 | CTTCGCCGGCGTGCGCAACCTGCTCGCCAACTC | Amplification of $\mathit{ftsZ}^{606\text{-}1113}$ for making pJH180 containing $\mathrm{FtsZ}^{\mathrm{D205G}}$ |
| Primer091 | CATTGTGATTGCTCCTTGAAAAGTTACTTCGCCGGTTCGCTG | Amplification of *pbp3* for pJH195 |
| Primer092 | CTTTTCAAGGAGCAATCACAATGCAAGCAGCCAGAATTCGC | Amplification of *ftsZ* for pJH195 |
| Primer093 | CACCGGGCGGTGGCGGTAGCGTGAGCAAGGGCGAGGA | Amplification of *mch* for pJH195 |
| Primer094 | CCTGGGTTTTGCCCGCTAGATTACTTGTACAGCTCGTCCA | |
| Primer095 | AAAGAGGAGAAAGGTACCGCATGTGGTATGCGCCCGC | Amplification of *mng* for pJWK81 |
| Primer096 | CCTGGGTTTTGCCCGCTAGATCAGGGCTGCGCGGGT | Amplification of *pbp1B* for pJWK81 |
| Primer097 | TGATCTAGCGGGCAAAACCCAG | Amplification of pJWK67 backbone for pJWK81 |
| Primer098 | AGCGCGAACCGCTCTTGTACAGCTCGTCCATGC | Amplification of *mch* for pJWK7 |
| Primer099 | GTACAAGAGCGGTTCGCGCTCGTCCCAGG | Amplification of *pbp1A* for pJWK7 |
| Primer100 | TGGGATCCCCCGGGCTGCAGTTATGGCCTCGGCGGCA | |
| Primer101 | GATGGCGCGAGCCTGAC | Validation of the *mng* insertion |
| Primer102 | CATCCATGCCCATCACATCG | |
| Primer103 | AGTTCATGCGCTTCAAGGTG | Validation of the *mch* insertion |
| Primer104 | GATGGTGTAGTCCTCGTTGT | |
| Primer105 | GAGCGCATGCACTACGTC | Validation of the *halo* insertion |
| Primer106 | CAGCAGATTCAGACCCGG | |
| Primer107 | GATAACAATTAAGCTTTTCCCGGGAGGAGGTACCTTATGGTGTCAAAGGGTGAAGA | Amplification of *mng* for pJWK88 |
| Primer108 | CTCCGCCAGAACCGCCTCCACCCTTGTATAACTCATCCATGCCC | |

| | | |
|---|---|---|
| Primer109 | GTGGAGGCGGTTCTGGCGGAGGTGGCTCTACGGAAAAC AAAGGATCTTCTCAG | Amplification of *sapbp2* for pJWK88 |
| Primer110 | CCTAGTTTTTAGTTGAATATACCTGTTAATC | |
| Primer111 | AACAGGTATATTCAACTAAAAAACTAGGAGGAAATTTA AATGTTAGAATTTGAACAAGG | Amplification of $saftsZ^{1-165}$ for pJWK88 |
| Primer112 | AGAGCCACCTCCGCCAGAACCGCCTCCACCAGCTTTA | |
| Primer113 | TCTAAAGCTGGTGGAGGCGGTTCTGGCGGAGGTGGCTC TGCTGAAATCGGTACG | Amplification of *halo* for pJWK88 |
| Primer114 | AGATTCAGATCCGCCGCCACCAGACCCACCACCGCCTC CAGATATTTCTAACGTAC | |
| Primer115 | GGCGGTGGTGGGTCTGGTGGCGGCGGATCTGAATCT | Amplification of $saftsZ^{166-1173}$ for pJWK88 |
| Primer116 | CGCCTCGGAGCGACTCGAGTTAACGTCTTGTTCTTCTTG AA | |
| Primer117 | CTCGAGTCGCTCCGAGGC | Amplification of pCN5 backbone for pJWK88 |
| Primer118 | CCCGGGAAAAGCTTAATTGTTATC | |
| Primer119 | ATTAAGCTTTTCCCGGGAGGAGGTACCTTATGAAGAAT TTTAGAAGTATT | Amplification of *saftsw* for pJWK89 |
| Primer120 | GACCAGATCCTGGTCCTTGTCCTGATCCTTCCAAATTAA ATTGTCTTCTTATAT | |
| Primer121 | TCAGGACAAGGACCAGGATCTGGTCAAGGTTCTGGTGC TGAAATCGGTACGGGATT | Amplification of *halo* for pJWK89 |
| Primer122 | CATTTAAATTTCCTCCTAGTTTTTTATCCAGATATTTCTA ACGTACTT | |
| Primer123 | GATAAAAAACTAGGAGGAAATTTAAATGTTAGAATTTG AACAAGGAT | Amplification of $saftsZ^{1-165}$ for pJWK89 |
| Primer124 | TCTAAAGCTGGTGGAGGCGGTTCTGGCGGAGGTGGCTC TGTGTCAAAGGGTGAAGAAG | Amplification of *mng* for pJWK89 |
| Primer125 | AGATTCAGATCCGCCGCCACCAGACCCACCACCGCCCT TGTATAACTCATCCATGC | |
| Primer126 | ATATTCAACTAAAAAACTAGGAGGAAATTTAAATGAAG AATTTTAGAAGTATTTTAC | Amplification of *saftsw* for pJWK90 |
| Primer127 | GCGCCTCGGAGCGACTCGAGTTATCCAGATATTTCTAA CGTA | Amplification of *halo* for pJWK90 |
| Primer128 | TAAGAAGGAGATATACCATGCGCGACCTGCCCGAAGTC | Amplification of $pbp1A^{\Delta 1-73}$-$His_{10}$ for pJH130 |
| Primer129 | GAACCGTGGTGGTGATGGTGATGATGGTGGTGATGCTC GAGTGGCCTCGGCGGCAGCG | |
| Primer130 | ATCACCACCACGGTTCTGATGAGGTTGATGCTGGATCC AGCGCCTGGTCCCACCCTCA | Amplification of *Twin-Strep-tag* for pJH130 |

| | | |
|---|---|---|
| Primer131 | GTTAGCAGCCGGATCTCACTTTTCGAACTGGGGGTGGCTC | |
| Primer132 | TGAGATCCGGCTGCTAACAAAG | Amplification of pET-28a(+) backbone for pJH130, pJH131 and pJH132 |
| Primer133 | CATGGTATATCTCCTTCTTAAAGTTAA | |
| Primer134 | AAGAAGGAGATATACCATGCCCGATTACCGCGAGCTCG | Amplification of *pbp1B*$^{\Delta 1\text{-}48}$-*His*$_{10}$ for pJH131 |
| Primer135 | GAACCGTGGTGGTGATGGTGATGATGGTGGTGATGCTCGAGGGGCTGCGCGGGTTTCTG | |
| Primer136 | CATCACCACCACGGTTCTGATGAGGTTGATGCTGGATCCAGCGCCTGGTCCCACCCTCA | Amplification of *Twin-Strep-tag* for pJH131 |
| Primer137 | CTTTAAGAAGGAGATATACCATGAACTCGATTACCACCAAGC | Amplification of *pbp3*$^{\Delta 1\text{-}35}$-*His*$_{10}$ for pJH132 |
| Primer138 | GAACCGTGGTGGTGATGGTGATGATGGTGGTGATGCTCGAGCTTCGCCGGTTCGCTGAC | |
| Primer139 | CATCACCACCACGGTTCTGATGAGGTTGATGCTGGATCCAGCGCCTGGTCCCACCCTCA | Amplification of *Twin-Strep-tag* for pJH132 |

Note:
a. Underlined sequences indicate regions complementary to the target gene or DNA fragments, used for hybridization.

**Supplementary Table 3. Cell culture, imaging conditions, and statistics of each figure.**

| Figure | Strain | Culture and labeling | Imaging | Statistics |
|---|---|---|---|---|
| 1F; 1H | DRWT | 2×TGY, 30 °C;<br>50 µM TMRDA, 2 h[a]. | Excitation: 549 nm;<br>Intensity: 1.00%;<br>Emission: 562-629 nm;<br>Frame Average 4;<br>Line Accumulation 3.<br>**(Confocal)** | Three replicates;<br>48 cells |
| 1I | | 2×TGY, 30 °C;<br>25 µM TMRDA, 45 min[a]. | | Three replicates;<br>Demography:<br>2000 cells. |
| S1B, column1 | DRWT | 2×TGY, 30 °C;<br>1 µM TMRDA, 2 h[a]. | Excitation: 549 nm;<br>Intensity: 1.00%;<br>Emission: 562-629 nm;<br>Frame Average 4;<br>Line Accumulation 3.<br>**(Confocal)** | Three replicates;<br>> 200 cells |
| S1B, column2 | | 2×TGY, 30 °C;<br>5 µM TMRDA, 2 h[a]. | | Three replicates;<br>> 200 cells |
| S1B, column3 | | 2×TGY, 30 °C;<br>10 µM TMRDA, 2 h[a]. | | Three replicates;<br>> 200 cells |
| S1B, column4 | | 2×TGY, 30 °C;<br>25 µM TMRDA, 2 h[a]. | | Three replicates;<br>> 200 cells |
| S1B, column5 | | 2×TGY, 30 °C;<br>50 µM TMRDA, 2 h[a]. | | Three replicates;<br>> 100 cells |
| S1B, column6 | | 2×TGY, 30 °C;<br>100 µM TMRDA, 2 h[a]. | | Three replicates;<br>> 200 cells |
| S1G, row1 | DRWT | 2×TGY, 30 °C;<br>5 µM RhoDA;<br>0 µM TMRDA, 2 h[a]. | Excitation 488 nm;<br>Intensity: 4.00%;<br>Emission range: 502-537 nm;<br>Frame Average 4;<br>Line Accumulation 3.<br>**(Confocal)** | Three replicates;<br>> 100 cells |
| S1G, row2 | | 2×TGY, 30 °C;<br>5 µM RhoDA;<br>1 µM TMRDA, 2 h[a]. | | Three replicates;<br>> 100 cells |
| S1G, row3 | | 2×TGY, 30 °C;<br>5 µM RhoDA;<br>5 µM TMRDA, 2 h[a]. | | Three replicates;<br>> 100 cells |
| S1G, row4 | | 2×TGY, 30 °C;<br>5 µM RhoDA;<br>10 µM TMRDA, 2 h[a]. | | Three replicates;<br>> 100 cells |
| S1G, row5 | | 2×TGY, 30 °C;<br>5 µM RhoDA;<br>25 µM TMRDA, 2 h[a]. | | Three replicates;<br>> 100 cells |
| 2A, column1;<br>2C, column1 and 6 | JH304 | 2×TGY, 30 °C;<br>2 µM Potomac Red[b]. | 488 nm, 55 $W/cm^2$, 50 ms;<br>647 nm, 10 $W/cm^2$, 50 ms.<br>**(HiLO, Excitation, Power density, exposure time)** | Three replicates;<br>Demography:<br>1207 cells. |
| 2A, column2;<br>2C, column2 | JW335 | 2×TGY, 30 °C;<br>2 µM Potomac Red[b]. | 488 nm, 55 $W/cm^2$, 50 ms;<br>647 nm, 10 $W/cm^2$, 50 ms.<br>**(HiLO, Excitation, Power density, exposure time)** | Three replicates;<br>Demography:<br>1197 cells. |
| 2A, column3;<br>2C, column3 | JH312 | 2×TGY, 30 °C;<br>2 µM Potomac Red[b]. | 488 nm, 55 $W/cm^2$, 50 ms;<br>647 nm, 10 $W/cm^2$, 50 ms.<br>**(HiLO, Excitation, Power density, exposure time)** | Three replicates;<br>Demography:<br>1070 cells. |
| 2A, column4;<br>2C, column4 | JH378 | 2×TGY, 30 °C;<br>2 µM Potomac Red[b]. | 488 nm, 55 $W/cm^2$, 50 ms;<br>647 nm, 10 $W/cm^2$, 50 ms.<br>**(Excitation, Power density, exposure time)** | Three replicates;<br>Demography:<br>1233 cells. |
| 2A, column5;<br>2C, column5 | JH375 | 2×TGY, 30 °C;<br>2 µM Potomac Red[b];<br>10 µM JF503-Halo, 1 h. | 488 nm, 55 $W/cm^2$, 50 ms;<br>647 nm, 10 $W/cm^2$, 50 ms.<br>**(HiLO, Excitation, Power density, exposure time)** | Three replicates;<br>Demography:<br>1374 cells. |
| 2F; 2G; S5C, column2 | JW441 | 2×TGY, 30 °C;<br>2 µM Potomac Red[b];<br>2 µM JF552-Halo, 1 h. | 488 nm, 55 $W/cm^2$, 50 ms;<br>561 nm, 56 $W/cm^2$, 50 ms;<br>647 nm, 8 $W/cm^2$, 50 ms.<br>**(HiLO, Excitation, Power density, exposure time)** | Three replicates;<br>PCC: 609 cells.<br>Demography:<br>560 cells. |

| | | | | |
|---|---|---|---|---|
| S4B, row1 | JH213 | 2×TGY, 30 °C. | 488 nm, 55 $W/cm^2$, 50 ms. **(HiLO, Excitation, Power density, exposure time)** | One replicates. |
| S4B, row2 | JW330 | 2×TGY, 30 °C. | 488 nm, 55 $W/cm^2$, 50 ms. **(HiLO, Excitation, Power density, exposure time)** | One replicates. |
| S5A, row1 | JW425 | 2×TGY, 30 °C;<br>2 μM Potomac Red[b];<br>2 μM JF552-Halo, 1 h. | 488 nm, 55 $W/cm^2$, 50 ms;<br>561 nm, 56 $W/cm^2$, 50 ms;<br>647 nm, 10 $W/cm^2$, 50 ms.<br>**(HiLO, Excitation, Power density, exposure time)** | Three replicates;<br>PCC: 836 cells (with membrane);<br>PCC: 406 cells (with FtsZ). |
| S5A, row2 | JH337 | 2×TGY, 30 °C;<br>2 μM Potomac Red[b]. | 488 nm, 55 $W/cm^2$, 50 ms;<br>561 nm, 56 $W/cm^2$, 50 ms;<br>647 nm, 16 $W/cm^2$, 50 ms.<br>**(HiLO, Excitation, Power density, exposure time)** | Three replicates;<br>PCC: 949 cells (with membrane);<br>PCC: 955 cells (with FtsZ). |
| S5A, row3 | JW445 | 2×TGY, 30 °C;<br>2 μM Potomac Red[b]. | 488 nm, 55 $W/cm^2$, 50 ms;<br>561 nm, 56 $W/cm^2$, 50 ms;<br>647 nm, 33 $W/cm^2$, 50 ms.<br>**(HiLO, Excitation, Power density, exposure time)** | Three replicates;<br>PCC: 858 cells (with membrane);<br>PCC: 868 cells (with FtsZ). |
| S5A, row4 | JW423 | 2×TGY, 30 °C;<br>2 μM Potomac Red[b];<br>2 μM JF552-Halo, 1 h. | 488 nm, 55 $W/cm^2$, 50 ms;<br>561 nm, 56 $W/cm^2$, 50 ms;<br>647 nm, 16 $W/cm^2$, 50 ms.<br>**(HiLO, Excitation, Power density, exposure time)** | Three replicates;<br>PCC: 621 cells (with membrane)<br>PCC: 380 cells (with FtsZ). |
| S5C, column1; | JW440 | 2×TGY, 30 °C;<br>2 μM Potomac Red[b];<br>2 μM JF552-Halo, 1 h. | 488 nm, 55 $W/cm^2$, 50 ms;<br>561 nm, 56 $W/cm^2$, 50 ms;<br>647 nm, 8 $W/cm^2$, 50 ms.<br>**(HiLO, Excitation, Power density, exposure time)** | Three replicates;<br>PCC: 859 cells. |
| S5C, column3; | JH384 | 2×TGY, 30 °C;<br>2 μM Potomac Red[b]; | 488 nm, 55 $W/cm^2$, 50 ms;<br>561 nm, 56 $W/cm^2$, 50 ms;<br>647 nm, 16 $W/cm^2$, 50 ms.<br>**(HiLO, Excitation, Power density, exposure time)** | Three replicates;<br>PCC: 631 cells. |
| 3A, column1; S7A | DRWT | 2×TGY, 30 °C;<br>2 μM Cy5DA , 1 h[a]. | Excitation 642 nm<br>Intensity: 1.0 $kW/cm^2$<br>Exposure Time: 25ms<br>Emission filter: 667/30 and 690/50<br>(see 3D-SMLM imaging part) | 79 septa |
| 3A, column2; S7B | | 2×TGY, 30 °C;<br>16 μg/mL aztreonam, 1 h;<br>2 μM Cy5DA, 1 h[a]. | | 46 septa |
| 3A, column3; S7C | | 2×TGY, 30 °C;<br>62.5 μg/mL cefsulodin, 1 h;<br>10 μM Cy5DA, 1 h[a]. | | 91 septa |
| 3A, column4; S7D | | 2×TGY, 30 °C;<br>2 μg/mL moenomycin, 1 h;<br>6 μM Cy5DA, 1 h[a]. | | 13 cells |
| 3B, WT; S8A; S8E. | DRWT | 2×TGY, 30 °C. | Tomography<br>Angular range: −60° to +60°;<br>Intervals: 3° ;<br>1-second exposure time. | 7 cells |
| 3B, Aztreonam; S8B. | | 2×TGY, 30 °C;<br>16 μg/mL aztreonam, 1 h. | | 8 cells |
| 3B, Cefsulodin; S8C; S8F. | | 2×TGY, 30°C;<br>62.5 μg/mL cefsulodin, 1 h. | | 11 cells |

| | | | | |
|---|---|---|---|---|
| 3B, Moenomycin; S8D; S8G. | | 2×TGY, 30 °C; 2 µg/mL moenomycin, 1 h. | | 13 cells |
| 3D, row1 | DRWT | 2×TGY, 30 °C; 50 µM TMRDA, 1 h[a]. | Excitation: 549 nm; Intensity: 2.00%; Emission: 562-629 nm; Frame Average 4; Line Accumulation 3. **(Confocal)** | Three replicates; > 500 cells |
| 3D, row2 | | 2×TGY, 30 °C; 32 µg/mL aztreonam, 1 h; 50 µM TMRDA, 1 h[a]. | | Three replicates; > 200 cells |
| 3D, row3 | | 2×TGY, 30 °C; 62.5 µg/mL cefsulodin, 1 h; 50 µM TMRDA, 1 h[a]. | | Three replicates; > 200 cells |
| 3D, row4 | | 2×TGY, 30 °C; 4 µg/mL moenomycin, 1 h; 50 µM TMRDA, 1 h[a]. | | Three replicates; > 900 cells |
| S9D, row1 | JW117 | 2×TGY, 30 °C; 50 µM TMRDA, 2 h[a]. | Excitation 549 nm; Intensity: 1.90%; Emission range: 562-629 nm; Frame Average: 4; Line Accumulation: 3. **(Confocal)** | Three replicates; > 200 cells |
| S9D, row2 | JW116 | | | Three replicates; > 400 cells |
| 3F, row1 | DRWT | 2×TGY, 30 °C; 100 nM Potomac Gold[b]. | Time-lapse 561 nm, 11 W/cm$^2$, 50ms, 5min. **(HiLO, Excitation, Power density, exposure time, Intervals)** | 33 cells |
| 3F, row2 | | 2×TGY, 30 °C; 100 nM Potomac Gold[b]; 1.6 µg/mL aztreonam. | | 47 cells |
| 3F, row3 | | 2×TGY, 30 °C; 100 nM Potomac Gold[b]; 6.25 µg/mL cefsulodin. | | 114 cells |
| 3F, row4 | | 2×TGY, 30 °C; 100 nM Potomac Gold[b]; 0.2 µg/mL moenomycin. | | 108 cells |
| 4B; S10A; S10I; S10J | JW337 | 2×TGY, 30 °C; 100 nM Potomac Gold[b]; 0.02 nM JFX650-Halo, 30 min. | Splitter 561 nm, 90 W/cm$^2$, 50 ms, 60 frames; 647 nm, 5 W/cm$^2$, 1000 ms, 200 frames. **(HiLO, Excitation, Power density, exposure time)** | Three replicates. |
| 4C; S10B | JW449 | | | |
| S10C | JW337 | 2×TGY, 30 °C; 0.02 nM JFX650-Halo, 30 min[c]. | 647 nm, 5 W/cm$^2$, 1000 ms, 200 frames. **(HiLO, Excitation, Power density, exposure time)** | Three replicates. |
| S10G, row1 | JH213 | 2×TGY, 30 °C. | 488 nm, 21 W/cm$^2$, 50 ms, 1-second intervals, 100 frames. **(HiLO)** | Three replicates. 111 cells |
| S10G, row2; S6H | JH317 | | | Three replicates. 76 cells |
| 5B | JW462 | TSB, 37°C; 2 µM JFX650-Halo, 30 min. | 488 nm, 21 W/cm$^2$, 1000 ms; 647 nm, 3 W/cm$^2$, 50 ms. **(TIRF, Excitation, Power density, exposure time)** | Three replicates |
| S11AB | JW461 | TSB, 37 °C; 2 µM JFX650-Halo, 30 min. | 488 nm, 55 W/cm$^2$, 50 ms; 647 nm, 3 W/cm$^2$, 50 ms. **(TIRF, Excitation, Power density, exposure time)** | Three replicates; Demography: 466 cells |

| | | | | |
|---|---|---|---|---|
| S11CD | JW460 | TSB, 37 ℃;<br>2 μM JFX650-Halo, 30 min. | 488 nm, 55 W/cm$^2$, 50 ms;<br>647 nm, 3 W/cm$^2$, 50 ms.<br>**(TIRF, Excitation, Power density, exposure time)** | Three replicates;<br>Demography:<br>218 cells |
| 5E, row1 | *S. aureus* RN4220 | TSB, 37 ℃;<br>FADA$_{560}$[17], 1 μM[b]. | Time-lapse<br>488 nm, 5 W/cm$^2$,<br>50 ms, 5 min.<br>**(TIRF, Excitation, Power density, exposure time, Intervals)** | Three replicates |
| 5E, row2 | | TSB, 37 ℃;<br>FADA$_{560}$, 1 μM ;<br>4 μg/mL meropenem[b]. | | Three replicates |
| 5E, row3 | | TSB, 37 ℃;<br>FADA$_{560}$, 1 μM ;<br>2 μg/mL cefotaxime[b]. | | Three replicates |
| S11A, row1 | *S. aureus* RN4220 | TSB, 37 ℃;<br>10 μM RhoDA, 10 min[a]. | Excitation 488 nm;<br>Intensity: 4.00%;<br>Emission range: 502-537 nm;<br>Frame Average 4;<br>Line Accumulation 3.<br>**(Confocal)** | Three replicates;<br>> 700 cells |
| S11A, row2 | | TSB, 37 ℃;<br>4 μg/mL meropenem, 30 min;<br>10 μM RhoDA, 10 min[a]. | | Three replicates;<br>> 800 cells |
| S11A, row3 | | TSB, 37 ℃;<br>2 μg/mL cefotaxime, 30 min;<br>10 μM RhoDA, 10 min[a]. | | Three replicates;<br>> 600 cells |

Note:
a. Samples were labeled with shaking at 30 °C, followed by fixation with 75% ethanol.
b. Included in the agarose gel pad.
c. Following labeling, the cells were washed with MM medium and subsequently fixed with 2.6% paraformaldehyde and 0.05% glutaraldehyde.

**Supplementary Table 4. Septal thickness measured by 3D-SMLM and electron-TOMO.**

| Treatment | Septum | $W_{SMLM}$ (μ±s.d. in nm), n | $W_{TOMO}$ (μ±s.d. in nm), n |
|---|---|---|---|
| Untreated | S0 | 109.7±20.6, 51 | 29.0±4.5, 9 |
| | S1 leading[a] | NA[b] | 20.1±3.0, 14 |
| | S1 | 106.2±30.9, 65 | 24.7±2.9, 14 |
| Aztreonam | S1 | 146.4±25.7, 80 | 47.5±15.3, 30 |
| Cefsulodin | S0 | 85.9±17.8, 118 | 19.0±2.4, 9 |
| | S1 | 85.9±21.1, 42 | 19.8±2.6, 22 |
| Moenomycin | S0 | 88.3±20.4, 89 | 19.3±1.8, 13 |
| | S1 | NA[c] | 19.3±1.8, 13 |

Note:
a. The thickness of S1 without the wedge (>50 nm distance from the peripheral PG).
b. Note measured
c. Few moenomycin treated cells in the SMLM experiment showed unclosed septa (S1).

**Supplementary Table 5. Single-molecule tracking results and FtsZ treadmilling speed distributions.**

| Single-molecule tracking results | | | | | | | |
|---|---|---|---|---|---|---|---|
| **POI** | **Genotype** | **Culture** | **Position** | **$P_{moving}$% (s.e.m)** | **$P_{V1}$% (s.e.m)** | **$V_1$ (nm/s) (s.e.m)** | **$V_2$ (nm/s) (s.e.m)** |
| FtsW | WT | 2×TGY | leading edge of S1 | 17.8 (1.7) | 18.1 (21.6) | 2.7 (3.6) | 15.5 (1.4) |
| | WT | +Aztreonam | | 7.3 (1.2) | 100 (NA) | 14.9 (1.3) | NA[a] (NA) |
| | WT | +Cefsulodin | | 20.7 (3.7) | 13.4 (28.6) | 2.1 (4.2) | 13.2 (2.1) |
| | $Z^{D205G}$ | 2×TGY | | 13.5 (3.2) | 75.4 (22.5) | 4.0 (0.9) | 12.7 (2.5) |
| PBP1B | WT | 2×TGY | S0 | 9.5 (1.4) | 100 (NA) | 12.3 (0.9) | NA[a] (NA) |
| | WT | +Aztreonam | | 18.8 (0.7) | 100 (NA) | 12.6 (0.7) | NA[a] (NA) |
| | WT | +Cefsulodin | | 1.0 (0.6) | NA[b] (NA) | NA (NA) | NA (NA) |
| | WT | +Moenomycin | | 1.0 (0.2) | NA[b] (NA) | NA (NA) | NA (NA) |

| FtsZ treadmilling speed | | | | |
|---|---|---|---|---|
| **POI** | **Genotype** | **Culture** | **Treadmilling speed (nm/s) (s.e.m)** | ***Nz*** |
| FtsZ | WT | 2×TGY | 22.0 (0.4) | 273 |
| | D205G | 2×TGY | 6.5 (0.6) | 200 |

a. Single population of the velocity distribution.
b. Few moving molecules to construct the distribution.

**Supplementary Table 6. Directional velocity changes of FtsW movement trajectories after 2D-projection correction (unwrapping).**

| POI | Genotype | Culture | Position | Origin R | | 150% R | | 100% R | |
|---|---|---|---|---|---|---|---|---|---|
| | | | | *$V_1$* (nm/s) (s.e.m) | *$V_2$* (nm/s) (s.e.m) | *$V_1$* (nm/s) (s.e.m) | *$V_2$* (nm/s) (s.e.m) | *$V_1$* (nm/s) (s.e.m) | *$V_2$* (nm/s) (s.e.m) |
| FtsW | WT | 2×TGY | leading edge of S1 | 2.7 (5.2) | 15.5 (1.5) | 3.0 (6.3) | 17.2 (1.8) | 3.5 (7.3) | 21.0 (2.5) |
| | $Z^{D205G}$ | 2×TGY | | 4.0 (0.7) | 13.0 (3.1) | 4.5 (1.0) | 15.5 (5.3) | 5.5 (1.8) | 20.8 (5.7) |
| | WT | +Aztreonam | | 11.8 (4.6) | 14.1 (2.6) | 4.0 (7.9) | 15.6 (4.4) | 5.3 (13.4) | 20.7 (17.5) |
| | WT | +Cefsulodin | | 2.0 (4.0) | 13.0 (3.1) | 2.2 (3.7) | 14.5 (2.5) | 2.5 (5.0) | 17.9 (2.0) |

**Supplementary Table 7. Minimum inhibitory concentration (MIC).**

| Species | Antibiotics | MW(g/mol) | MIC(μg/mL) | MIC(μM) |
|---|---|---|---|---|
| *D. radiodurans* | Aztreonam | 435.43 | 3.2 | 7.3 |
| | Cefsulodin Sodium Salt Hydrate | 554.52 | 12.5 | 22.5 |
| | Moenomycin complex | 1583.58 | 0.4 | 0.25 |
| *S. aureus* | Meropenem | 383.46 | 2.0 | 5.2 |
| | Cefotaxime | 455.47 | 1.0 | 2.2 |

### General considerations of chemistry.

All reactions were carried out in oven-dried glassware under nitrogen atmosphere, unless stated otherwise. Commercial chemicals were purchased from Bidepharm, Macklin Inc. or Aladdin, and used without further purification.
Anhydrous solvents from Macklin were used without further treatment. Flash column chromatography was performed with silica gel (230-400 mesh). Reverse-phase semi-preparative high-pressure liquid chromatography was performed on Thermo Fisher ultimate 3000 system equipped with an LPG-3400SDN pump and an VWD-3100 RS detector for product visualization with a SUPELCO ascentis C18 HPLC column (5 μm, 25 cm × 21.2 mm) or a SUPELCO ascentis C18 HPLC column (5 μm, 25 cm × 10 mm). Nuclear magnetic resonance (NMR) spectra were recorded at room temperature on the AVANCE III 400 with chemical shifts (δ) reported in ppm relative to the solvent residual signals. $CDCl_3$: H 7.26 ppm, DMSO: H 2.5 ppm, $CD_3CN$: H 1.94 ppm. Coupling constants are reported in Hz. High resolution mass spectra (HRMS) were measured on a Orbitrap XL ETDTM HRMS system.

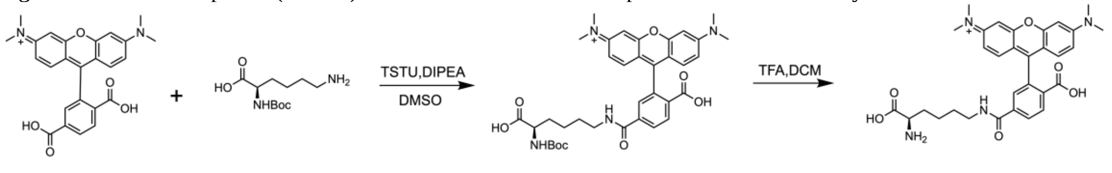


**Supplementary Scheme 1**. Synthetic route of TMRDA.

**Compound 1.** To a solution of 6-carboxytetramethylrhodamine (6-TMR, 0.035 mmol, 15.0 mg) in 200 μL dry DMSO, DIPEA (0.18 mmol, 31.3 μL) and a 0.032 mmol (9.6 mg) solution of TSTU in 200 μL dry DMSO were added. After stirring at room temperature for 10 min, the mixture was gradually added to a solution of Boc-D-Lys-OH (0.053 mmol, 13.1mg) in 400 μL dry DMSO. After stirring at room temperature for 40 min, the reaction was quenched by addition of 100 μL acetic acid. The product was purified by RP-HPLC (0.1% TFA in $H_2O$/acetonitrile 30 - 60%, 35 min) and lyophilized to give compound **1**. Retention time 16.8 min, yield 85%. **$^1$H NMR** (400 MHz, DMSO) 8.746 (t, J = 5.7 Hz, 1H), 8.282 (d, J = 8.2 Hz, 1H), 8.224 (dd, J = 8.2, 1.7 Hz, 1H), 7.851 (d, J = 1.7 Hz, 1H), 7.024 (m, 4H), 6.949 (s, 2H), 3.816 (dd, J = 9.7, 4.7 Hz, 1H), 3.251 (s, 14H), 1.909 – 1.443 (m, 4H), 1.337 (s, 11H). **HRMS** (ESI, pos. mode) m/z calc. for $C_{36}H_{43}N_4O_8^+$ 659.3075, found 659.3075 $[M]^+$.

**TMRDA** (**2**)**.** Compound **1** was dissolved in $CH_2Cl_2$/30% TFA. The resulting mixture was stirred at room temperature for 0.5 h. Then the solvent was removed by vacuum and the residue was further purified by RP-HPLC (0.1% TFA in $H_2O$/acetonitrile 15 - 40%, 36 min) and lyophilized to give **TMRDA** (**2**)**.** Retention time 19.2 min, yield 88%. **$^1$H NMR** (400 MHz, DMSO) δ 8.412 – 8.114 (m, 3H), 7.775 (s, 1H), 7.032 – 6.605 (m, 6H), 3.880 (t, $J$ = 6.2 Hz, 1H), 3.242 (m, 2H), 3.157 (s, 12H), 1.760 (m, 2H), 1.593 – 1.205 (m, 4H). **HRMS** (ESI, pos. mode) m/z calc. for $C_{31}H_{35}N_4O_6^+$ 559.2551, found 559.2559 $[M]^+$.

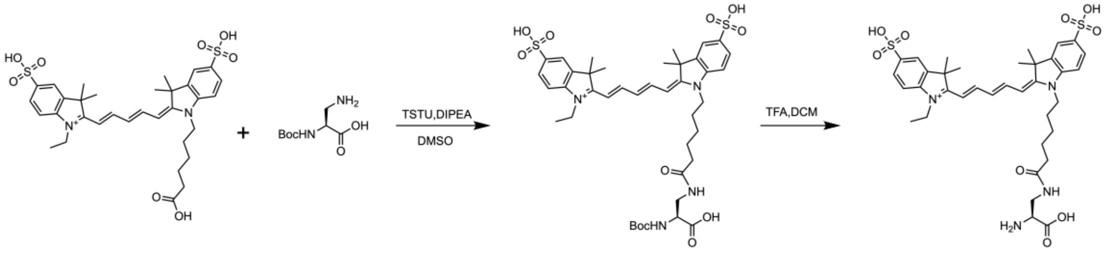


**Supplementary Scheme 2**. Synthetic route of Cy5DA.

**Compound 3.** To a solution of sulfo-cy5 carboxylic acid (0.023 mmol, 15.0 mg) in 200 μL dry DMSO, DIPEA (0.46 mmol, 8.0 μL) and a 0.021 mmol (6.3 mg) solution of TSTU in 200 μL dry DMSO were added. After stirring at room temperature for 10 min, the mixture was gradually added to a solution of Boc-D-Dap-OH (0.035 mmol, 7.1 mg) in 400 μL dry DMSO. After stirring at room temperature for 40 min, the reaction was quenched by addition of 100 μL acetic acid. The product was purified by RP-HPLC (0.1% TFA in $H_2O$/acetonitrile 15 - 45%, 35 min) and lyophilized to give compound **3**. Retention time 22.6 min, yield 86%. **$^1$H NMR** (400 MHz, DMSO) δ 8.353 (t, $J$ = 13.1 Hz, 2H), 7.820 – 7.800 (q, $J$ = 2.0 Hz, 2H), 7.664 – 7.624 (m, 2H), 7.344 – 7.298 (m, 2H), 6.619 – 6.557 (m, 1H), 6.325 – 6.271 (m, 2H), 4.135 – 4.007 (m, 5H), 3.314 – 3.193 (m, 2H), 2.072 – 2.042 (m, 2H), 1.683 (s, 14H), 1.580 – 1.461 (m, 2H), 1.350 (s, 11H), 1.259 (t, $J$ = 7.1 Hz, 3H). **HRMS** (ESI, pos. mode) m/z calc. for $C_{41}H_{55}N_4O_{11}S_2^+$ 843.3303, found 843.3325 $[M]^+$.

**Cy5DA** (**4**)**.** Compound **3** was dissolved in $CH_2Cl_2$/30% TFA. The resulting mixture was stirred at room temperature for 0.5 h. Then the solvent was removed by vacuum and the residue was further purified by RP-HPLC (0.1% TFA in $H_2O$/acetonitrile 10 - 45%, 36 min) and lyophilized to give compound **Cy5DA** (**4**)**.** Retention time 19.4 min, yield 82%. **$^1$H NMR** (400 MHz, DMSO) δ 8.540 – 8.267 (m, 2H), 7.808 (t, $J$ = 2.1 Hz, 2H), 7.650 (dt, $J$ = 8.2, 2.0 Hz, 2H), 7.320 (t, $J$ = 8.9 Hz, 2H), 6.578 (t, $J$ = 12.3 Hz, 1H), 6.298 (dd, $J$ = 13.8, 6.1 Hz, 2H), 4.113 (dt, $J$ = 16.1, 6.5 Hz, 4H), 3.921 (t, $J$ = 5.6 Hz, 1H), 3.430 – 3.323 (m, 2H), 2.231 – 2.006 (m, 2H), 1.681 (s, 14H), 1.518 (p, $J$ = 7.5 Hz, 2H), 1.319 – 1.240 (m, 5H). **HRMS** (ESI, pos. mode) m/z calc. for $C_{36}H_{47}N_4O_9S_2^+$ 743.2779, found 743.2792 $[M]^+$.

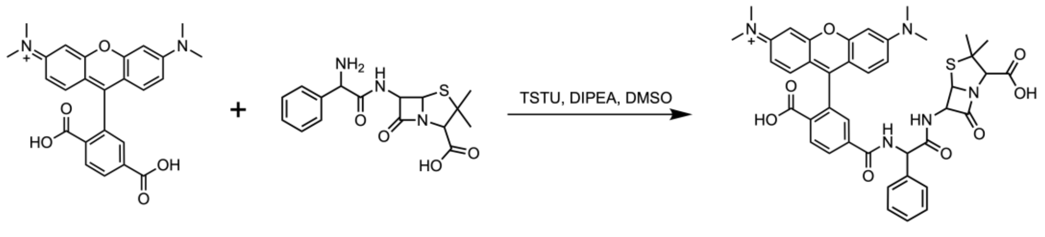


**Supplementary Scheme 3.** Synthesis of **ampicilin-TMR**.

**ampicilin-TMR (5).** The TSTU (0.055 mmol, 16.6 mg) was dissolved in DMSO and added dropwise to a solution of 6-TMR (0.046 mmol, 20.0 mg) and DIPEA (0.23 mmol, 40.1 μL) in DMSO. After stirring the mixture 15 minutes at room temperature, then the resulting solution was added dropwise to a solution of ampicillin sodium salt (0.092 mmol, 34.2 mg) and DIPEA (0.23 mmol, 40.1 μL) in DMSO was stirred at room temperature for 30 min. The reaction was quenched with acetic acid and was further purified by RP-HPLC (pH=6.0 100 mM ammonium acetate in $H_2O$/acetonitrile 20 - 65%, 30 min) and lyophilized to give **ampicilin-TMR (5)**. Retention time 15.5 min, yield 88%. **$^1$H NMR** (400 MHz, DMSO) δ 9.234 (d, J = 7.9 Hz, 1H), 9.019 (d, J = 7.6 Hz, 1H), 8.215 (dd, J = 8.0, 1.4 Hz, 1H), 8.057 (d, J = 8.0 Hz, 1H), 7.859 – 7.746 (m, 1H), 7.537 – 7.389 (m, 2H), 7.340 – 7.203 (m, 3H), 6.740 – 6.350 (m, 6H), 5.877 (d, J = 7.9 Hz, 1H), 5.478 (dd, J = 7.5, 4.0 Hz, 1H), 5.382 (d, J = 4.0 Hz, 1H), 4.152 (s, 1H), 2.939 (d, J = 2.6 Hz, 12H), 1.418 (s, 3H), 1.357 (s, 3H). **HRMS** (ESI, pos. mode) m/z calc. for $C_{41}H_{40}N_5O_8S^+$ 762.2592, found 762.2585 $[M]^+$.

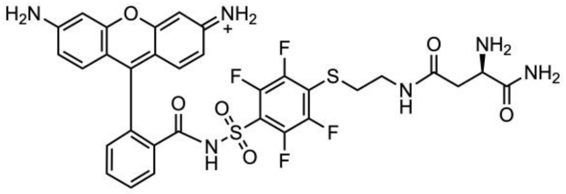


**Supplementary Scheme 4.** Chemical structure of **RhoDA**.

**RhoDA(6).** RhoDA was prepared according to the previously reported procedure[17]. RP-HPLC (0.1% TFA in $H_2O$/acetonitrile 20 - 95%, 35 min), retention time 17.3 min, yield 90%. **$^1$H NMR** (400 MHz, DMSO) δ 8.141 – 8.042 (m, 1H), 7.756 – 7.571 (m, 2H), 7.227 – 7.125 (m, 1H), 6.904 (d, J = 9.1 Hz, 2H), 6.723 (dd, J = 9.1, 2.1 Hz, 2H), 6.655 (d, J = 2.0 Hz, 2H), 3.964 (dd, J = 8.1, 4.7 Hz, 1H), 3.201 (h, J = 6.9 Hz, 2H), 3.009 (t, J = 7.0 Hz, 2H), 2.720 – 2.527 (m, 2H). **HRMS** (ESI, pos. mode) m/z calc. for $C_{32}H_{27}F_4N_6O_6S_2^+$ 731.1364, found 731.1362 $[M]^+$.

**Movie Captions**

***Supplementary Movie 1***

Time-lapse imaging (HiLO mode) of DRWT cells growing on 1.5% agarose gel pad with 2×TGY medium at 30 °C, corresponding to Figure 3F (row 1). The gel pad was supplemented with 200 nM Potomac Gold. The video was recorded every ~10 min. Scale bar, 1 μm.

***Supplementary Movie 2***

Time-lapse imaging (HiLO mode) of DRWT cells treated with 1.6 μg/mL aztreonam (0.5×MIC) growing on 1.5% agarose gel pad with 2×TGY medium at 30 °C, corresponding to Figure 3F (row 2). The gel pad was supplemented with 200 nM Potomac Gold and 1.6 μg/mL aztreonam. The video was recorded every ~10 min. Scale bar, 1 μm.

***Supplementary Movie 3***

Time-lapse imaging (HiLO mode) of DRWT cells treated with 6.25 μg/mL cefuslodin (0.5×MIC) growing on 1.5% agarose gel pad with 2×TGY medium at 30 °C, corresponding to 3F (row 3). The gel pad was supplemented with 200 nM Potomac Gold and 6.25 μg/mL cefsulodin. The video was recorded every ~10 min. Scale bar, 1 μm.

***Supplementary Movie 4***

Time-lapse imaging (HiLO mode) of DRWT cells treated with 0.2 μg/mL moenomycin (0.5×MIC) growing on 1.5% agarose gel pad with 2×TGY medium at 30 °C, corresponding to 3F (row 4). The gel pad was supplemented with 200 nM Potomac Gold and 0.2 μg/mL moenomycin. The video was recorded every ~10 min. Scale bar, 1 μm.

***Supplementary Movie 5***

Time-lapse imaging (HiLO mode) of fast single FtsW-Halotag molecules labeled with 0.02 nM JFX650 in MM medium, corresponding to Figure 4B. The gel pad was supplemented with 100 nM Potomac Gold. The arrow points to the start position of the molecule of interest. Scale bar, 1 μm.

***Supplementary Movie 6***

Time-lapse imaging (HiLO mode) of slow single FtsW-Halotag molecules labeled with 0.02 nM JFX650 in MM medium. The gel pad was supplemented with 100 nM Potomac Gold. The arrow points to the start position of the molecule of interest. Scale bar, 1 μm.

***Supplementary Movie 7***

Time-lapse imaging (HiLO mode) of single fixed FtsW-Halotag molecules prelabeled with 0.02nM JFX650 in MM medium, corresponding to Figure S10C. The arrow points to the start position of the molecule of interest. Scale bar, 1 μm.

***Supplementary Movie 8***

Time-lapse imaging (TIRF mode) of cells expressing mNG-FtsZ in 2×TGY medium, corresponding to Figure S10G (row 1). The arrow points to the Z-ring and indicates where the kymograph is extracted. Scale bar, 1 μm.

***Supplementary Movie 9***

Time-lapse imaging (TIRF mode) of cells over-expressing mNG-FtsZ$^{D205G}$ in 2×TGY medium, corresponding to Figure S10G (row 2). The arrow points to the Z-ring and indicates where the kymograph is extracted. Scale bar, 1 μm.

***Supplementary Movie 10***

Time-lapse imaging (HiLO mode) of single Halotag-PBP1B molecules labeled with 0.02 nM JFX650 in MM medium, corresponding to Figure S10B. The gel pad was supplemented with 100 nM Potomac Gold. The arrow points to the start position of the molecule of interest. Scale bar, 1 μm.

***Supplementary Movie 11***

Time-lapse imaging (HiLO mode) of single Halotag-PBP1B molecules labeled with 0.02 nM JFX650 in MM medium, corresponding to Figure 4C. The gel pad was supplemented with 100 nM Potomac Gold. The arrow points to the start position of the molecule of interest. Scale bar, 1 μm.

***Supplementary Movie 12***

Time-lapse imaging (TIRF mode) of *S. aureus* RN4220 cells growing on 1.5% agarose gel pad with TSB medium, corresponding to Figure 5E (row 1). The video was recorded every ~5 min. Scale bar, 500 nm.

***Supplementary Movie 13***

Time-lapse imaging (TIRF mode) of *S. aureus* RN4220 cells treated with 4 μg/mL meropenem (2×MIC) growing on 1.5% agarose gel pad with TSB medium, corresponding to Figure 5E (row 2). The video was recorded every ~5 min. Scale bar, 1000 nm.

***Supplementary Movie 14***

Time-lapse imaging (TIRF mode) of *S. aureus* RN4220 cells treated with 2 μg/mL cefotaxime (2×MIC) growing on 1.5% agarose gel pad with TSB medium, corresponding to Figure 5E (row 3). The video was recorded every ~5 min. Scale bar, 1000 nm.

**NMR spectra**

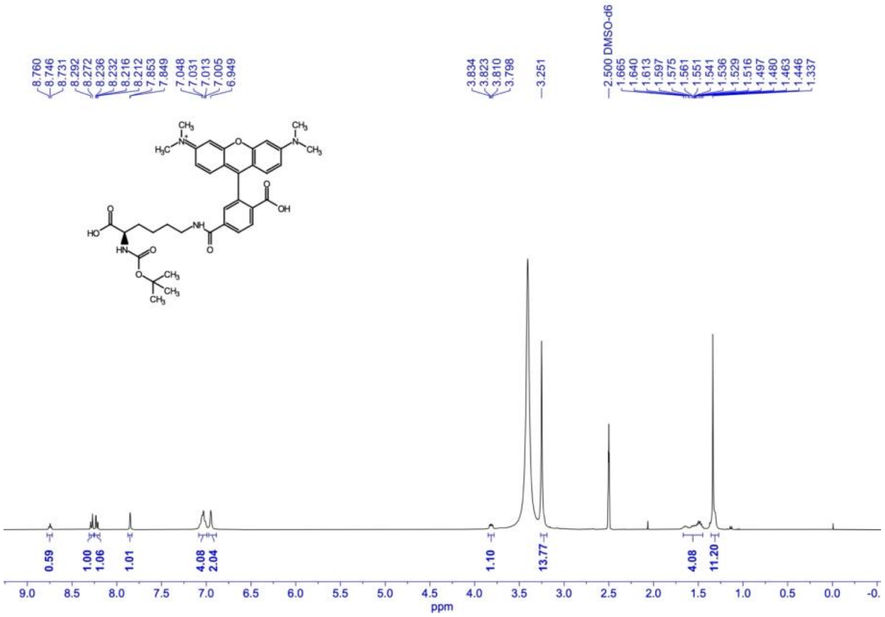

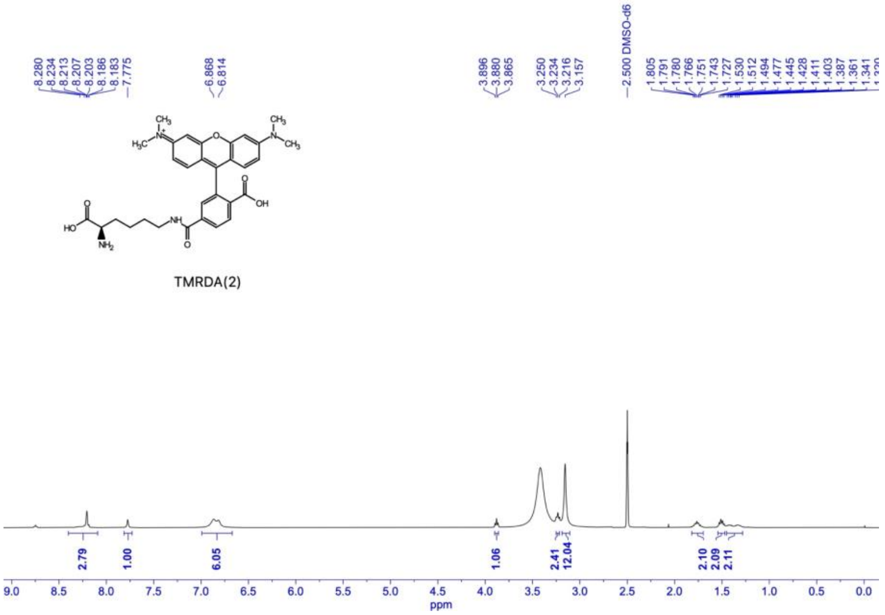

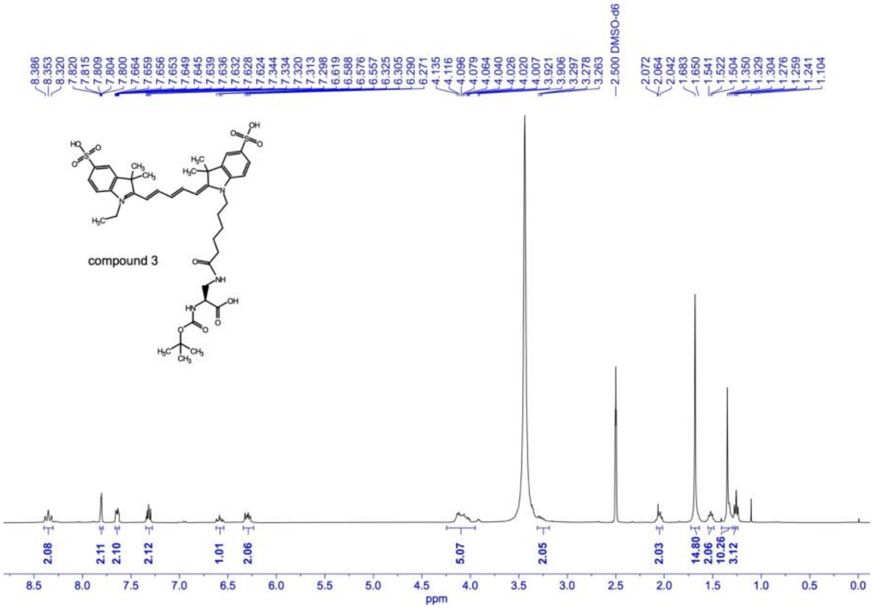
compound 3
2.500 DMSO-d6
ppm

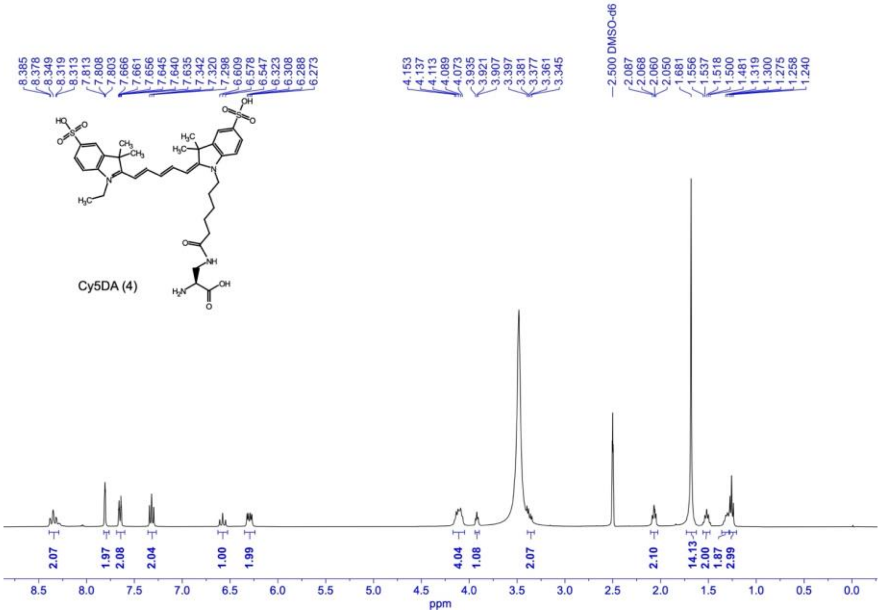
Cy5DA (4)
2.500 DMSO-d6
ppm

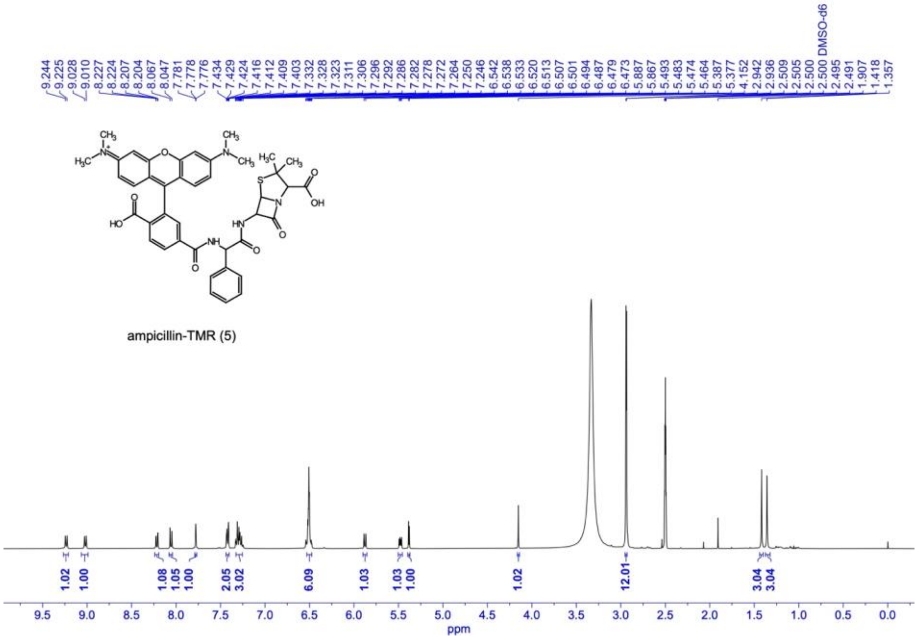


ampicillin-TMR (5)

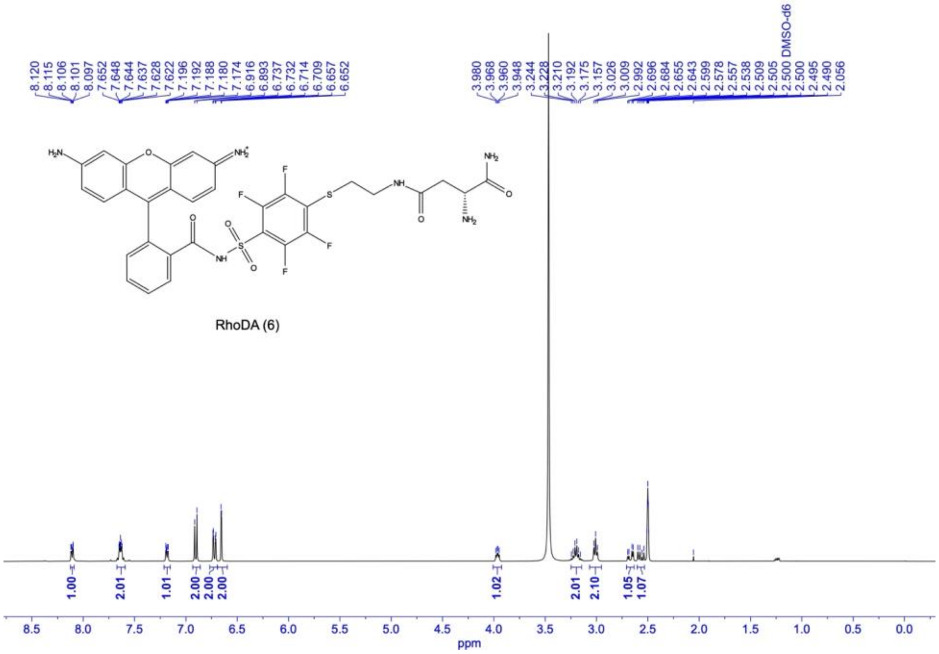


RhoDA (6)
